# Limited neutral and adaptive genomic divergence suggests *Acropora cervicornis* can be managed as a single conservation unit across its range

**DOI:** 10.64898/2026.08.26.747420

**Authors:** Paige J. Duffin, Maria Ruggeri, Trinity Conn, Iliana B. Baums, Macarena Blanco-Pimentel, Pol Bosch, Lisa Carne, Nichole Danser, Phanor Montoya Maya, Megan Morikawa, Erinn M. Muller, R. Scott Winters, Andrew C. Baker, Ross Cunning, Craig Dahlgren, John E. Parkinson, Carly D. Kenkel

## Abstract

Genomic signatures can provide key insight into the evolutionary history and remaining adaptive potential of threatened populations. As demographic decline erodes both diversity and the processes maintaining it, understanding how remaining variation is distributed becomes increasingly important for conserving species like the staghorn coral, *Acropora cervicornis*, a foundational but critically endangered Caribbean reef-builder. We analyzed 46 high-coverage *A. cervicornis* genomes from 10 locations across the tropical western Atlantic to evaluate neutral and adaptive structure, genomic diversity, demographic history, inbreeding, and connectivity, and generated a regional haplotype reference panel for future genomic monitoring. Genome-wide analyses recovered recurring regional substructure, but differentiation was modest and partly explained by isolation-by-distance and spatial variation in effective migration. Subpopulations had similar levels of genomic diversity, shared demographic history, and limited evidence of local adaptation. These patterns support interpreting sampled Caribbean populations as a single evolutionarily significant unit (ESU) containing multiple regional management units (MUs), rather than as deeply divergent evolutionary lineages. Despite substantial retained variation and low current inbreeding, estimated contemporary effective population size was small, suggesting an increased vulnerability to the effects of drift as demographic collapse continues, especially if structure is reinforced by isolated management. Together, our findings emphasize the urgent need for interventions that preserve and enhance genomic diversity, including risk-managed assisted gene flow. Supported by the haplotype reference panel developed here, these strategies will require coordinated efforts across regional entities to conserve and restore *A. cervicornis* as a jointly managed, single ESU.

## Introduction

The accelerating decline of global biodiversity due to anthropogenic change has made characterizing and preserving the evolutionary potential of threatened populations both a central goal and a central challenge in conservation biology. As population size, habitat continuity, and natural recruitment decline, species lose the genetic variation required to maintain fitness and respond adaptively to environmental change (Reed & Frankham 2003; DeWoody et al. 2021; Kardos et al. 2021). Consequently, conservation programs increasingly consider intervention strategies that actively shape genetic composition in focal species, including translocation, assisted reproduction, ex situ propagation, genetic rescue, and assisted gene flow, or AGF (e.g., Weeks et al. 2017; Browne et al. 2019; Werden et al. 2020; Rivers et al. 2020; Novak et al. 2021; Wylie et al. 2025). These strategies can restore connectivity and increase standing genetic variation, but ideally, they also consider which variation is preserved, propagated, mixed, and/or moved. Therefore, a key priority is to determine how remaining genetic diversity is distributed across declining populations, and how that structure should guide intervention actions (Frankham et al. 2011; Aitken & Whitlock 2013; Frankham & Admin 2017).

Genetic population structure arises when evolutionary and demographic processes produce differences in allele frequencies among groups of individuals and can provide a powerful basis for informing conservation boundaries. However, inferred genetic clusters are constrained by sampling design and analytical resolution, contributing to broader challenges in translating genetic structure into management units (Pritchard et al. 2000; Frantz et al. 2009; Lawson et al. 2018). Genetic structure patterns are often treated as if they inherently define conservation or management units, disproportionately favoring recommendations for separate management to preserve perceived uniqueness (Ralls et al. 2018; Bell et al. 2019; Liddell et al. 2021). The decision to maintain populations separately implies that the risks of mixing, such as outbreeding depression, outweigh those of continued isolation, including inbreeding and loss of genetic diversity; however, growing evidence suggests that research and management groups often fail to evaluate these competing risks explicitly (Cook & Sgrò 2019; Liddell et al. 2021). When inappropriately applied to small, threatened populations, isolated management can further reduce effective population size (*N_e_*), intensify genetic drift, and accelerate the loss of genetic diversity (Weeks et al. 2016), with direct consequences for fitness and future adaptive potential (Garner et al. 2005; Hoffmann & Willi 2008; Ørsted et al. 2019).

Well-informed conservation management therefore requires characterizing not only how genetic variation is distributed among populations, but also how much diversity remains and how vulnerable it is to loss. Genetic assessments can evaluate complementary dimensions of these risks, ideally with genome-wide datasets capable of distinguishing neutral from adaptive variation. For example, nucleotide diversity (π) measures standing variation, runs of homozygosity (ROH) provide insight into inbreeding, and outlier scans help assess whether differentiated variation has potential adaptive significance. Effective population size can further inform conservation benchmarks for avoiding contemporary inbreeding depression (current *N_e_*) and maintaining long-term adaptive potential (historical *N_e_*). Together, these approaches complement population structure analyses and help distinguish, for example, long-term evolutionary divergence from subpopulation structure within a broadly connected metapopulation, which is especially useful in studying marine species with high dispersal potential (Pritchard et al. 2000; Gagnaire et al. 2015; Perez et al. 2018; Schmidt et al. 2024).

Coral restoration has become one of the clearest settings in which conservation action stands to benefit from population genomic insight. In response to declining reef health, early restoration emphasized fragmentation and outplanting to restore biomass (Highsmith 1982), but it is now recognized that successful restoration also depends on genetic diversity to create resilient reefs (Baums 2008; Schopmeyer et al. 2012). Many programs track host genets (identified via their multilocus genotypes), maintain diverse lineages in nurseries, and monitor outplant performance (Lirman & Schopmeyer 2016), yet uncertainty remains about which and how many genets to preserve and propagate. As demographic collapse erodes diversity and the processes maintaining it, managers increasingly consider genetic and reproductive interventions to enhance genomic diversity (National Academies of Sciences & Medicine 2019; CRC Genetics Working Group 2025). However, effective implementation relies on genomic assessments of how diversity is distributed, how populations are connected, and where translocation is likely to help rather than harm recovery (Baums et al. 2019; Muller et al. 2026).

The staghorn coral, *Acropora cervicornis,* sits at the center of this conservation challenge as a foundational Caribbean reef-builder and one of the region’s most intensively restored species due to its critically endangered status and easy propagation. White Band Disease caused over 80% range-wide mortality in the 1970s and 1980s (Gladfelter 1982; Jackson 2014), followed by further losses from mass bleaching, hurricanes, and eutrophication (Knowlton et al. 1981; Cramer et al. 2020; Manzello et al. 2025). Natural recovery has been limited by reduced population density (Cramer et al. 2020) and little to no natural recruitment of sexually produced offspring (Hughes & Tanner 2000). In some regions, including Florida’s Coral Reef, these pressures have pushed *A. cervicornis* toward functional extinction (Manzello et al. 2025), making effective genetic management increasingly urgent.

Previous genetic studies have provided important insight into staghorn coral and related Caribbean acroporids, but differences in sampling and methodology leave range-wide management implications uncertain. Broad-scale Caribbean studies have repeatedly identified regional genetic structure in both *A. cervicornis* (Vollmer & Palumbi 2007; García-Urueña et al. 2022) and *A. palmata* (Baums et al. 2005, 2026), dividing populations into roughly northern, eastern, southern, and western regions. However, prior *A. cervicornis* analyses relied on reduced marker sets (at most 6.2k SNPs, García-Urueña et al. 2022), limiting power to critically evaluate population structure against complementary evidence of evolutionary independence, magnitude of differentiation, migration, demographic history, inbreeding, local adaptation, and ecological context. Additional insight into historical connectivity and genomic health is therefore needed to evaluate the future resilience of staghorn coral populations.

Here, we analyzed 46 high-coverage *A. cervicornis* genomes from 10 sampling locations spanning much of the species’ Caribbean range to test whether regional differences are strong enough to justify continued management of populations as discrete regional units. We estimated pairwise differentiation, isolation-by-distance, and effective migration to describe connectivity; compared nucleotide diversity, private allelic variation, demographic history, and ROH to assess genomic erosion; scanned for range-wide and regional selection; evaluated symbiont composition; and developed a haplotype reference panel for low-coverage imputation and future genomic monitoring. Together, these analyses inform risk-managed restoration, preservation, and monitoring for a critically endangered coral whose recovery now depends on active intervention.

## Methods

### Sample curation and whole-genome sequencing

In 2021, restoration practitioners and researchers provided samples from 3-5 genetically diverse *Acropora cervicornis* per location from Florida, Belize, Mexico, the Dominican Republic, Jamaica, Curaçao, and Aruba. Wild samples were collected from Dry Tortugas National Park in 2023 (permit DRTO-2023-SCI-0008), and Panama whole-genome data were downloaded from NCBI BioProject PRJNA950067 (Vollmer et al. 2023). Together, samples represented 10 locations spanning much of the *A. cervicornis* range (Figure 1a).

**Figure 1.**
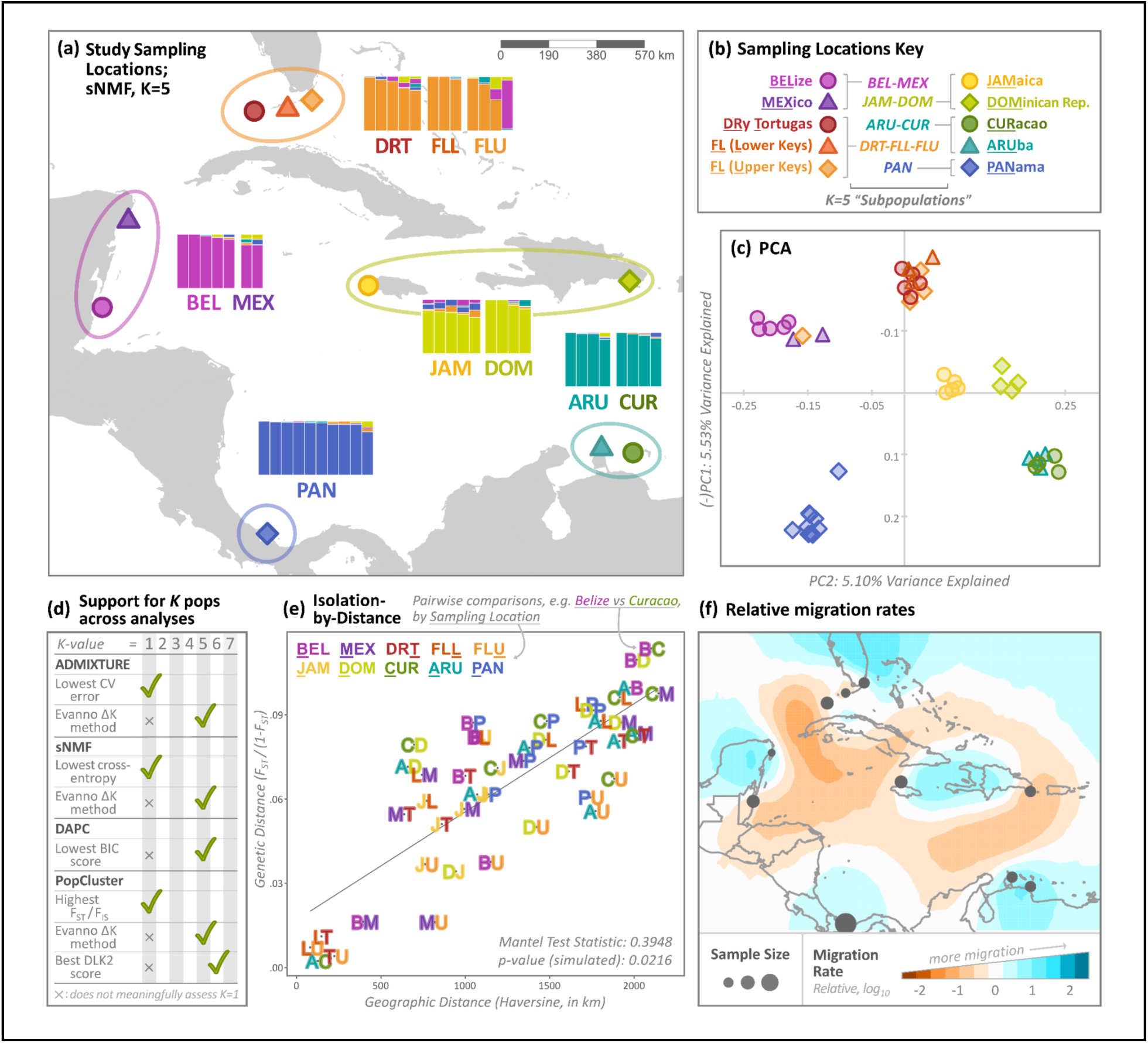
*A. cervicornis* population structure at neutral loci (n=177,516). (a) Study sampling locations (n=10) and sNMF results for K=5. Map to scale. Ellipses surrounding sampling location points represent identified “subpopulations” (K=5 post hoc groupings). Bar plots depict sNMF-derived ancestry coefficients by sample, grouped by sampling location. (b) Sampling location key. Underlined letters highlight nomenclature for site abbreviations. Italicized labels denote nomenclature for subpopulations. (c) PCA results colored by sampling location. PC1 was sign-inverted and plotted on the y-axis, with PC2 plotted on the x-axis, to show that major axes of genetic variation broadly reflect Caribbean geography. (d) Summary of K-selection results across population structure analyses: ADMIXTURE (Alexander et al. 2009), sparse Non-Negative Matrix Factorization (sNMF; Frichot et al. 2014), Discriminant Analysis of Principle Components (DAPC; Jombart et al. 2010), and PopCluster (Wang 2025). Green check marks indicate the best-supported K for each approach, whereas “x” symbols indicate cases where K=1 could not be evaluated due to limitations of the inference method or K-selection criterion (Appendix S7.1). (e) Isolation-by-distance analysis among sampling locations. Pairwise genetic distance, calculated as linearized *F_ST_* [*F_ST_*/(1 − *F_ST_*)], is plotted against Haversine geographic distance between sampling locations. Points represent pairwise comparisons between sampling locations, with labels indicating the two locations being compared using one-letter abbreviations. (f) Estimated effective migration surfaces showing relative migration rates between sampling locations compared to an isolation-by-distance model assuming a deme size of 500. Colors indicate slower (orange) or faster (blue) migration rates on a log_10_ scale. Sampling locations are indicated by dark gray dots, whose size corresponds to the number of individuals analyzed.

For newly acquired samples, genomic DNA was extracted by physical and enzymatic lysis and alcohol precipitation (Wilson et al. 2002), then purified and size-selected with Zymo Genomic DNA Clean & Concentrator kits. After quality assessment with gel electrophoresis, high-quality DNA was sent to Admera Health Biopharma for paired-end library preparation (Kapa Hyper Prep minimal PCR kit) and sequencing (Illumina NovaSeq X Plus, 150 bp pairedend reads).

### Genotyping and filtering

Bioinformatic analyses were performed on the USC Center for Advanced Research Computing cluster with detailed methods in Appendix S1. Newly generated raw reads (NCBI BioProject PRJNA1094960) were trimmed for adapters, low-quality bases, and 5′ bias. Because Panama samples were sequenced independently, initial processing followed (Vollmer et al. 2023) (Appendix S1.2). All reads were then quality-filtered and mapped to combined *A. cervicornis* host (GCA_964034985.1) and *Symbiodinium ‘fitti’* symbiont (Reich et al. 2021) genomes using BWA-MEM (Li 2013), then PCR duplicates were marked with Picard (“Picard” n.d.).

Sample-specific genotypes were called using bcftools v1.19 mpileup and multiallelic calling functions (Li 2011; Danecek et al. 2021), then merged across samples. Variant and invariant sites were filtered for depth, genotype quality, mapping quality, and missingness. SNPs near indels, showing strand bias, or deviating from Hardy–Weinberg expectations after population-structure correction with RUTH (Kwong et al. 2021) were also removed. Putative clonal Panama samples were identified using KING kinship coefficients in plink2 (Gaunt et al. 2007; Chang et al. 2015); 10 unique Panama genets were retained. One Curaçao sample (CU21E_1049) was removed from downstream analyses due to evidence of contamination (clear outlier in a preliminary PCA; *F_IS_* = -0.32) (Table S1.2).

### Neutral patterns of population structure

To evaluate population structure, we generated a neutral SNP panel to minimize the influence of selection and linkage disequilibrium, starting with the filtered file described above and retaining only biallelic SNPs with allele count >3, MAF ≥0.05, and missingness <0.1; we removed putative outliers detected through PCAdapt (Luu et al. 2017) and *F_ST_*-based OutFLANK (Whitlock & Lotterhos 2015) and LD-pruned remaining SNPs in plink2, yielding a panel of 177,516 variants (Appendix S1.3; Table S1.4).

We evaluated population structure using complementary ordination and clustering approaches (Appendix S2). PCA was performed with PCAdapt and DAPC with adegenet (Jombart et al. 2010). Ancestry coefficients were estimated with ADMIXTURE (Alexander et al. 2009), sNMF in LEA (Frichot et al. 2014; Frichot & François 2015), and PopCluster (Wang 2025). Models were evaluated across K=1–10 (though some methods do not evaluate K=1, see Figure 1d) using cross-validation error, cross-entropy, Bayesian information criterion, CLUMPAK (Kopelman et al. 2015), ΔK (Evanno et al. 2005), and PopCluster-specific metrics. Support for five post hoc clusters informed downstream analyses, hereafter termed “subpopulations”: BEL-MEX, DRT-FLL-FLU, JAM-DOM, ARU-CUR, and PAN.

Pairwise *F_ST_* was calculated from the pruned neutral and the full SNP panel (n= 2,527,725 SNPs) to evaluate differentiation among sampling locations and regional subpopulations (Figure 1a,b). Sampling-location comparisons were downsampled to at most four samples per site to reduce unequal representation (n=37); all were retained for subpopulation comparisons (n=46). Weighted *F_ST_* and confidence intervals were estimated using StAMPP (Weir & Cockerham 1984; Pembleton et al. 2013).

Isolation-by-distance was tested by Mantel correlation (100k permutations) between pairwise *F_ST_*/(1−*F_ST_*) and geographic distance (haversine method) in ade4 (Dray & Dufour 2007). To further evaluate spatial variation in connectivity among sampling locations, effective migration surfaces were modeled using EEMS (Petkova et al. 2016) across multiple deme-size settings, with convergence and migration surfaces visualized in rEEMSplots.

Full methods are reported in Appendix S2. One Upper Florida Keys sample (CRF_Acer-099) was excluded from subsequent post hoc analyses (genomic diversity and N_e_ estimates) due to complex admixture among three subpopulations. See Table S1.2 for sample lists by analysis.

### Current and historical effective population size (N_e_)

Recent *N_e_* and 90% confidence intervals were estimated from linkage disequilibrium of filtered SNPs using CurrentNe2 (Santiago et al. 2025), assuming an average recombination rate of 3.324 for major scaffolded chromosomes (Locatelli et al. 2024). Sequentially Markovian Coalescent (SMC) models were used to model historical changes in *N_e_* (500-1M years ago) using SMC++ (Terhorst et al. 2017). Filtered SNPs were converted into smc format, masking uncalled, filtered, and repetitive regions. The reference sample (distinguished lineage) was varied across chromosome files to generate replicate datasets for model estimation. *N_e_* was run using 3000 thinning, 50 EM iterations, 40 knots, a mutation rate of 4e^-9^ bp/year and a 5-year generation time (Matz et al. 2018; Fuller et al. 2020) (Appendix S3). CurrentNe2 and SMC++ were run separately for each subpopulation, with SMC++ also run on all samples combined (the “metapopulation”).

### Genomic diversity statistics

Both variant and invariant loci were retained for genomic diversity statistics. To account for bias due to sample size, eight individuals were randomly selected from each subpopulation (K=5). Pixy v2.0.0 (Korunes & Samuk 2021) was used to calculate nucleotide diversity (π), Watterson’s theta, and Tajima’s D in 100kb windows. Windows with at least 30kb callable loci were retained and used to calculate global (genome-wide) diversity statistics for each subpopulation. Private alleles were quantified using bcftools and filtered to include only alleles with ≥10x cumulative depth across various allele frequencies (0.1, 0.25, 0.5, 0.75, 1). Genome-wide heterozygosity was estimated including both variant and invariant loci using vcftools. Following Weeks et al. (2016), we additionally tested whether genetic diversity covaried with population-specific differentiation by regressing expected heterozygosity (*H_e_*) against population-specific *F_ST_* at both sampling-location and K=5 subpopulation scales (Appendix S3.2).

ROH were identified with GARLIC (Szpiech et al. 2017) and further analyzed in R. To focus on ROH likely to reflect more recent inbreeding, we removed very short ROH and estimated the expected time to common ancestry for ROH of different lengths using recombination maps adapted from (Locatelli et al. 2024) to the *A. cervicornis* genome assembly used in this study. We then calculated *ID_risk_* following Kyriazis et al. (2025), using the fraction of the genome contained in functionally long ROH and heterozygosity outside ROH regions (Appendix S3.3).

### Adaptive patterns

We evaluated putative signatures of selection at two scales. First, we treated all samples as a single metapopulation (K=1) to identify genomic regions with evidence of selection shared across the Caribbean. Second, we used subpopulation assignments to identify outlier loci or genomic regions diverging between five regional clusters, indicative of local adaptation (Appendix S4).

For the metapopulation scan, π and Tajima’s D were recalculated in 10 kb windows using Pixy. Candidate windows were defined as those in the extreme tails of the genome-wide nucleotide diversity distribution and showing Tajima’s D values consistent with balancing or purifying selection. Outlier windows were further examined at finer resolution, annotated for candidate genes, and visually assessed to determine whether genotype clustering reflected sampling location or subpopulation assignment.

We used SNP-based and window-based approaches to test for divergent selection among the five identified subpopulations. SNP-based outlier scans were conducted using PCAdapt (Privé et al. 2020), BayeScan (Duforet-Frebourg et al. 2014), and OutFLANK (Whitlock & Lotterhos 2015). Window-based scans were performed by calculating pairwise *F_ST_* and (sub)population-specific π in 10 kb windows for all pairwise contrasts among subpopulations. Candidate windows were identified as those with elevated *F_ST_* and extreme log_2_-transformed π ratios, then visualized as meadow plots (Duffin et al. in press). Contiguous outlier windows were merged into candidate peaks, and peak-length distributions were visualized for comparison across subpopulation pairs as in Duffin et al. (in press) (Appendix S4.1).

### Symbiont identification

Symbiont genera were screened by qPCR following Cunning & Baker (2013) and Palacio-Castro et al. (2021) using probe and primer sets specific to *Symbiodinium*, *Cladocopium*, and *Durusdinium* (Appendix S5.1), confirming *A. cervicornis’* predominant association with *Symbiodinium “fitti”* (Reich et al. 2021). Reads from high coverage samples mapping to the *S. “fitti”* genome were then parsed for ITS2 and population-level analyses (Appendix S5.2). Community composition at the ITS2, allele-frequency, and dominant-haplotype level were compared across sampling locations using PERMANOVAs from the R package vegan (Dixon 2003).

### Haplotype panel construction and validation

To generate a phased *A. cervicornis* haplotype reference panel, we used a two-step workflow combining read-backed and population-based phasing (Figure 3a). SNPs were filtered to retain high confidence biallelic loci with no missing genotypes and allele counts >3. Read-backed phasing was performed for each individual using WhatsHap v2.4 (Martin et al. 2016) and phased VCFs were merged. Before population-based phasing, strongly admixed or ambiguously assigned samples were removed to limit phasing artifacts, leaving 44 panel samples. SNPs deviating from Hardy–Weinberg expectations after accounting for population structure were removed using RUTH (Kwong et al. 2021), and the final filtered dataset was phased by scaffold with Beagle v5.5 using scaffold-specific genetic maps (Browning & Browning 2007). Scaffold outputs were concatenated into the final *A. cervicornis* reference panel.

We validated imputation accuracy using five *A. cervicornis* samples independently sequenced at both high and low coverage (Figure 3b). High-coverage genotypes served as the truth set, while low-coverage genotypes were imputed using leave-one-out reference panels. Low-coverage genotype likelihoods were first imputed with Beagle v4.1 (Browning & Browning 2007) and filtered by genotype probability to remove low-confidence calls, then imputed again with Beagle v5.5 (Browning et al. 2021) using the phased reference panel. Final genotypes were filtered using Beagle’s dosage R² metric (DR2≥0.99). Accuracy was evaluated against high-coverage genotypes using raw, genotype-probability-filtered, and newly imputed calls, and summarized by overall and genotype-class-specific concordance and global dosage correlation. Full methods are provided in Appendix S6.

## Results

Whole genome sequencing (WGS) of 36 new *Acropora cervicornis* sourced from 9 Caribbean locations yielded an average of 57.6M paired reads per sample. Of these, 48.6M paired reads and an additional 5.3M unpaired reads per sample passed quality filters, yielding mean sequencing coverage of 37.51x (± 2.38x, range: 32.78x to 41.99x) across the main staghorn coral chromosomes (Table S1.1). Publicly available Panama WGS samples (n=10; (Vollmer et al. 2023) had slightly higher but more variable coverage (49.14x ± 15.60x, range: 23.22x to 71.14x). Mapping all 46 samples to the *A. cervicornis* genome recovered 267M high-confidence sites, of which 3.8M were variable (1.4% of the genome).

### Broad connectivity with substructure reflecting Caribbean geography

Across 177,516 neutral, unlinked SNPs, principal component analysis (PCA) revealed a pattern that broadly mirrored Caribbean geography (Figure 1, Appendix S7). PC1 separated samples primarily along a north-south gradient, while PC2 resolved east-west structure (Figure 1c). This ordination produced five visually distinct regional groupings: Aruba/Curaçao (ARU/CUR); Dominican Republic/Jamaica (DOM/JAM); Panama (PAN); Dry Tortugas/Lower/Upper Florida Keys (DRT/FLL/FLU), and Belize/Mexico (BEL/MEX), with one FLU individual grouping with BEL/MEX (Figure 1c).

When comparing support for the number of genetic clusters across population structure methods and K-selection criteria (summarized in Figure 1d), every criterion capable of evaluating K=1 favored K=1, consistent with substantial shared ancestry and limited discrete subdivision across the sampled range. Among criteria restricted to K≥2, support was strongest for K=5 (ADMIXTURE, sNMF, DAPC, PopCluster), with one line of alternative support for K=6 (PopCluster) (Appendix S7.1). We therefore interpret K=1 as the dominant ancestry signal but use K=5 to test whether regional structure reflects longer-term evolutionary divergence. In downstream analyses and interpretations, we refer to these post hoc K=5 clusters as “subpopulations” and the combined K=1 sample set as the “metapopulation.”

Across the supported K≥2 ancestry-coefficient analyses, nearly all samples showed dominant assignment to clusters geographically consistent with their sampling origin (Appendix S7). The clearest exception was the FLU sample which repeatedly grouped with BEL/MEX across approaches (Figure 1a; Table S7.1). JAM also showed elevated mixed ancestry relative to DOM, despite grouping together at K=5 (Figure 1a; Appendix S7.2, Appendix S7.3).

Pairwise *F_ST_* values calculated from the pruned neutral SNP panel were generally low across both sampling locations (average: 0.061 ± 0.026) and subpopulations (average: 0.066 ± 0.012, Appendix S8). Pairwise *F_ST_* values were lowest among repeatedly clustered locations, including DRT/FLL/FLU, ARU/CUR, and BEL/MEX, and were generally higher among more distant regional comparisons and comparisons involving PAN (Figure S8.1).

Isolation-by-distance and effective migration analyses further indicated that geographic distance likely contributes to regional structure (Appendix S8). Pairwise genetic distance increased significantly with geographic distance among sampling locations, supporting IBD (Mantel statistic=0.395, simulated p=0.022; Figure 1e). Effective migration was higher than expected under an IBD model among neighboring coastal regions, including DRT/FLL/FLU, BEL/MEX, JAM/DOM, and ARU/CUR, but lower than expected across broader Caribbean Sea comparisons, and coastal regions neighboring PAN (Figure 1f).

### Consistent Symbiont Composition

All corals hosted Symbiodiniaceae from the genus *Symbiodinium* (Appendix S9), with the majority of ITS2 database hits matching type A3g and A3l variants (Figure S9.1), consistent with the species *Symbiodinium ‘fitti’* (*nomen nudum*). Symbiont populations did not differ by sampling location when considering ITS2 variants or finer-scale genome-wide metrics such as allele frequencies and dominant haplotypes (PERMANOVA p>0.05, Figure S9.2).

### Patterns of genomic diversity and demographic history

Demographic history trajectories and current *N_e_* estimates were similar across subpopulations, indicating similar past and present subpopulation demographics. Global Tajima’s D was slightly negative, but close to neutral, across all subpopulations (-0.339 to - 0.147, Figure 2a). Genome-wide heterozygosity ranged from 0.224-0.318%. Demographic history models of *N_e_* over time indicated a historical population expansion after the last glacial maximum (Figure 2f), followed by relatively stable population sizes in the recent past (500-5,000 years ago). However, nucleotide diversity (π) and current *N_e_* estimates suggest a more recent population collapse. π was 0.219-0.240% (Figure 2b), which corresponds to a cumulative, long-term *N_e_* of 137k-150k individuals assuming the same mutation rate as demographic history models (Figure 2f) while LD-based methods estimate that current *N_e_* is 72-279 within subpopulations (Figure 2g).

**Figure 2:**
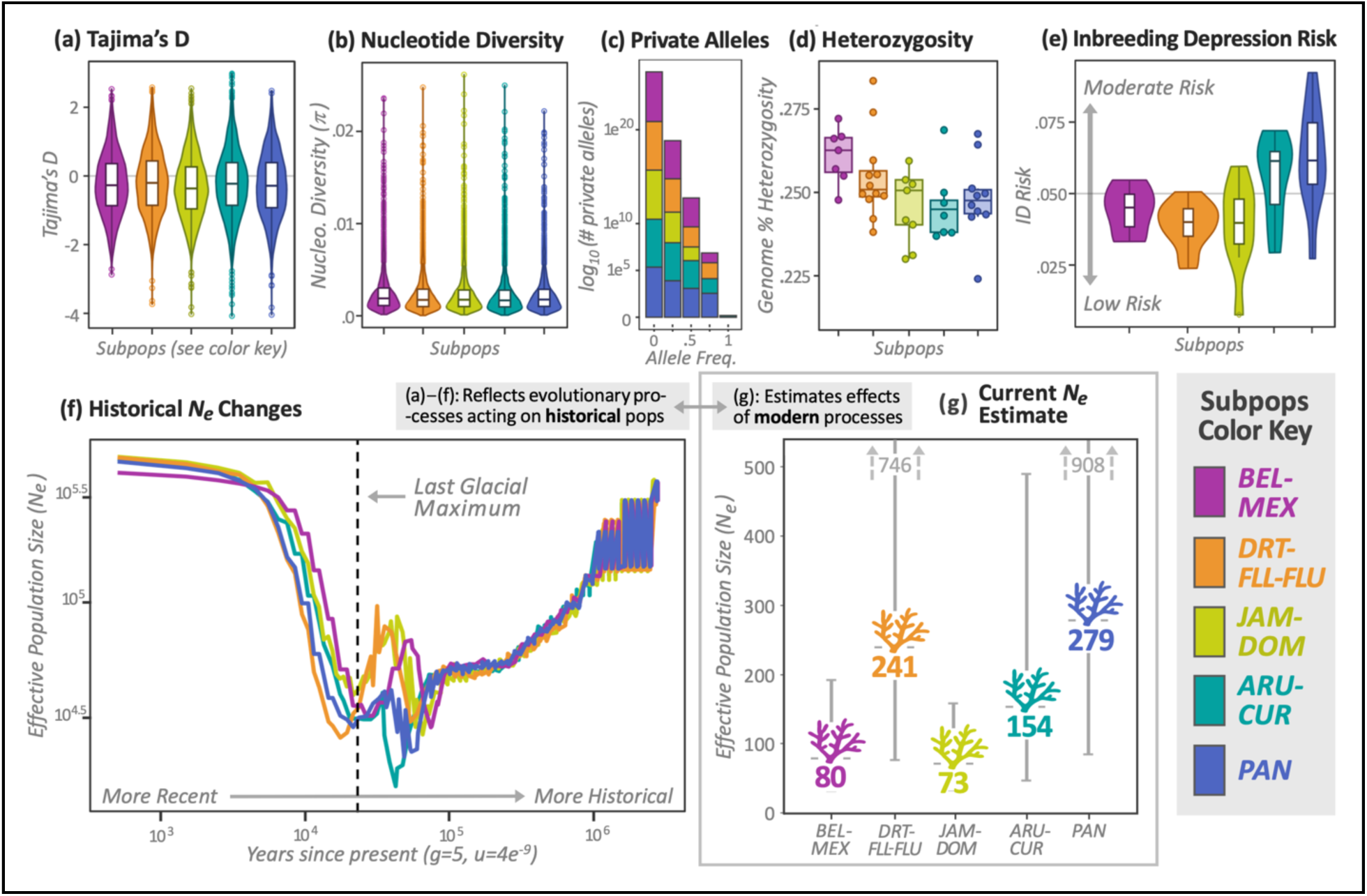
Genomic diversity and demographic history across subpopulations. All genomic diversity and demographic history estimates were analytically grouped and visually colored by subpopulation (see bottom right ‘Subpop[ulation]s Color Key’). Panels (a) and (b) show the distribution of Tajima’s D and π (nucleotide diversity per bp), respectively, calculated in 100kb windows across the genome. Individual points reflect 100kb outlier windows. (c) The log10-transformed number of alleles private to each subpopulation across different allele frequency thresholds. (d) Genome-wide heterozygosity, represented as a percentage. Individual points reflect individual samples within each subpopulation. (e) Inbreeding depression risk (ID Risk) where values above or below the dashed line are respectively considered at moderate or low risk for inbreeding depression. (f) Historical effective population size (N_e_) modeled from allele coalescence over time (SMC++), assuming a generation time of 5 years and mutation rate of 4e^-9^. The dashed line indicates the last glacial maximum (LGM) when sea level was at a historical low. (g) Current *N_e_* estimates (1-100 generations) based on linkage disequilibrium between loci assuming a recombination rate of 3.324. Error bars represent 90% confidence intervals.

Global genomic diversity statistics were also similar across subpopulations (Appendix S10). JAM-DOM had the lowest global Tajima’s D (-0.339), and DRT-FLL-FLU the highest, though still negative, value (-0.147); other subpopulations were intermediate. Nucleotide diversity was highest in BEL-MEX (0.00240) and lowest in ARU-CUR (0.00219), but otherwise similar across subpopulations. Furthermore, expected heterozygosity (*H*_e_) was strongly inversely related to population-specific *F_ST_* (R²=0.95, p=0.0047; Figure S10.2).

Subpopulations nevertheless contained unique variation. Of 3.8M SNPs analyzed, 22% were private alleles present in at least one genotype, averaging 4.5% unique alleles per subpopulation. When standardized across the 318-Mb genome, however, private alleles represented only 0.054% of possible variation. Their abundance also decreased exponentially with increasing allele-frequency thresholds, indicating that most were rare and only one was at fixation (Figure 2c).

ROH length and fractions of genomic coverage (FROH) did not differ significantly among subpopulations in either the >100-kb or >200-kb datasets (ANOVA, p > 0.05; Table S10.3), providing no evidence of unusually elevated inbreeding in any sampled region. Because ROH ≥ 228.9 kb were inferred to have arisen within approximately the past 25 generations, subsequent analyses focused on ROH >200 kb. Only 46 were identified across all samples, with mean FROH low and similar among subpopulations (range: 0.0201-0.0334). Mean heterozygosity outside ROH was similar across subpopulations but high (1.87–1.96 variants/kb) relative to other rapidly declining species (e.g., Robinson et al. 2021), although sourceable comparisons were limited to vertebrates, which generally have lower heterozygosity than invertebrates (Gayral et al. 2013). In *A. cervicornis*, this trend suggests substantial deleterious variation may remain masked and become exposed after several generations of breeding within small populations. Accordingly, mean *ID_risk_* scores ranged from 0.0388 to 0.0625, placing all subpopulations in the “Moderate Risk” category (Kyriazis et al. 2025) (Figure 2e). See Appendix S10 for details.

### Functional variation is not strongly partitioned among regional subpopulations

Because population structure analyses supported historically broad Caribbean-wide connectivity with some degree of spatial differentiation, we evaluated signatures of selection at two scales, testing for candidate genomic regions under selection across the full metapopulation (K=1), then asking whether divergent selection was evident among the five identified subpopulations (Appendix S11).

Six 10kb genomic regions were putatively under selection in the metapopulation (Appendix S11.1).Three showed low nucleotide diversity and negative Tajima’s D, consistent with purifying selection, and contained the candidate genes *WBP11* and *ABCA3* (Table S11.1). Three additional regions showed high nucleotide diversity and positive Tajima’s D, consistent with balancing selection, and contained *OAS3* and *TTC37* (Table S11.1). Because *OAS3* is associated with innate immune response pathways (Sadler & Williams 2008) and may be relevant to coral disease, we further evaluated variation across this gene. Sample-specific *OAS3* genotypes did not cluster by sampling location or subpopulation (Figure S11.2), suggesting that the balancing-selection signal was not driven by local adaptation.

Additionally, we found limited evidence for divergent selection among subpopulations. Of the three locus-based outlier methods, only PCAdapt identified candidate loci (n=196; Table S11.4). Because none overlapped BayeScan or OutFLANK results, these candidates were not prioritized for annotation or follow-up. Window-based meadow plots, which scan for putative selective sweeps using elevated *F_ST_* and shifts in π (Duffin et al. in press), visualized sparse candidate outlier windows across pairwise subpopulation-level contrasts (Figure S11.3). While a few modest candidate peaks were detected (Figure S11.4, Table S11.2), including a region driven by reduced nucleotide diversity in Panama and overlapping a putative *LYS1* annotation (Table S11.3), these signals were not consistently recovered across contrasts and did not closely track genome-wide *F_ST_* or geographic distance (Appendix S11).

### Community resource: Caribbean-wide haplotype reference panel

We generated a haplotype reference panel for *A. cervicornis* from 44 high-coverage genomes spanning 10 Caribbean sites (Figure 3a, Appendix S12). After quality filtering and two-step phasing, the panel contained 1,677,307 high-confidence SNPs. Mean sequencing coverage was 45.39 ± 10.77× across final reference panel SNPs (Figure 3a).

**Figure 3.**
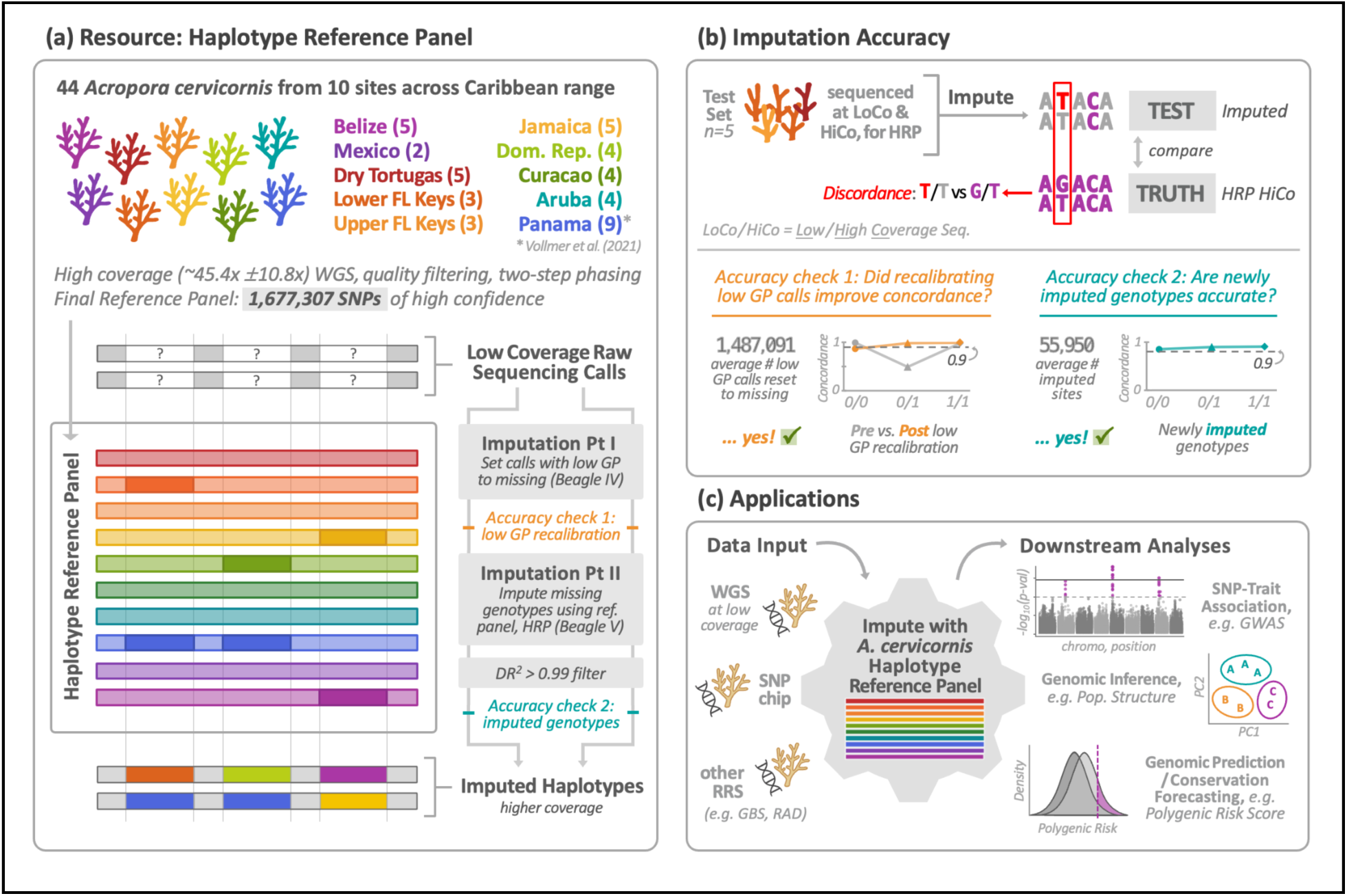
(a) Haplotype reference panel composition and imputation process. (b) Imputation accuracy of the reference panel. (c) Exemplar applications of *A. cervicornis* datasets after imputing with the haplotype reference panel resource. Acronyms: WGS = Whole-Genome Sequencing; SNP = Single Nucleotide Polymorphism; GP = Genotype Probability; HRP = Haplotype Reference Panel; DR^2^ = Dosage, R-squared; LoCo = Low Coverage (sequencing); HiCo = High Coverage (sequencing); RRS = Reduced Representation Sequencing; GBS = Genotyping-By-Sequencing; RAD = Restriction-Site Associated DNA (sequencing); GWAS = Genome-Wide Association Study.

We evaluated low-coverage genotype imputation in five samples sequenced at both high and low coverage (Figure 3b), using high-coverage genotypes as truth sets. Prior to correction, raw low-coverage calls averaged 85.60% concordance, but accuracy varied by genotype: 98.70% for 0/0, 98.00% for 1/1, and 48.74% for 0/1. Genotype-probability recalibration reduced retained calls from 2,699,811 to 1,212,720 but increased mean concordance to 91.54%, with especially strong improvement in heterozygotes (97.67%) (Figure 3b).

Newly imputed genotypes averaged 93.15% concordance across 55,950 sites per sample, with similar accuracy across genotype classes (92.46% - 94.58%). Global Pearson R² increased from 64.13% for raw calls to 87.77% after recalibration and 90.41% after imputation. Together, these results show that our imputation pipeline using the *A. cervicornis* haplotype reference panel both rectifies unreliable low-coverage genotype calls and accurately imputes missing genotypes, providing a genomic resource for downstream analyses (Figure 3c).

## Discussion

As coral populations continue to decline, it will be important to consider the spatial distribution of genetic variation to support successful conservation outcomes. Here, our analyses indicate that *A. cervicornis* genomes reflect historically high levels of gene flow across the sampled Caribbean. Effective population size, metrics of genomic diversity, demographic history, and symbiont composition were similar across regions, and scans for divergent selection did not reveal broad, repeated, or region-specific signatures of local adaptation. At the same time, significant genetic differentiation, isolation-by-distance, and spatial variation in effective migration indicate that demographic exchange is not uniform. Together, these patterns are most consistent with a single evolutionary significant unit (ESU) containing multiple regional management units (MUs): regional subpopulations show restricted demographic interchange but little evidence of long-term evolutionary independence (Table 1). Importantly, substantial genomic diversity persists despite recent demographic decline, even as small contemporary *N_e_* indicates increasing vulnerability to drift and inbreeding. We therefore urgently recommend conserving existing variation, maintaining and/or restoring genetic connectivity (including through AGF; Table 1), and tracking regional ancestry to prevent further genomic erosion. The haplotype reference panel developed here provides a scalable tool for supporting these efforts and monitoring their outcomes.

**Table 1.**
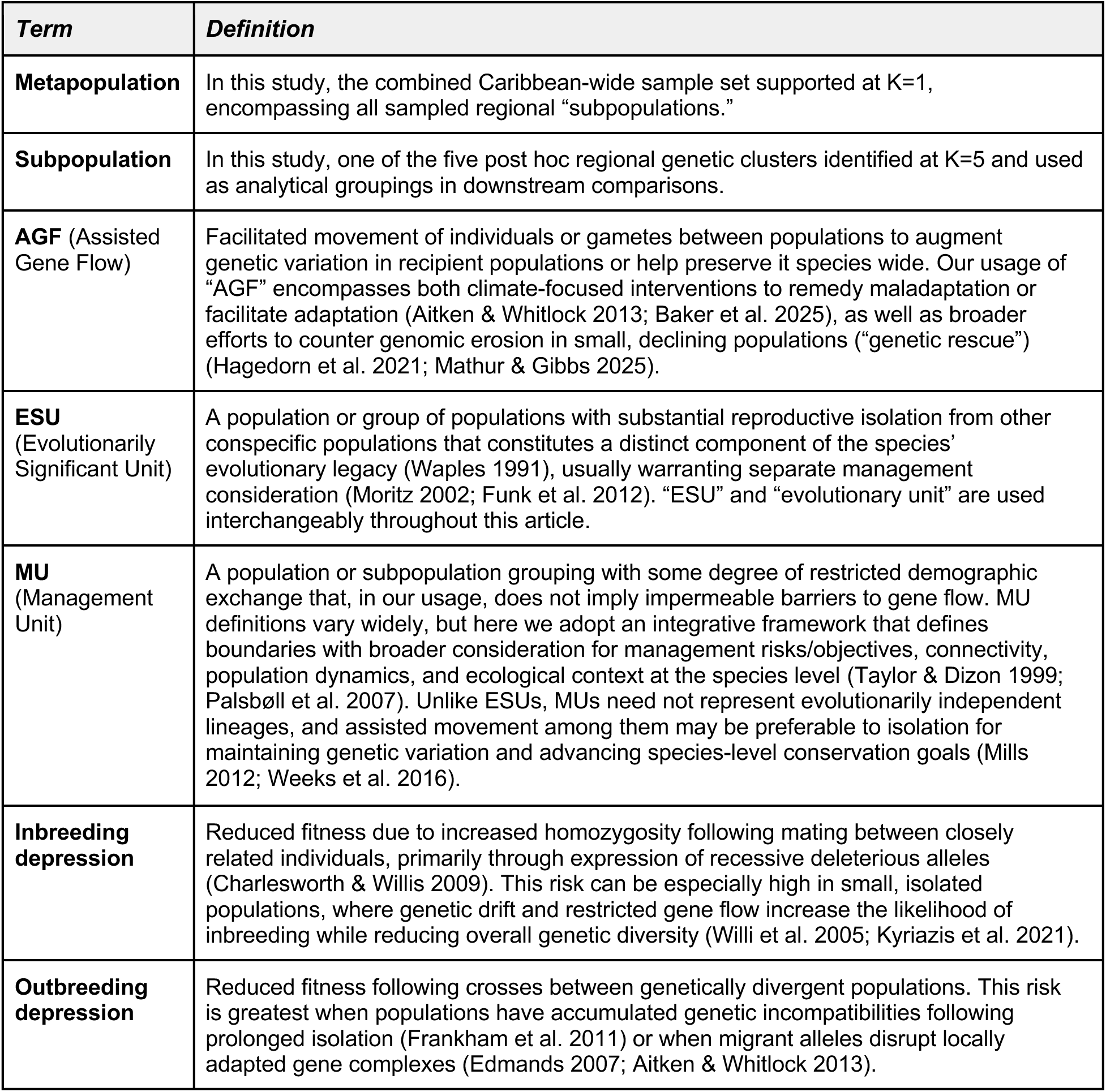
Definitions and study-specific usage of key population genetic and conservation terms used throughout this article.

### Reinforcing regional isolation risks accelerating genomic erosion

Our results support a distinction between evolutionary and demographic population structure that is important for conservation management. Viewed through an ESU-MU framework, the regional structure detected here provides evidence for multiple MUs within a shared Caribbean ESU rather than either complete panmixia or deeply divergent evolutionary units. The hierarchy of population structure we observed is consistent with this interpretation. Every clustering approach that could meaningfully evaluate K=1 identified it as the best solution, while the K=5 solution summarized recurring regional affinities, or “subpopulations,” among Belize/Mexico, Florida/Dry Tortugas, Jamaica/Dominican Republic, and Panama. However, patterns of genomic diversity as well as locus- and window-based outlier scans found limited evidence for repeated divergent selection among these subpopulations, suggesting that putatively selected variation was not consistently associated with regional ancestry strongly enough to generate clear local adaptation signals in our dataset.

Spatial analyses further suggest that regional structure reflects partial restrictions to gene flow rather than discrete barriers. Consistent with prior work in *A. cervicornis* (García-Urueña et al. 2022), differentiation increased with geographic distance, while effective migration estimates indicated that connectivity was also shaped by seascape features: neighboring coastal sites were more connected than expected under isolation-by-distance, whereas some open-water comparisons showed reduced connectivity. Currents, gyres, and passages may therefore act as partial filters to larval dispersal, such as potentially reduced exchange across the Mona Passage (Baums et al. 2005). Accordingly, repeated assignment of one Upper Florida Keys sample to the Belize/Mexico cluster suggests possible long-distance dispersal within the past generation, although we interpret this cautiously (Lawson et al. 2018). At minimum, it demonstrates that sampling location does not necessarily predict shared genomic ancestry, as some existing management strategies assume (Baker et al. 2025).

Our interpretation of regional subpopulation structure is generally consistent with previous population genetic studies of *A. cervicornis*, although estimates of the magnitude of differentiation vary. Earlier studies using mitochondrial and nuclear markers detected strong differentiation between regions separated by >500 km (Vollmer & Palumbi 2007), whereas SNP-based analyses suggest more moderate differentiation across similar distances (Drury et al. 2016; García-Urueña et al. 2022). These differences may partly reflect sampling design and marker resolution, as genome-scale datasets can refine estimates of population structure and alter conservation interpretations derived from reduced marker sets (Supple & Shapiro 2018). Our relatively low genomic differentiation is more similar to a recent WGS comparison between Florida and Panama (*F_ST_* =0.04; Vollmer et al. 2023). Together, these results help reframe earlier evidence of restricted gene flow with a genome-wide view in which regional differentiation reflects reduced exchange rather than long-term separation.

The management significance of this distinction becomes clearer when regional structure is considered alongside temporal genomic signals. Contemporary genomic diversity is shaped by both historical and recent events, with different implications for population health and adaptive potential (Jamieson & Allendorf 2012). Long-term *N_e_* (>1,000 generations) reflects historical mutation-drift dynamics, while current *N_e_* (<100 generations) inferred from associations among loci, more directly captures recent demographic conditions and extinction risk (Hare et al. 2011). Our long-term *N_e_* estimates greatly exceed proposed benchmarks for buffering drift-driven loss of genomic diversity over broad timescales (*N_e_* > 500, Franklin 1980; Franklin et al. 2014, or *N_e_* >1,000, Frankham et al. 2014). In contrast, current *N_e_* estimates were much smaller, approaching and, in some subpopulations, falling below the more conservative minimum proposed as a threshold for avoiding extinction due to inbreeding depression (*N_e_* > 100, (Frankham et al. 2014; but see *N_e_* > 50 in Franklin 1980; Franklin et al. 2014). Although our values may be underestimated because of life history trait based-model violations, our low current *N_e_* estimates more closely reflect the ongoing demographic collapse of *A. cervicornis* (Cramer et al. 2020; Manzello et al. 2025) than metrics shaped by longer-term processes. Together, these contrasting signals suggest that sampled individuals retain substantial genomic diversity inherited from historically large and connected populations, but that diversity is increasingly vulnerable to loss under contemporary decline.

The persistence of this historical genomic signal is not unexpected given the relative recency of staghorn coral population decline. Nucleotide diversity and coalescence change slowly, and therefore do not yet resolve the approximately 80% census decline of *A. cervicornis* since the 1970s (Jackson 2014). Tajima’s D is more sensitive to rare-variant loss (van Oosterhout et al. 2026), but its cumulative signal means our near-neutral values may likewise reflect historical expansion followed by very recent contraction or somatic mutations (Vasquez Kuntz et al. 2022; Conn et al. 2025). Other genomic metrics similarly provide little evidence that severe erosion has already occurred: low-frequency private alleles and potential maintenance of functional variation in the immune gene *OAS3* via balancing selection suggest persistence of genomic variation, while ROH were relatively short and covered only 2.5% of the genome, yielding low to moderate *ID_risk_* scores and indicating that sampled individuals were not yet highly inbred (Kyriazis et al. 2025). Together, these patterns suggest that demographic collapse is occurring faster than its consequences are yet visible in measures of genomic diversity and inbreeding, potentially imposing a drift debt that will increasingly curtail adaptive capacity in future generations (Gargiulo et al. 2025).

That mismatch is especially concerning because ongoing habitat loss, reduced colony density, and limited sexual recruitment have likely weakened the ecological processes that historically maintained gene flow among reefs (Williams et al. 2008, 2020; Goergen et al. 2019; Baums et al. 2019; CRC Genetics Working Group 2025). Notably, our collections also predate the 2023 marine heatwave, which caused up to 90% mortality of *A. cervicornis* (DeBuysser et al. 2025; Le Gall et al. 2025; Manzello et al. 2025), likely pushing contemporary populations closer to conditions under which drift and inbreeding become increasingly consequential. Large historical *N_e_* and relatively high heterozygosity outside ROH regions may further indicate that remaining individuals carry substantial masked genetic load (Bertorelle et al. 2022), creating the potential for inbreeding depression as declining *N_e_* exposes deleterious variation and reduces fitness (Hedrick & Garcia-Dorado 2016).

Moreover, the significant negative relationship we observed between genetic diversity (*H_e_*) and differentiation (population-specific *F_ST_*) across subpopulations mirrors patterns in other threatened species with small, fragmented populations, where efforts to preserve drift-generated population ‘uniqueness’ may actually exacerbate the processes driving genomic erosion (Weeks et al. 2016). Together with small contemporary *N_e_*, this trend suggests that drift may become increasingly consequential in *A. cervicornis*, accelerating diversity loss while reducing the efficacy of purifying selection and increasing the probability that slightly deleterious variants become fixed over time (Grossen et al. 2020; Robinson et al. 2023). Although we do not demonstrate that differentiated variants are themselves deleterious and therefore cannot infer that genetic load has already increased within subpopulations, the conditions promoting such accumulation are increasingly present (Grossen & Ramakrishnan 2024). This interpretation makes a growing body of work identifying a bias in the field of conservation genetics particularly relevant: assessments that do not consider the full scope of genomic risks tend to overemphasize the importance of maintaining subspecies-level “purity,” disproportionately favoring practices of separate management beyond what the available evidence supports (Ralls et al. 2018; Bell et al. 2019; Liddell et al. 2021). In *A. cervicornis*, we therefore warn that continued conservation strategies maintaining or reinforcing regional isolation would likely compound emerging risks of genomic erosion by further reducing *N_e_* and restricting gene flow, exacerbating drift-driven loss and inbreeding depression. Notably, we also find little fixed or near-fixed differentiation, suggesting that this process has not yet produced deep divergence among subpopulations, thereby mitigating concern about outbreeding-depression risk following recommended translocation of contemporary corals (see Table 1 and discussion below).

### Conservation recommendations

Collectively, these spatial and temporal patterns suggest that, while present *A. cervicornis* samples retain a genomic legacy of historically favorable conditions, these benefits may only temporarily mask the consequences that lag behind demographic collapse. With this in mind, we emphasize that our characterization of subpopulations as MUs does not imply that their differentiation should be preserved through isolated management, as this threatens to compound the genomic risks associated with small contemporary *N_e_*, weakening connectivity, and ongoing demographic decline. Importantly, however, substantial standing variation, low inbreeding, and minor regional divergence indicate that the genomic consequences of recent decline remain limited, creating a time-sensitive window for intervention before severe genomic erosion occurs. The following recommendations therefore focus on preserving remaining variation while supporting the evolutionary processes needed to sustain it.

A top priority should be biobanking and cryobanking of wild populations and diverse lineages (Hagedorn et al. 2021). Because substantial genomic diversity remains, preserving that variation now can safeguard genotypes for future restoration efforts that may otherwise be lost as demographic decline continues. Living and cryopreserved collections should therefore seek broad representation across the species’ range rather than disproportionately preserving locations or lineages already well covered.

Whereas biobanking and cryobanking are widely accepted conservation tools, interventions that deliberately move genetic variation across regions through translocation and AGF remain more controversial (Vinton et al. 2025). Given the considerable historic genetic mixing throughout the sampled range, evidence of shared demographic histories, comparable genomic diversity, limited evidence for local adaptation, and little fixed or near-fixed differentiation, our results suggest that AGF is unlikely to generate strong outbreeding depression in this species (Table 1; Appendix S13). In one promising demonstration, experimental AGF crosses using cryopreserved sperm from geographically distant *A. palmata* samples produced viable, genetically novel offspring (Hagedorn et al. 2021) with no apparent performance deficits (Muller et al. 2025). Therefore, although AGF should be implemented within an appropriate risk-management framework, our findings imply that increasing gene flow poses less genomic risk than allowing drift and inbreeding to intensify under continued regional isolation (Appendix S13). Regulatory and permitting restrictions, however, remain major barriers to AGF research and implementation in Caribbean acroporids (reviewed in Baker et al. 2025).

Non-genomic risks also require consideration, and regional differences in symbiont communities could complicate long-distance translocations. All samples in our dataset hosted the same Symbiodiniaceae species; however, finer-scale symbiont composition was not resolved here, and known associations with other symbiont species warrant further study. Disease transmission was also not evaluated, but monitoring parental and offspring health during experimental crosses in closed, land-based systems could reduce disease risk while allowing remaining genetic uncertainties surrounding AGF to be evaluated empirically before intervention at larger scales.

Finally, the haplotype reference panel developed in this study provides an applied tool for putting these strategies into practice while tracking ancestry, avoiding over- or under-representation of lineages in nurseries or biobanking, and evaluating intervention outcomes. It can be integrated with expanding phenotype datasets, such as NOAA’s AcDC (*Acropora cervicornis* Data Coordination Hub; Kiel et al. 2025), to support association mapping and genomic prediction methods that help identify high-performing genotypes or crosses and subsequently balance their use with retention of overall genomic diversity.

Taken together, our findings shift the framing of AGF and related genetic interventions from artificial disruptions of isolated populations to managed extensions of the same processes that have historically shaped variation across the species’ range. Our recommendations for broad biobanking and cryopreservation, regulated translocations, and genetic monitoring together provide a framework for actively managing genomic variation while sufficient diversity remains to conserve and restore the critical Caribbean reef builder *A. cervicornis*.

## Acknowledgments

Florida coral were sampled under permits FKNMS-2019-012 and FKNMS-2021-172. Archived tissue samples from Belize and Curaçao were obtained from Penn State University collections. Collections and exports from the Dominican Republic, Jamaica, and Mexico complied with applicable research, export, and CITES requirements. In the DR, activities were additionally conducted under an agreement between the Ministry of Environment and Natural Resources and Iberostar, consistent with access- and benefit-sharing principles under the Convention on Biological Diversity and the Nagoya Protocol. Research in Mexico was conducted under a CINVESTAV permit and in Jamaica under authorizations from the National Environment and Planning Agency. Aruba samples were exported under CITES permit 1509.

This work was supported by NSF grants OCE-2023155 to ACB; OCE-2023187 to JEP; and OCE-2023705 to RC, CD, and CDK, and the Human Frontiers Science Program # RGP0042/2020 to I.B.B. Salary support was provided to PJD by the Mary G. Jameson Foundation and the Gordon and Betty Moore Foundation, MR by NSF PRFB #2508011, and TLC by the Walder Foundation. Additional support provided by the Paul M. Angell Family Foundation, the Paul G. Allen Foundation, the Dr. Scholl Foundation, and the Brunswick Foundation.

## Data availability

Raw sequencing reads generated in this study are archived and publicly available on the NCBI SRA database (BioProject PRJNA1094960). Panama samples (Vollmer et al 2023) can be sourced from BioProject PRJNA950067. Scripts for read processing, genotyping, and population genetic analyses are publicly available in the following Github repositories: https://github.com/mruggeri55/Acer-HiCoWGS-popgen, https://github.com/paigeduffin/Acer-HiCoWGS-popgen, https://github.com/TrinityConn/Acer-HiCoWGS-popgen.

## Paperpile references

Aitken SN, Whitlock MC. 2013. Assisted gene flow to facilitate local adaptation to climate change. Annual Review of Ecology, Evolution, and Systematics 44:367–388. Annual Reviews.

Alexander DH, Novembre J, Lange K. 2009. Fast model-based estimation of ancestry in unrelated individuals. Genome Research 19:1655–1664. Cold Spring Harbor Laboratory.

Baker AC et al. 2025. Proactive assisted gene flow for Caribbean corals in an era of rapid coral reef decline. Science (New York, N.Y.) 389:344–347. American Association for the Advancement of Science (AAAS).

Baums IB. 2008. A restoration genetics guide for coral reef conservation. Molecular Ecology 17:2796–2811. Wiley.

Baums IB et al. 2019. Considerations for maximizing the adaptive potential of restored coral populations in the western Atlantic. Ecological Applications 29:e01978. Wiley.

Baums IB, Locatelli NS, de Luca KL, Kitchen SA. 2026, April 18. Population structure and gene flow in the endangered Caribbean reef-building coral, Acropora palmata. bioRxiv. Available from 10.64898/2026.04.15.718759.

Baums IB, Miller MW, Hellberg ME. 2005. Regionally isolated populations of an imperiled Caribbean coral, Acropora palmata. Molecular Ecology 14:1377–1390. Wiley.

Bell DA, Robinson ZL, Funk WC, Fitzpatrick SW, Allendorf FW, Tallmon DA, Whiteley AR. 2019. The exciting potential and remaining uncertainties of genetic rescue. Trends in Ecology & Evolution 34:1070–1079. Elsevier BV.

Bertorelle G, Raffini F, Bosse M, Bortoluzzi C, Iannucci A, Trucchi E, Morales HE, van Oosterhout C. 2022. Genetic load: genomic estimates and applications in non-model animals. Nature Reviews. Genetics 23:492–503. Springer Science and Business Media LLC.

Browne L, Wright JW, Fitz-Gibbon S, Gugger PF, Sork VL. 2019. Adaptational lag to temperature in valley oak (Quercus lobata) can be mitigated by genome-informed assisted gene flow. Proceedings of the National Academy of Sciences of the United States of America 116:25179–25185. Proceedings of the National Academy of Sciences.

Browning BL, Tian X, Zhou Y, Browning SR. 2021. Fast two-stage phasing of large-scale sequence data. American Journal of Human Genetics 108:1880–1890. Elsevier BV.

Browning SR, Browning BL. 2007. Rapid and accurate haplotype phasing and missing-data inference for whole-genome association studies by use of localized haplotype clustering. American Journal of Human Genetics 81:1084–1097. Elsevier BV.

Chang CC, Chow CC, Tellier LC, Vattikuti S, Purcell SM, Lee JJ. 2015. Second-generation PLINK: rising to the challenge of larger and richer datasets. GigaScience 4:7. Oxford University Press (OUP).

Charlesworth D, Willis JH. 2009. The genetics of inbreeding depression. Nature Reviews. Genetics 10:783–796. Springer Science and Business Media LLC.

Conn T, Renton J, Chamberland V, Dellaert Z, Reusch TBH, Werner B, Baums I. 2025, April 15. Mosaic accumulation of somatic genetic variation and estimates of age in the long-lived reef-building coral Acropora palmata. Available from https://www.biorxiv.org/content/10.1101/2025.04.11.641527v1.abstract (accessed July 13, 2026).

Cook CN, Sgrò CM. 2019. Poor understanding of evolutionary theory is a barrier to effective conservation management. Conservation Letters 12:e12619. Wiley.

Cramer KL, Jackson JBC, Donovan MK, Greenstein BJ, Korpanty CA, Cook GM, Pandolfi JM. 2020. Widespread loss of Caribbean acroporid corals was underway before coral bleaching and disease outbreaks. Science Advances 6:eaax9395. American Association for the Advancement of Science (AAAS).

CRC Genetics Working Group. 2025. Safeguarding Florida’s coral reefs: The urgency of Assisted Gene Flow for Elkhorn coral conservation. Coral Restoration Consortium. Available from 10.5281/ZENODO.14920439.

Cunning R, Baker AC. 2013. Excess algal symbionts increase the susceptibility of reef corals to bleaching. Nature Climate Change 3:259–262. Springer Science and Business Media LLC.

Danecek P et al. 2021. Twelve years of SAMtools and BCFtools. GigaScience 10:giab008. Oxford University Press (OUP).

DeBuysser J, Alexander K, Hertler H. 2025, August 18. Mortality and recovery rates of Acropora fragments on in-situ nurseries after the 2023 bleaching event in the Turks and Caicos Islands. Available from https://assets-eu.researchsquare.com/files/rs-7301952/v1/af30ed0a-4206-4bcc-9673-6c6df5532036.pdf.

DeWoody JA, Harder AM, Mathur S, Willoughby JR. 2021. The long-standing significance of genetic diversity in conservation. Molecular Ecology 30:4147–4154. Wiley.

Dixon P. 2003. VEGAN, a package of R functions for community ecology. Journal of Vegetation Science: Official Organ of the International Association for Vegetation Science 14:927–930. Wiley.

Dray S, Dufour A-B. 2007. Theade4Package: Implementing the duality diagram for ecologists. Journal of Statistical Software 22:1–20. Foundation for Open Access Statistic.

Drury C, Dale KE, Panlilio JM, Miller SV, Lirman D, Larson EA, Bartels E, Crawford DL, Oleksiak MF. 2016. Genomic variation among populations of threatened coral: Acropora cervicornis. BMC Genomics 17:286. Springer Science and Business Media LLC.

Duforet-Frebourg N, Bazin E, Blum MGB. 2014. Genome scans for detecting footprints of local adaptation using a Bayesian factor model. Molecular Biology and Evolution 31:2483–2495. Oxford University Press (OUP).

Edmands S. 2007. Between a rock and a hard place: evaluating the relative risks of inbreeding and outbreeding for conservation and management: RELATIVE RISKS OF INBREEDING AND OUTBREEDING. Molecular Ecology 16:463–475. Wiley.

Evanno G, Regnaut S, Goudet J. 2005. Detecting the number of clusters of individuals using the software STRUCTURE: a simulation study. Molecular Ecology 14:2611–2620. Wiley.

Frankham R, Admin B. 2017, December 26. Backstory of: Genetic rescue of small inbred populations: meta-analysis reveals large and consistent benefits of gene flow. Available from 10.22541/au.151425005.59846175.

Frankham R, Ballou JD, Eldridge MDB, Lacy RC, Ralls K, Dudash MR, Fenster CB. 2011. Predicting the probability of outbreeding depression. Conservation Biology: The Journal of the Society for Conservation Biology 25:465–475. Wiley.

Frankham R, Bradshaw CJA, Brook BW. 2014. Genetics in conservation management: Revised recommendations for the 50/500 rules, Red List criteria and population viability analyses. Biological Conservation 170:56–63. Elsevier BV.

Franklin IA. 1980. Conservation biology, an evolutionary-ecological perspective.

Franklin IR, Allendorf FW, Jamieson IG. 2014. The 50/500 rule is still valid-Reply to Frankham et al. Biological Conservation 176:284–285. ui.adsabs.harvard.edu.

Frantz AC, Cellina S, Krier A, Schley L, Paris J. 2009. Using spatial Bayesian methods to determine the genetic structure of a continuously distributed population: clusters or isolation by distance? The Journal of Applied Ecology 46:493–505. Wiley.

Frichot E, François O. 2015. LEA: An R package for landscape and ecological association studies. Methods in Ecology and Evolution 6:925–929. Wiley.

Frichot E, Mathieu F, Trouillon T, Bouchard G, François O. 2014. Fast and efficient estimation of individual ancestry coefficients. Genetics 196:973–983. Oxford University Press (OUP).

Fuller ZL et al. 2020. Population genetics of the coral Acropora millepora: Toward genomic prediction of bleaching. Science (New York, N.Y.) 369:eaba4674. American Association for the Advancement of Science (AAAS).

Funk WC, McKay JK, Hohenlohe PA, Allendorf FW. 2012. Harnessing genomics for delineating conservation units. Trends in Ecology & Evolution 27:489–496. Elsevier BV.

Gagnaire P-A, Broquet T, Aurelle D, Viard F, Souissi A, Bonhomme F, Arnaud-Haond S, Bierne N. 2015. Using neutral, selected, and hitchhiker loci to assess connectivity of marine populations in the genomic era. Evolutionary Applications 8:769–786. Wiley.

García-Urueña R, Kitchen SA, Schizas NV. 2022. Fine scale population structure of Acropora palmata and Acropora cervicornis in the Colombian Caribbean. PeerJ 10:e13854. PeerJ.

Gargiulo R, Budde KB, Heuertz M. 2025. Mind the lag: understanding genetic extinction debt for conservation. Trends in Ecology & Evolution 40:228–237. Elsevier BV.

Garner A, Rachlow JL, Hicks JF. 2005. Patterns of genetic diversity and its loss in mammalian populations: Mammalian genetic diversity. Conservation Biology: The Journal of the Society for Conservation Biology 19:1215–1221. Wiley.

Gaunt TR, Rodríguez S, Day IN. 2007. Cubic exact solutions for the estimation of pairwise haplotype frequencies: implications for linkage disequilibrium analyses and a web tool “CubeX.” BMC Bioinformatics 8:428. Springer Science and Business Media LLC.

Gayral P et al. 2013. Reference-free population genomics from next-generation transcriptome data and the vertebrate-invertebrate gap. PLoS Genetics 9:e1003457. Public Library of Science (PLoS).

Gladfelter WB. 1982. White-band disease in Acropora palmata: Implications for the structure and growth of shallow reefs. Bull. Mar. Sci 32:639–643.

Goergen EA, Moulding AL, Walker BK, Gilliam DS. 2019. Identifying causes of temporal changes in Acropora cervicornis populations and the potential for recovery. Frontiers in Marine Science 6. Frontiers Media SA. Available from 10.3389/fmars.2019.00036.

Grossen C, Guillaume F, Keller LF, Croll D. 2020. Purging of highly deleterious mutations through severe bottlenecks in Alpine ibex. Nature Communications 11:1001. Springer Science and Business Media LLC.

Grossen C, Ramakrishnan U. 2024. Genetic load. Current Biology 34:R1216–R1220. Elsevier BV.

Hagedorn M et al. 2021. Assisted gene flow using cryopreserved sperm in critically endangered coral. Proceedings of the National Academy of Sciences of the United States of America 118:e2110559118. Proceedings of the National Academy of Sciences.

Hare MP, Nunney L, Schwartz MK, Ruzzante DE, Burford M, Waples RS, Ruegg K, Palstra F. 2011. Understanding and estimating effective population size for practical application in marine species management: Applying effective population size estimates to marine species management. Conservation Biology: The Journal of the Society for Conservation Biology 25:438–449. Wiley.

Hedrick PW, Garcia-Dorado A. 2016. Understanding inbreeding depression, purging, and genetic rescue. Trends in Ecology & Evolution 31:940–952. Elsevier BV.

Highsmith RC. 1982. Reproduction by Fragmentation in Corals. Marine Ecology Progress Series 7:207–226. Inter-Research Science Center.

Hoffmann AA, Willi Y. 2008. Detecting genetic responses to environmental change. Nature Reviews. Genetics 9:421–432. Springer Science and Business Media LLC.

Hughes TP, Tanner JE. 2000. RECRUITMENT FAILURE, LIFE HISTORIES, AND LONG-TERM DECLINE OF CARIBBEAN CORALS. Ecology 81:2250–2263.

Jackson J. 2014. Global Coral Reef Monitoring Network; International Union for the Conservation of. Nature:1970–2012. IUCN.

Jamieson IG, Allendorf FW. 2012. How does the 50/500 rule apply to MVPs? Trends in Ecology & Evolution 27:578–584. Elsevier BV.

Jombart T, Devillard S, Balloux F. 2010. Discriminant analysis of principal components: a new method for the analysis of genetically structured populations. BMC Genetics 11:94.

Kardos M, Armstrong EE, Fitzpatrick SW, Hauser S, Hedrick PW, Miller JM, Tallmon DA, Funk WC. 2021. The crucial role of genome-wide genetic variation in conservation. Proceedings of the National Academy of Sciences of the United States of America 118:e2104642118. Proceedings of the National Academy of Sciences.

Kiel PM, Karp R, Jankulak M, Enochs IC. 2025, March 25. Acropora cervicornis Data Coordination Hub (AcDC), an open-access tool for aligning datasets and evaluating genet-specific performance. Available from 10.5194/oos2025-371.

Knowlton N, Lang JC, Rooney M, Clifford P. 1981. Evidence for Delayed Mortality in Hurricane-Damaged Jamaican Staghom Corals. Nature 294:251–252.

Kopelman NM, Mayzel J, Jakobsson M, Rosenberg NA, Mayrose I. 2015. Clumpak: a program for identifying clustering modes and packaging population structure inferences across K. Molecular Ecology Resources 15:1179–1191. Wiley.

Korunes KL, Samuk K. 2021. pixy: Unbiased estimation of nucleotide diversity and divergence in the presence of missing data. Molecular Ecology Resources 21:1359–1368. Wiley.

Kwong AM et al. 2021. Robust, flexible, and scalable tests for Hardy-Weinberg equilibrium across diverse ancestries. Genetics 218:iyab044. Oxford University Press (OUP).

Kyriazis CC, Robinson JA, Lohmueller KE. 2025. Long runs of homozygosity are reliable genomic markers of inbreeding depression. Trends in Ecology & Evolution 40:874–884. Elsevier BV.

Kyriazis CC, Wayne RK, Lohmueller KE. 2021. Strongly deleterious mutations are a primary determinant of extinction risk due to inbreeding depression. Evolution Letters 5:33–47. Oxford University Press (OUP).

Lawson DJ, van Dorp L, Falush D. 2018. A tutorial on how not to over-interpret STRUCTURE and ADMIXTURE bar plots. Nature Communications 9:3258. Springer Science and Business Media LLC.

Le Gall L, Johnson JV, Goodbody-Gringley G. 2025. Extirpation of Acropora cervicornis genotypes from a coral nursery during the 2023 marine heatwave undermines conservation efforts. Frontiers in Marine Science 12. Frontiers Media SA. Available from 10.3389/fmars.2025.1599155.

Liddell E, Sunnucks P, Cook CN. 2021. To mix or not to mix gene pools for threatened species management? Few studies use genetic data to examine the risks of both actions, but failing to do so leads …. Biological Conservation. Elsevier. Available from https://www.sciencedirect.com/science/article/pii/S0006320721001245.

Li H. 2011. A statistical framework for SNP calling, mutation discovery, association mapping and population genetical parameter estimation from sequencing data. Bioinformatics (Oxford, England) 27:2987–2993. Oxford University Press (OUP).

Li H. 2013, March 16. Aligning sequence reads, clone sequences and assembly contigs with BWA-MEM. Available from http://arxiv.org/abs/1303.3997.

Lirman D, Schopmeyer S. 2016. Ecological solutions to reef degradation: optimizing coral reef restoration in the Caribbean and Western Atlantic. PeerJ 4:e2597. PeerJ.

Locatelli NS, Kitchen SA, Stankiewicz KH, Osborne CC, Dellaert Z, Elder H, Kamel B, Koch HR, Fogarty ND, Baums IB. 2024. Chromosome-level genome assemblies and genetic maps reveal heterochiasmy and macrosynteny in endangered Atlantic Acropora. BMC Genomics 25:1119. Springer Science and Business Media LLC.

Luu K, Bazin E, Blum MGB. 2017. pcadapt: an R package to perform genome scans for selection based on principal component analysis. Molecular Ecology Resources 17:67–77. Wiley.

Manzello DP et al. 2025. Heat-driven functional extinction of Caribbean Acropora corals from Florida’s Coral Reef. Science (New York, N.Y.) 390:361–366. American Association for the Advancement of Science (AAAS).

Martin M, Patterson M, Garg S, O Fischer S, Pisanti N, Klau G, Schöenhuth A, Marschall T. 2016. WhatsHap: fast and accurate read-based phasing. bioRxiv. Available from 10.1101/085050.

Mathur S, Gibbs HL. 2025. Genomic evaluation of assisted gene flow options in an endangered rattlesnake. Molecular Ecology 34:e70014. Wiley.

Matz MV, Treml EA, Aglyamova GV, Bay LK. 2018. Potential and limits for rapid genetic adaptation to warming in a Great Barrier Reef coral. PLoS Genetics 14:e1007220.

Mills LS. 2012. Conservation of Wildlife Populations: Demography, genetics, and management. Wiley, New Delhi, India.

Moritz C. 2002. Strategies to protect biological diversity and the evolutionary processes that sustain it Systematic Biology 51:238–254. Oxford University Press (OUP).

Muller EM et al. 2026. Success of restoration strategies in preventing extirpation of 2 critically endangered coral species. Conservation Biology: The Journal of the Society for Conservation Biology 40:e70168. Wiley.

Muller E, Petrik C, Conn T, Osborne C, Villoch M, Clark A, Koch H, O’Neil K, Engelsma C, Baums I. 2025, March 11. Assisted gene flow yields *Acropora palmata* corals with robust physiological performance under warmer water temperatures in a land-based nursery. bioRxiv. Available from https://www.biorxiv.org/content/10.1101/2025.03.03.641242v1.abstract (accessed August 11, 2026).

National Academies of Sciences Engineering, Medicine. 2019. A Research Review of Interventions to Increase the Persistence and Resilience of Coral Reefs. The National Academies Press, Washington, DC.

Novak BJ, Phelan R, Weber M. 2021. US conservation translocations: Over a century of intended consequences. Conservation Science and Practice 3.

Ørsted M, Hoffmann AA, Sverrisdóttir E, Nielsen KL, Kristensen TN. 2019. Genomic variation predicts adaptive evolutionary responses better than population bottleneck history. PLoS Genetics 15:e1008205. Public Library of Science (PLoS).

Palacio-Castro AM, Dennison CE, Rosales SM, Baker AC. 2021. Variation in susceptibility among three Caribbean coral species and their algal symbionts indicates the threatened staghorn coral, Acropora cervicornis, is particularly susceptible to elevated nutrients and heat stress. Coral Reefs 40:1601–1613. Springer Science and Business Media LLC.

Palsbøll PJ, Bérubé M, Allendorf FW. 2007. Identification of management units using population genetic data. Trends in Ecology & Evolution 22:11–16. Elsevier BV.

Pembleton LW, Cogan NOI, Forster JW. 2013. StAMPP: an R package for calculation of genetic differentiation and structure of mixed-ploidy level populations. Molecular Ecology Resources 13:946–952. Wiley.

Perez MF, Franco FF, Bombonato JR, Bonatelli IAS, Khan G, Romeiro-Brito M, Fegies AC, Ribeiro PM, Silva GAR, Moraes EM. 2018. Assessing population structure in the face of isolation by distance: Are we neglecting the problem? Diversity & Distributions 24:1883– 1889. Wiley.

Petkova D, Novembre J, Stephens M. 2016. Visualizing spatial population structure with estimated effective migration surfaces. Nature Genetics 48:94–100. Springer Science and Business Media LLC.

Picard. (n.d.). Available from http://broadinstitute.github.io/picard (accessed June 9, 2026).

Pritchard JK, Stephens M, Donnelly P. 2000. Inference of population structure using multilocus genotype data. Genetics 155:945–959. Oxford University Press (OUP).

Privé F, Luu K, Vilhjálmsson BJ, Blum MGB. 2020. Performing highly efficient genome scans for local adaptation with R package pcadapt version 4. Molecular Biology and Evolution 37:2153–2154. Oxford University Press (OUP).

Ralls K, Ballou JD, Dudash MR, Eldridge MDB, Fenster CB, Lacy RC, Sunnucks P, Frankham R. 2018. Call for a paradigm shift in the genetic management of fragmented populations: Genetic management. Conservation Letters 11:e12412. Wiley.

Reed DH, Frankham R. 2003. Correlation between fitness and genetic diversity. Conservation Biology: The Journal of the Society for Conservation Biology 17:230–237. Wiley.

Reich HG, Kitchen SA, Stankiewicz KH, Devlin-Durante M, Fogarty ND, Baums IB. 2021. Genomic variation of an endosymbiotic dinoflagellate (Symbiodinium “fitti”) among closely related coral hosts. Molecular Ecology 30:3500–3514. Wiley.

Rivers N, Daly J, Temple-Smith P. 2020. New directions in assisted breeding techniques for fish conservation.source> Reproduction, Fertility, and Development 32:807–821. CSIRO Publishing.

Robinson JA, Bowie RCK, Dudchenko O, Aiden EL, Hendrickson SL, Steiner CC, Ryder OA, Mindell DP, Wall JD. 2021. Genome-wide diversity in the California condor tracks its prehistoric abundance and decline. Current Biology 31:2939–2946.e5. Elsevier BV.

Robinson J, Kyriazis CC, Yuan SC, Lohmueller KE. 2023. Deleterious variation in natural populations and implications for conservation genetics. Annual Review of Animal Biosciences 11:93–114. Annual Reviews.

Sadler AJ, Williams BRG. 2008. Interferon-inducible antiviral effectors. Nature Reviews. Immunology 8:559–568. Springer Science and Business Media LLC.

Santiago E, Köpke C, Caballero A. 2025. Accounting for population structure and data quality in demographic inference with linkage disequilibrium methods. Nature Communications 16:6054. Springer Science and Business Media LLC.

Schmidt TL, Thia JA, Hoffmann AA. 2024. How can genomics help or hinder wildlife conservation? Annual Review of Animal Biosciences 12:45–68. Annual Reviews.

Schopmeyer SA et al. 2012. In situ coral nurseries serve as genetic repositories for coral reef restoration after an extreme Cold-Water event. Restoration Ecology 20:696–703. Wiley.

Supple MA, Shapiro B. 2018. Conservation of biodiversity in the genomics era. Genome Biology 19:131. Springer Science and Business Media LLC.

Szpiech ZA, Blant A, Pemberton TJ. 2017. GARLIC: Genomic autozygosity regions likelihood-based inference and classification. Bioinformatics (Oxford, England) 33:2059–2062. Oxford University Press (OUP).

Taylor BL, Dizon AE. 1999. First policy then science: why a management unit based solely on genetic criteria cannot work. Molecular Ecology 8:S11–6. Wiley.

Terhorst J, Kamm JA, Song YS. 2017. Robust and scalable inference of population history from hundreds of unphased whole genomes. Nature Genetics 49:303–309. Springer Science and Business Media LLC.

van Oosterhout C, Speak SA, Birley T, Hitchings LWG, Bortoluzzi C, Percival-Alwyn L, Urban L, Groombridge JJ, Segelbacher G, Morales HE. 2026. Genomic erosion in the assessment of species’ extinction risk and recovery potential. The Journal of Heredity:esag011. Oxford University Press (OUP).

Vasquez Kuntz KL, Kitchen SA, Conn TL, Vohsen SA, Chan AN, Vermeij MJA, Page C, Marhaver KL, Baums IB. 2022. Inheritance of somatic mutations by animal offspring. Science Advances 8:eabn0707. American Association for the Advancement of Science (AAAS).

Vollmer SV, Palumbi SR. 2007. Restricted gene flow in the Caribbean staghorn coral Acropora cervicornis: implications for the recovery of endangered reefs. The Journal of Heredity 98:40–50. Oxford University Press (OUP).

Vollmer SV, Selwyn JD, Despard BA, Roesel CL. 2023. Genomic signatures of disease resistance in endangered staghorn corals. Science (New York, N.Y.) 381:1451–1454.

Wang J. 2025. PopCluster: A population genetics model-based toolset for simulating, inferring and visualising individual admixture and population structure. Molecular Ecology Resources 25:e14058. Wiley.

Waples RS. 1991. Pacific salmon, Oncorhynchus spp., and the definition of“ species” under the Endangered Species Act. Marine Fisheries Review 53:11–22. books.google.com.

Weeks AR, Heinze D, Perrin L, Stoklosa J, Hoffmann AA, van Rooyen A, Kelly T, Mansergh I. 2017. Genetic rescue increases fitness and aids rapid recovery of an endangered marsupial population. Nature Communications 8:1071. Springer Science and Business Media LLC.

Weeks AR, Stoklosa J, Hoffmann AA. 2016. Conservation of genetic uniqueness of populations may increase extinction likelihood of endangered species: the case of Australian mammals. Frontiers in Zoology 13:31. Springer Science and Business Media LLC.

Weir BS, Cockerham CC. 1984. Estimating F-statistics for the analysis of population structure. Evolution; International Journal of Organic Evolution 38:1358. Oxford University Press (OUP).

Werden LK, Sugii NC, Weisenberger L, Keir MJ, Koob G, Zahawi RA. 2020. Ex situ conservation of threatened plant species in island biodiversity hotspots: A case study from Hawai ‘i. Biological Conservation.

Whitlock MC, Lotterhos KE. 2015. Reliable Detection of Loci Responsible for Local Adaptation: Inference of a Null Model through Trimming the Distribution of FST. The American Naturalist 186:S24–S36.

Williams DE, Miller MW, Kramer KL. 2008. Recruitment failure in Florida Keys Acropora palmata, a threatened Caribbean coral. Coral Reefs 27:697–705. Springer Science and Business Media LLC.

Williams DE, Nedimyer K, Miller MW. 2020. Genotypic inventory of Acropora palmata (elkhorn coral) populations in south Florida. NOAA National Marine Fisheries Service Southeast Fisheries Science Center Protected Resources and Biodiversity Division Report.

Willi Y, Van Buskirk J, Fischer M. 2005. A threefold genetic allee effect: population size affects cross-compatibility, inbreeding depression and drift load in the self-incompatible Ranunculus reptans: Population size affects cross-compatibility, inbreeding depression and drift load in the self-incompatible Ranunculus reptans Genetics 169:2255–2265. Oxford University Press (OUP).

Wilson K et al. 2002. Genetic mapping of the black tiger shrimp Penaeus monodon with amplified fragment length polymorphism. Aquaculture 204:297–309.

Wylie MJ, Kitson J, Russell K, Yoshizaki G, Yazawa R, Steeves TE, Wellenreuther M. 2025. Fish germ cell cryobanking and transplanting for conservation. Molecular Ecology Resources 25:e13868.

## In press

Duffin PJ, Schiebelhut LM, Dawson MN, Wares JP. In press. Genomic separation of Salish Sea and Pacific outer coast populations of the keystone sea star *Pisaster ochraceus*. Evolution.

