## Supplementary Materials for "Limited neutral and adaptive genomic divergence suggests *Acropora cervicornis* can be managed as a single conservation unit across its range"

### Appendix S1. Sample and sequence processing methods.

#### Appendix S1.1. Detailed methods for processing reads, calling genotypes, and filtering loci.

For newly sequenced samples, raw reads were trimmed using Trim Galore v0.6.6 (Krueger 2015) to remove sequencing adapters, low-quality ends with Phred scores <20, and 5′ bias. Reads were then filtered with FASTX-Toolkit to retain reads with ≥99% base-call accuracy across at least 90% of the read. Paired-end reads were separated into read pairs for which both mates passed filtering and reads for which only one mate passed filtering. Both paired and unpaired reads were mapped independently to a combined *A. cervicornis* host (assembly jaAcrCerv1.1; NCBI accession GCA_964034985.1) and *S. fitti* (Reich et al. 2021) genome reference using the BWA-mem algorithm (Li 2013). Paired and single-end mapped reads were merged and PCR duplicates were marked using the Picard v2.26.2 (“Picard” n.d.) MarkDuplicates tool.

Because Panama samples were generated independently, their initial read processing followed (Vollmer et al. 2023). Briefly, sequencing adapters, low-quality ends, poly-G and poly-X tails were trimmed using fastp. Reads with >40% unqualified bases, lengths <140 bp, or low sequence complexity were removed. To standardize these data with newly generated samples, Panama reads were then subjected to the same downstream read-quality filtering used for all other samples, retaining only reads with ≥99% base-call accuracy (Phred score of 20 or higher) across at least 90% of the read using a custom Python script (Appendix S1.2). Panama reads were then mapped and duplicate-marked using the same procedure described above.

Genotypes were called for both variant and invariant loci using bcftools v1.19 mpileup and multiallelic calling functions (Li 2011; Danecek et al. 2021). Sample-specific genotype files were merged with bcftools merge. Variant and invariant loci were filtered for depth, quality, and missingness (MEAN(FMT/DP)>=10 && MEAN(FMT/DP)<=50 && QUAL>20 && GQ>20 && MQ>40 && F_MISSING<0.1). SNPs were additionally filtered to exclude those within 5bp of indels, strand bias (SOR>4), and those deviating from Hardy–Weinberg equilibrium after accounting for population structure using RUTH v3.29.4 (HWE_SLP_I≤−5 or ≥5; approximately $p<1\times10^{-5}$) (Kwong et al. 2021). See Figure S1.1 for RUTH workflow overview.

To screen publicly available Panama samples for putative clones, pairwise KING kinship coefficients were calculated with PLINK2. Sample pairs with kinship coefficients >0.3 were considered clonal, and 10 unique Panama genotypes were randomly retained for downstream analyses (Table S1.1). One Curaçao individual, CU21E_1049, was excluded from all downstream analyses because preliminary population-genetic screening identified it as likely contaminated, based on an extremely negative inbreeding coefficient and ambiguous population assignment.

#### Appendix S1.2. Trimming step for Panama samples (based on trimming in sample source article Vollmer et al. 2023).

```
fastp \
 --in1 ./raw_reads/${ID}_1.fastq.gz --in2 ./raw_reads/${ID}_2.fastq.gz \
 --out1 ./fastp_trimmed/${ID}_1_fastp.trimmed.fastq.gz --out2
    ./fastp_trimmed/${ID}_2_fastp.trimmed.fastq.gz \
  --qualified_quality_phred 20 --unqualified_percent_limit 40 \
  --length_required 140 \
```

```
--low_complexity_filter --complexity_threshold 30 \
--detect_adapter_for_pe \
--cut_tail --cut_tail_window_size 1 --cut_tail_mean_quality 20 \
--cut_front --cut_front_window_size 1 --cut_front_mean_quality 20 \
--cut_right --cut_right_window_size 10 --cut_right_mean_quality 20 \
--trim_poly_g --poly_g_min_len 10 --trim_poly_x \
--correction --unqualified_percent_limit 40 \
-h ./reports/${ID}.fastp.html -j ./reports/${ID}.fastp.json
```

*Python code for Panama samples' quality filtering step (as alternative to fastx-toolkit)*

fastx-toolkit version code summary: `cat input.fastq | fastq_quality_filter -q 20 -p 90 > output.clean`

Custom py script name: `qual.filt_90perc.over.20.py`

Usage example: `python3 qual.filt_90perc.over.20.py input.fastq output.clean`

Py script contents:

```
import argparse
def phred_quality_filter(input_file, output_file, quality_threshold=20,
min_percent_high_quality=0.9):
    with open(input_file, "r") as infile, open(output_file, "w") as outfile:
        while True:
            header = infile.readline().strip()
            if not header:
                break
            sequence = infile.readline().strip()
            plus = infile.readline().strip()
            quality = infile.readline().strip()
            high_quality_bases = sum(1 for char in quality if ord(char) - 33 >=
quality_threshold)
            high_quality_percentage = high_quality_bases / len(quality)
            if high_quality_percentage >= min_percent_high_quality:
                outfile.write(f"{header}\n{sequence}\n{plus}\n{quality}\n")
if __name__ == "__main__":
    parser = argparse.ArgumentParser(description="Filter FASTQ file based on Phred
quality scores.")
    parser.add_argument("input_file", help="Input FASTQ file")
    parser.add_argument("output_file", help="Output file for filtered reads")
    args = parser.parse_args()
    phred_quality_filter(args.input_file, args.output_file)
```

*Quality filtering python script validation*

Of the 10 Panama samples, haphazardly selected a subset of 2 samples (with 2x paired read files each) to validate python script function against fastx-toolkit version. Files were compared using the Unix `diff` function (e.g. `diff sample_fastxtoolkit.clean sample_pyscript.clean > print_differences.txt`), resulting in no identified differences. All 10 quality-filtered samples were compared across both methods to ensure identical line number and character number outputs across either method (`wc -l sample.clean` and `wc -m sample.clean`, respectively).

After these initial processing steps, Panama samples were subjected to the same read-quality filtering criteria as the remaining 36 newly acquired WGS samples sourced from diverse Caribbean reefs.

**Table S1.1. Selected metadata on 10 Panama samples sourced from Vollmer et al. (2023).** Adapted from the original study to include key sample identifier information collated across several supplemental files (https://www.science.org/doi/10.1126/science.adi3601, https://doi.org/10.5281/zenodo.8095056). The "Sample_ID" column lists identifiers assigned by Vollmer et al. (2023) and retained in this study. Columns in grey denote information not strictly relevant to the context of this study but included as potentially useful sample metadata for future reproductions.

| Sample_ID | LibraryName | Region | Reef | Reef_ID | Lat | Lon | Experiment |
|---|---|---|---|---|---|---|---|
| SRR24007609 | Ac_PA_CK140 | Panama | CK14 | CK140 | 9.25396667 | -82.12595 | SRX19811004 |
| SRR24007608 | Ac_PA_CK1410 | Panama | CK14 | CK1410 | 9.25396667 | -82.12595 | SRX19811005 |
| SRR24007601 | Ac_PA_CK146 | Panama | CK14 | CK146 | 9.25396667 | -82.12595 | SRX19811012 |
| SRR24007603 | Ac_PA_CK144 | Panama | CK14 | CK144 | 9.25396667 | -82.12595 | SRX19811010 |
| SRR24007602 | Ac_PA_CK145 | Panama | CK14 | CK145 | 9.25396667 | -82.12595 | SRX19811011 |
| SRR24007597 | Ac_PA_CK42 | Panama | CK4 | CK42 | 9.25861667 | -82.127083 | SRX19811016 |
| SRR24007659 | Ac_PA_SR2 | Panama | SR | SR2 | 9.25456667 | -82.127183 | SRX19810954 |
| SRR24007652 | Ac_PA_SR8 | Panama | SR | SR8 | 9.25456667 | -82.127183 | SRX19810961 |
| SRR24007661 | Ac_PA_SR10 | Panama | SR | SR10 | 9.25456667 | -82.127183 | SRX19810952 |
| SRR24007593 | Ac_PA_Tet2 | Panama | Tet | Tet2 | 9.27631667 | -82.101133 | SRX19810964 |

**Table S1.2. Master list of samples used across various analyses included in this study.** General description and usage of the three main sample lists used in this study followed by detailed information on samples included in each list. Mean sequencing depth per sample at the bam file stage also provided, calculated over all loci contained within the main 14 chromosomes ("all retained chromos") and after filtering to only include high-confidence SNPs retained for the haplotype reference panel ("final SNPs in HRP"). Sampling location abbreviations include ARU: Aruba; FLU: Upper Florida Keys; CUR: Curaçao; DOM: the Dominican Republic; DRT: Dry Tortugas; BEL: Belize; JAM: Jamaica; MEX: Mexico; FLL: Lower Florida Keys; and PAN: Panama.

| **Sample list n=46** | |
|---|---|
| *Description*: Main full sample list; Use(s) = Neutral SNP panel-based pop structure analyses; K=5 pairwise and pop-specific $F_{ST}$ calcs; Full SNP panel selection scans | *Samples removed*: all versions have CU21E_1049 removed due to suspected contamination |
| **Sample list n=45** | |
| *Description*: Use(s) = Genomic diversity (nucleotide diversity, π; Watterson's theta; Tajima's D; ROH) and $N_e$ estimates | *Samples removed*: all in n=46 list, plus: CRF_Acer-099 due to high admixture |

| Sample list n=44 | |
|---|---|
| *Description*: Use(s) = Haplotype reference panel | *Samples removed*: all in n=46 list, plus: CRF_Acer-099, SRR24007608 due to high admixture |

| Sample list n=37 | |
|---|---|
| *Description*: Use(s) = Sampling location based pairwise and pop-specific $F_{ST}$ calculations; IBD (sampling location based). Purpose = downsampling to reduce unequal representation (max n=4 per location) | *Samples removed*: all in n=46 list, plus:: DRTO_151, IBBZ_13837, JAM_11, SRR24007593, SRR24007597, SRR24007601, SRR24007603, SRR24007609, SRR24007652 |

**Sample list details table**

| sample ID | sample list | | | | sampling location | sequenced for… | mean depth per sample | |
|---|---|---|---|---|---|---|---|---|
| | n= 46 | n= 45 | n= 44 | n= 37 | | | all retained chromos | final SNPs in HRP |
| AR_113 | | | | | ARU | this study | 37.37 | 42.31 |
| AR_115 | | | | | ARU | this study | 39.09 | 44.27 |
| AR_117 | | | | | ARU | this study | 35.55 | 39.68 |
| AR_119 | | | | | ARU | this study | 40.67 | 45.87 |
| CRF_Acer-059 | | | | | FLU | this study | 38.94 | 42.74 |
| CRF_Acer-099 | | | | | FLU | this study | 38.88 | NA |
| CRF_Acer-102 | | | | | FLU | this study | 39.15 | 41.45 |
| CRF_Acer-120 | | | | | FLU | this study | 36.96 | 41.36 |
| CU21E_1050 | | | | | CUR | this study | 34.17 | 38.90 |
| CU21E_1051 | | | | | CUR | this study | 37.65 | 42.48 |
| CU21E_1052 | | | | | CUR | this study | 33.53 | 38.04 |
| CU21E_1053 | | | | | CUR | this study | 36.96 | 40.31 |
| DR_C012 | | | | | DOM | this study | 32.78 | 36.26 |
| DR_C051 | | | | | DOM | this study | 33.80 | 38.46 |
| DR_C080 | | | | | DOM | this study | 34.63 | 38.31 |
| DR_C171 | | | | | DOM | this study | 36.24 | 42.17 |
| DRTO_114 | | | | | DRT | this study | 38.62 | 44.64 |
| DRTO_143 | | | | | DRT | this study | 40.45 | 44.70 |
| DRTO_151 | | | | | DRT | this study | 39.90 | 46.08 |
| DRTO_178 | | | | | DRT | this study | 40.19 | 45.90 |
| DRTO_52 | | | | | DRT | this study | 41.99 | 46.07 |
| IBBZ_118 | | | | | BEL | this study | 35.86 | 41.53 |
| IBBZ_123 | | | | | BEL | this study | 38.91 | 45.38 |
| IBBZ_13797 | | | | | BEL | this study | 39.74 | 45.60 |
| IBBZ_13829 | | | | | BEL | this study | 41.60 | 46.37 |
| IBBZ_13837 | | | | | BEL | this study | 39.59 | 46.02 |
| JAM_01 | | | | | JAM | this study | 36.78 | 40.95 |
| JAM_02 | | | | | JAM | this study | 38.22 | 42.56 |

| | | | | | | | | |
|---|---|---|---|---|---|---|---|---|
| JAM_05 | | | | | JAM | this study | 35.22 | 40.04 |
| JAM_07 | | | | | JAM | this study | 37.14 | 41.56 |
| JAM_11 | | | | | JAM | this study | 36.66 | 41.55 |
| MEX_01 | | | | | MEX | this study | 33.32 | 37.66 |
| MEX_03 | | | | | MEX | this study | 38.34 | 42.92 |
| Mote_AC75 | | | | | FLL | this study | 35.76 | 41.42 |
| Mote_AC76 | | | | | FLL | this study | 38.15 | 42.78 |
| Mote_AC80 | | | | | FLL | this study | 37.63 | 42.20 |
| SRR24007593 | | | | | PAN | Vollmer (2023) | 64.27 | 75.08 |
| SRR24007597 | | | | | PAN | Vollmer (2023) | 53.08 | 63.23 |
| SRR24007601 | | | | | PAN | Vollmer (2023) | 47.48 | 57.03 |
| SRR24007602 | | | | | PAN | Vollmer (2023) | 40.61 | 48.94 |
| SRR24007603 | | | | | PAN | Vollmer (2023) | 46.85 | 54.51 |
| SRR24007608 | | | | | PAN | Vollmer (2023) | 55.36 | NA |
| SRR24007609 | | | | | PAN | Vollmer (2023) | 71.14 | 83.48 |
| SRR24007652 | | | | | PAN | Vollmer (2023) | 26.92 | 32.05 |
| SRR24007659 | | | | | PAN | Vollmer (2023) | 62.48 | 76.90 |
| SRR24007661 | | | | | PAN | Vollmer (2023) | 23.22 | 27.28 |

**Table S1.3. Depth summary stats across BAMs** used in population structure, demography, and adaptive analyses (n=46), with and without Panama ["PAN"] samples from Vollmer et al. (2023).

| for n=46 | average | std dev | max | min |
|---|---|---|---|---|
| all samples | 40.0401 | 8.75423 | 71.139 | 23.216 |
| exclud PAN | 37.5123 | 2.38432 | 41.9859 | 32.783 |

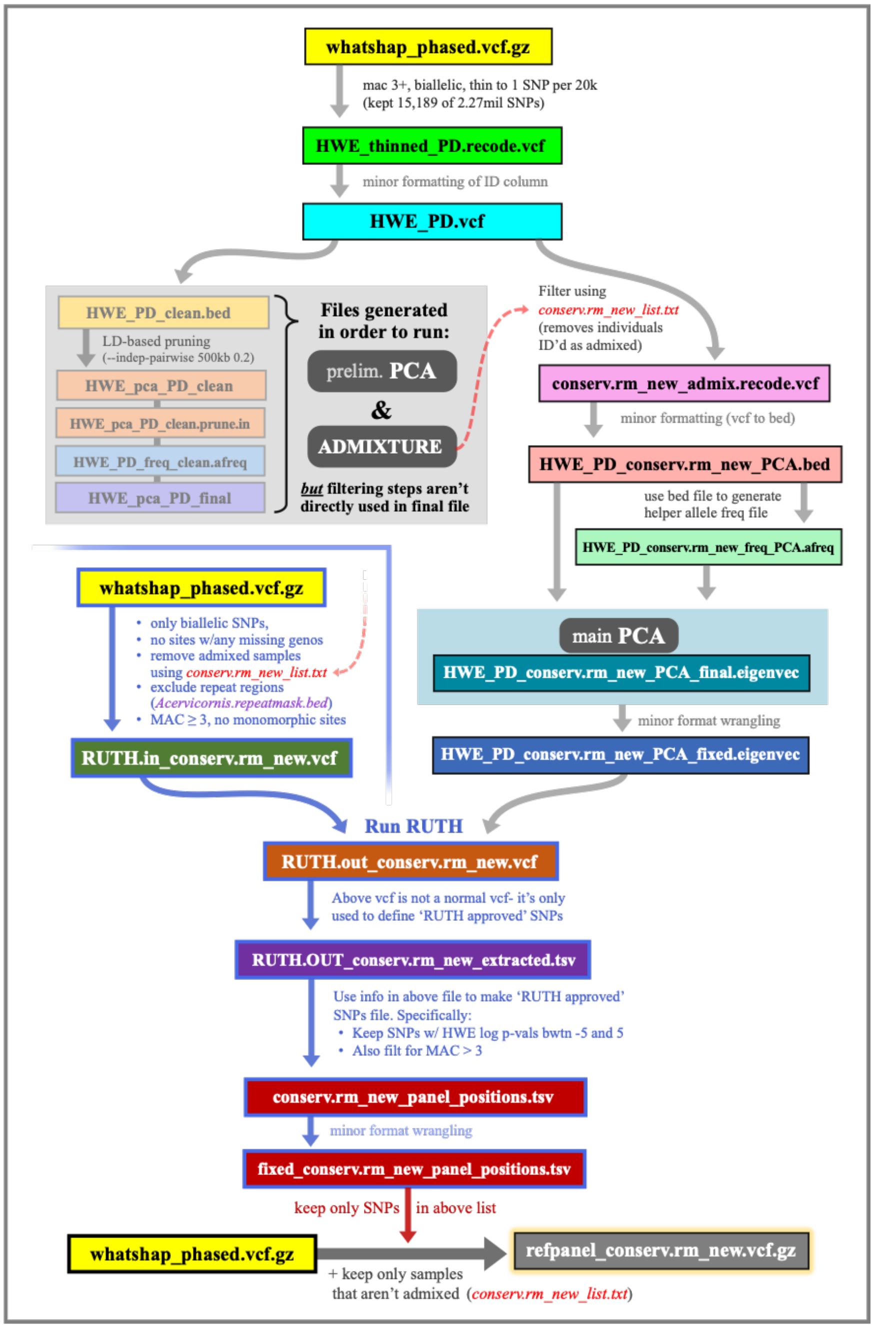

**Figure S1.1. RUTH HWE filtering workflow.** Detailed methods pipeline to identify and remove SNPs deviating from Hardy-Weinberg equilibrium while accounting for population structure using RUTH (Kwong et al., 2021, v3.29.4). This example tracks files undergoing RUTH filtering during the generation of the haplotype reference panel.

**Appendix S1.3. Detailed methods for neutral panel generation.**

For neutral population-structure analyses, SNPs were additionally filtered to include only bi-allelic loci (-m2) with allele counts greater than 3 (AC>3) using bcftools. Putative outlier loci were removed prior to neutral analyses using three steps (Table S1.4). First, 79,020 outliers were identified from initial PCA loadings (Figure S7.1) using PCAdapt v4.3.5 (Luu et al. 2017), which also informed post hoc groupings (K = 5) used as input for OutFLANK v0.2 (Whitlock & Lotterhos, 2017). Liberal identification of the lowest 0.1% of adjusted p-values (q-values < 0.7083018) in OutFLANK removed 974 SNPs. OutFLANK was then applied a second time with populations defined by geographic sampling locations (lowest 0.1% of q-values at < 0.3681863, Figure 1a), identifying 178 SNPs for removal. Collectively, and accounting for overlap between lists, these methods identified 79,041 SNPs for removal. SNPs in linkage disequilibrium were subsequently pruned using plink2 (--indep-pairwise 50 5 0.2; plink2/2.00a3.7LM), which resulted in the removal of 2,220,627 SNPs. Finally, SNPs were filtered to remove loci with >10% missing data and/or MAF <0.05, collectively removing another 50,542 SNPs. The final neutral SNP panel consisted of 177,516 putatively neutral variants (Table S1.4).

**Table S1.4. Outlier SNPs identified and removed in neutral SNP panel generation.**

| **File / Step Description** | **SNP Count** |
|---|---|
| Full quality-filtered SNP panel | 2,527,725 |
| Putative outliers removed | 79,041 |
| *PCAdapt (alpha=0.05)* | *79,020* |
| *OutFLANK (geographic locations, lowest 0.1% q-vals)* | *974* |
| *OutFLANK (K=5 post hoc groups, lowest 0.1% q-vals)* | *178* |
| SNPs removed in LD pruning | 2,220,627 |
| Quality filtering (>10% missing removed) | 16,249 |
| Quality filtering (<0.05 MAF removed) | 34,293 |
| **Final neutral SNP panel** | **177,516** |

### Appendix S2. Population structure methods.

**Appendix S2.1. Detailed methods for population structure, $F_{ST}$, IBD, and migration rate analyses.**

We applied multiple complementary approaches to evaluate population structure using the pruned neutral SNP panel. First, principal component analysis (PCA) was performed in R using PCAdapt (Luu et al. 2017), with scree plots used to confirm that the first two PCs captured the dominant axes of neutral genetic variation; PCA results were then visually assessed for discrete geographic or genetic groupings (Figure S7.3). We next used Discriminant Analysis of Principal Components (DAPC; Jombart et al. 2010), running 50 independent replicates across K = 1–10, with Bayesian information criterion (BIC) values used to evaluate model support. Individual ancestry coefficients were estimated using ADMIXTURE (Alexander et al. 2009); v1.3.0), with 20 random-seed runs tested for each value of K = 1–10 and 10-fold cross-validation enabled using the --cv=10 flag. ADMIXTURE results were evaluated using the software's cross-validation error and summarized with CLUMPAK (Kopelman et al. 2015), which provided replicate congruence, similarity statistics, and Evanno ΔK estimates (Evanno et al. 2005). Sparse Non-negative Matrix Factorization (sNMF; Frichot et al. 2014) was also used to estimate individual ancestry coefficients through the R package LEA v2.20 (Frichot & François 2015), again using 20 random-seed runs for each K = 1–10. sNMF results were evaluated using cross-entropy with the masked-data option, as well as CLUMPAK-based congruence, similarity statistics, and Evanno ΔK estimates. Lastly, we used PopCluster (Wang 2025) as an additional model-based approach, running five random-seed replicates for each K = 1–10. PopCluster results were evaluated using the software's $F_{ST}$/$F_{IS}$ and DLK2 outputs, together with CLUMPAK-based congruence, similarity statistics, and Evanno ΔK estimates.

Pairwise $F_{ST}$ was calculated from both the pruned neutral and the full SNP panel to evaluate population differentiation at two organizational scales: sampling locations and post hoc K=5 population groupings (BEL-MEX, DRT-FLL-FLU, JAM-DOM, ARU-CUR, and PAN, Figure 1a,b). For sampling-location comparisons, individuals were downsampled to a maximum of four per location to reduce bias from uneven sample size, resulting in a total of 37 individuals retained for this analysis (sample IDs listed in Table S1.2). For the K=5 post hoc population comparisons, all 46 individuals were retained. Pairwise weighted $F_{ST}$ statistics (Weir & Cockerham 1984) were calculated using StAMPP (v1.6.3; (Pembleton et al. 2013) and confidence intervals (95%) were calculated using 100 bootstraps for the neutral SNP panel. Due to computational constraints, lower bootstrap values were used for the full SNP panel (n=3 and n=5 for sampling location and K=5 population runs, respectively) but were deemed sufficient due to highly confident p-values, reported as zero across all calculations.

To test for isolation-by-distance among sampling locations, we calculated linearized genetic distance $F_{ST}/(1-F_{ST})$ from pairwise $F_{ST}$ estimates described above. Geographic distance between sampling locations was calculated in R using pairwise great-circle distances based on the Haversine formula, implemented with distHaversine in the geosphere package v1.6-8 (Hijmans et al. 2017). The relationship between geographic and genetic distance was evaluated using a Mantel test implemented with mantel.rtest in the R package ade4 v1.7-24 (Dray & Dufour 2007), with 100,000 permutations used to assess significance. To further evaluate spatial patterns of genomic differentiation, Caribbean-wide migration rates were modeled for multiple deme sizes (300, 400, 500) with 5M MCMC iterations and a 1M burn-in period using Estimated Effective Migration Surfaces (EEMS) software (Petkova et al. 2016). The associated R package, rEEMSplots, was used to visualize model convergence and spatial patterns of migration.

#### Appendix S3. Demographic history, genetic diversity, and ROH methods.

##### Appendix S3.1. Detailed methods for PSMC and SMC++ demographic analyses.

Historical changes in effective population size were inferred using PSMC and SMC++. For both approaches, a mutation rate of $4e^{-9}$ bp/year and generation time of 5 years was used based on estimates from *Acropora millepora (Matz et al. 2018; Fuller et al. 2020)*. Pairwise SMC (PSMC) was run on consensus sequences of individual genomes generated in bcftools with the following default time-interval pattern: 4+25*2+4+6 (Li & Durbin 2011).

As PSMC can only model historical population sizes over 10,000 years ago, we additionally used the program SMC++ to model more recent demographic history (500-1M years ago) (Terhorst et al. 2017). First unphased SNPs were filtered for depth, quality, missingness, proximity to indels (5bp), and strand bias (MEAN(FMT/DP)>=10 && MEAN(FMT/DP)<=50 && QUAL>20 && GQ>20 && MQ>40 && F_MISSING<0.1 && SOR>4). Each chromosome from this filtered vcf was then converted into smc format using vcf2smc while masking uncalled, filtered, and repetitive regions. Replicate chromosome-level smc files were generated by varying the distinguished lineage to represent each sample. Effective population size was then modeled over time using 3000 thinning, 50 EM iterations, 40 knots, and a mutation rate of $4e^{-9}$. SMC++ was run for each population (K=5) separately as well as Caribbean-wide. One sample from the upper Florida Keys (CRF_Acer-099) was excluded due to its highly admixed ancestry profile in our population structure analyses.

##### Appendix S3.2. Detailed methods for genetic diversity versus genetic uniqueness analysis.

We followed the theoretical and methodological approach of Weeks et al. (2016), who credit (Coleman et al. 2013), to test the hypothesis that threatened species with increasingly small and fragmented populations will exhibit a negative relationship between genetic diversity levels (expected heterozygosity, $H_e$) and “uniqueness” among populations (population-specific $F_{ST}$) due to the increasing effects of genetic drift (Weeks et al. 2016).

VCF genotypes from the full SNP panel (n=37 downsampling as in pairwise $F_{ST}$ calculations, see Appendix S2.1) were imported with vcfR (Knaus & Grünwald 2017), converted to a diploid genlight object using adegenet (Jombart et al. 2010), and assigned population group according to sampling location or K=5 subpopulation assignment. Following conversion of genotypes into the allele-dosage format, population-specific $F_{ST}$ was estimated using hierfstat::fs.dosage (Goudet 2005). For expected heterozygosity ($H_e$), allele frequencies were calculated across loci and locus-specific $H_e$ was estimated as $2p(1-p)$, where $p$ is the allele frequency, and reported as mean $H_e$ across loci for each location.

##### Appendix S3.3. Detailed methods for ROH inbreeding risk analysis.

Length and distributions of Runs of Homozygosity (ROH) across individuals were detected using GARLIC v.1.1.6a (Szpiech et al. 2017). GARLIC implements the ROH calling method of Pemberton et al. (2012) and Blant et al. (2017) (Pemberton et al. 2012; Blant et al. 2017), while also incorporating a length-based classification module to identify different biological classes of ROH. GARLIC was run with an error rate of 0.001, and auto-detection of best window size.  Best practices for generating locations of centromeres for running GARLIC in non-model organisms were adapted from (Szpiech et al. 2017).

Further filtering and analyses of ROH were performed in R v. 4.5.2. Minimum Length of ROH (kb) was set 10 kb to exclude ancient population processes. To identify what length of ROH (kb) likely still harbors deleterious variation and is therefore functionally relevant in the existing population, runs of homozygosity were timed to their last common ancestor. Previous studies show that longer ROH contain more deleterious alleles (Szpiech et al. 2013), and theory suggests that most deleterious alleles will segregate after 25 generations such that two individuals that may breed will not harbor the same copy of a deleterious mutation (Kimura & Ota 1969; Kyriazis et al. 2025), so only ROH with a LCA of 25 generations or less is likely to contain deleterious variation in the homozygous state. Recombination rate was calculated using the maps generated from (Locatelli et al. 2024) adapted to the *A. cervicornis* genome used in this study (GenBank accession no. GCA_964034985.1). The expected number of generations to a common ancestor (g) for an ROH of a given length (kb) was calculated using the following equation:

$$g=100/2rL$$

Where r is the recombination rate, and L is the length of ROH in kbp. ROH thresholds for capturing the functional effects of inbreeding were set using the assumption that recessive deleterious segregate for no more than 25 generations. IDrisk was calculated according to Kyriazis et al. 2025 (Kyriazis et al. 2025):

$$IDrisk=FROH \times \pi non\text{-}ROH$$

Where $F_{ROH}$ is calculated as the fraction of the genome spanned by functionally-long ROH, and πnon-ROH is the heterozygosity in non-ROH regions per kb. Heterozygosity in non-ROH regions of the genome was calculated using plink2 (--het, plink2/2.00a3.7LM) (Chang et al. 2015).

*Rationale for ROH Length Choices*

Very few studies have investigated the presence and consequences of runs of homozygosity in reef-building corals. In systems where it is better studied such as humans and mammals, the length of ROH associated with functional consequences of inbreeding are much longer than any runs identified in corals. The minimum length of ROH (kb) that represents common ancestors 25 generations or sooner was ~228kbp, However, some studies have suggested in species with larger genomes that ROH > 100kbp may still contain deleterious variation, so further analyses were performed with both ROH > 200kbp and > 100kbp.

### Appendix S4. Adaptive scans methods.

**Appendix S4.1. Additional methods associated with adaptive variation scans.**

To identify genes putatively under selection, we performed SNP and window-based analyses at both range-wide and subpopulation scales. Range-wide analyses treated all samples as a single metapopulation (K=1) to identify genomic regions with evidence of selection across the Caribbean. Subpopulation analyses used K=5 post hoc groupings to identify outlier loci or genomic regions diverging between subpopulations, indicative of local adaptation.

*Metapopulation (K=1) Scale*

Genomic regions which could be under selection within the Caribbean-wide metapopulation were evaluated by recalculating pi and Tajima's D in smaller, 10kb windows using the filters and methods described in the genomic diversity section above. Windows with less than 5kb callable sites were removed. Candidate outlier windows were identified as those within the top or bottom 0.01% of the genome-wide pi distribution that also exhibited signatures of balancing or purifying selection (Tajima's D >2 or <-2 respectively). Diversity statistics within 10kb outlier regions were also calculated at 1kb intervals and annotated to identify putative genes under selection. To determine whether balancing selection was being driven by local adaptation, sample-specific genotypes were extracted for outlier genes and clustering by sampling location and/or population was visually assessed using the program pheatmap (Kolde 2025) in R.

*Subpopulation (K=5) Scale*

Using the K=5 post hoc population assignments (BEL-MEX, DRT-FLL-FLU, JAM-DOM, ARU-CUR, and PAN), we calculated population-specific nucleotide diversity ($\pi$) and pairwise $F_{ST}$ to generate meadow plots following Duffin et al. (in press). Meadow plots were used to visualize putative selective sweeps within regions by jointly identifying elevated population differentiation and strong shifts in nucleotide diversity between population groupings. Conceptually, these plots extend a Manhattan-style genome scan by plotting $F_{ST}$ across chromosomes and using $\pi$-ratio values to indicate which population shows reduced diversity within each outlier region.

Pixy v2.0.0 (Korunes & Samuk 2021) was used to calculate $\pi$ and pairwise $F_{ST}$ from variant and invariant loci in non-overlapping 10 kb windows. Windows with less than 5 kb of data per 10 kb window were excluded. Pixy outputs were imported into R and used to calculate pairwise $log_2$-transformed $\pi$ ratios for all 10 contrasts among the five K=5 post hoc population groupings. Negative $F_{ST}$ estimates were excluded before defining outlier windows.

For each pairwise comparison independently, candidate outlier windows were defined as windows in the top 1% of the empirical $F_{ST}$ distribution that also fell within either the lower or upper 1% of the empirical $log_2(\pi\text{-ratio})$ distribution. Meadow plots were generated in R to display windowed $F_{ST}$ across chromosomes, with point color representing the direction and magnitude of $log_2(\pi\text{-ratio})$ and enlarged points indicating candidate outlier windows.

To summarize the extent of outlier signals across meadow plots, we identified contiguous outlier peaks within each chromosome and pairwise comparison. Candidate peaks were permitted to span up to three consecutive non-outlier windows. Final peak length was reported as the number of outlier 10 kb windows within each region, excluding intervening non-outlier windows and windows removed because of insufficient data. These peak counts were then visualized as violin plots to compare the distribution of outlier-region lengths across pairwise K=5 contrasts.

We additionally used three SNP-based approaches to identify outlier loci potentially under regional selection: PCA-based PCAdapt v.4.0 (Privé et al. 2020), and $F_{ST}$-based methods BayeScan v.2.1 (Duforet-Frebourg et al. 2014) and OutFLANK v. 0.2 (Whitlock & Lotterhos 2015). In all methods SNPs required a p-value < 0.01 to be identified as outliers. OutFLANK was run with a minimum heterozygosity of 0.1 and a False discovery rate (FDR) of 1%. Both OutFLANK and Bayescan were run using pairwise comparisons using the K=5 post hoc population assignments.

#### Appendix S5. Symbiont composition methods.

##### Appendix S5.1. qPCR for symbiont genera identification.

To inform symbiont genomic analysis, symbiont communities were first screened via qPCR to identify which symbiont genera were present in samples. *Symbiodinium* was quantified in singleplex using pre-designed primers and probes specific to the *Symbiodinium* actin gene (Aact For: 5'–ATGAAGTGCGACGTGGACAT – 3', Aact Rev: 5'–GGAGGACAGGATGGAGCC T–3', Aact Probe: 5'-FAM-CGTTGGAGTAGAGGTC-MGB-3') (Palacio-Castro et al. 2021). *Cladocopium* and *Durusdinium* were quantified in multiplex following (Cunning & Baker 2013) using genera-specific primers and probes (Cact For: 5'-CCAGGTGCGATGTCGATATTC-3', Cact Rev: 5'-TGGTCATTCGCTCACCAATG-3', Cact Probe: 5'-VIC-AGGATCTCTATGCCAACG-MGB-3', Dact For: 5'-GGCATGGGGTAAGCACTTCTT-3', Dact Rev: 5'-GATCCTTGAACTAGCCTTGGAAAC-3', Dact Probe: 5'-FAM-CAAGAACGATACCGCC-MGB-3'). All reaction volumes were 25ul total, including 12.5 ul of Brilliant multiplex qPCR Master Mix, 0.375 ul of reference dye, and 2 ul of genomic DNA (20 ng total). Reactions were run in duplicate using the following cycling protocol (10min @ 95C {15s @ 95C ; 1min @ 60C ; 30s @ 72C} x 40) on the AriaMx system to determine Ct values. Ct values <38 were used to detect the presence of symbiont genera.

##### Appendix S5.2. Additional methods associated with symbiont screening.

Symbiont genera were initially screened via qPCR following (Cunning & Baker 2013; Palacio-Castro et al. 2021) using probe and primer sets specific to *Symbiodinium*, *Cladocopium*, and *Durusdinium* to confirm *A. cervicornis'* predominant association with *Symbiodinium fitti* (Reich et al. 2021). Probe and primer sets are provided in Appendix S5.1.

To evaluate finer-scale symbiont variation, reads from high coverage samples mapping to the *S. fitti* genome were parsed to characterize genetic variation within and across individuals. As individual samples could host more than one symbiont strain, allele frequencies were estimated from putative sample pools using Freebayes (–pooled-continuous) with a ploidy of 1 (Garrison & Marth 2012). Reads were filtered to retain alignments with a minimum mapping quality >40 and base quality >20 and a maximum of 4 SNPs with a depth of 2-200x were retained per sample per locus. Loci were filtered for a minimum depth of 10x in at least 90% of samples and GATK's VariantsToTable function was used to extract major haplotype and allele frequency information. Major haplotypes were treated as proxies for dominant symbiont strain, and allele frequencies were used as estimates of strain abundance following (Breusing et al. 2022) and analyzed across populations using principal component analyses in R.

To compare genome-wide symbiont results with ITS2-based studies, we additionally blasted aligned symbiont reads against the SymPortal database (Hume et al. 2019). Blast hits with a length of at least 149bp and 99% similarity were retained. High-quality hits were tallied for each ITS2 variant and the relative abundance in each sample was visualized using stacked bar plots in R. Differences in community composition at the ITS2, allele frequency, and dominant haplotype level were evaluated across sampling locations using PERMANOVAs from the R package vegan (Dixon 2003).

### Appendix S6. Haplotype reference panel methods.

#### Appendix S6.1. Additional methods associated with construction and validation of the *A. cervicornis* haplotype reference panel resource.

*Haplotype Reference Panel Construction*

The *A. cervicornis* haplotype reference panel was generated using a two-step phasing workflow. We first applied read-backed phasing within individuals, then used population-backed statistical phasing to refine haplotypes across the full panel. The starting SNP dataset was filtered to retain high-confidence variants with INFO/DP > 10, QUAL > 20, GQ > 20, MQ > 40, FS < 10, AC > 3, no missing genotypes, and SOR < 4. Multiallelic sites were decomposed into biallelic records using bcftools norm, and PL fields were converted to GL format using bcftools +tag2tag (bcftools v1.19; Danecek et al. 2021). The resulting BCF was split by individual and indexed for read-backed phasing.

Read-backed phasing was performed separately for each sample using WhatsHap v2.4 (Martin et al. 2016). Each run used the sample-specific BCF, the corresponding sorted BAM file, and the combined *A. cervicornis/S. fitti* reference genome (GCA_964034985.1_Acer_S.fitti.fa). Individual WhatsHap-phased VCFs were then merged into a multi-sample VCF (bcftools merge) and standardized variants IDs were set.

Before the second phasing step, we filtered the panel (at this stage, n=46) to identify and remove individuals likely to reduce haplotype quality because of strong admixture or population structure artifacts. For this check, SNPs destined for the admixture analysis were filtered to retain biallelic non-indel variants with MAC ≥ 3, thinned to one site per 20 kb, and LD-pruned in plink2 (Gaunt et al. 2007; Chang et al. 2015) using a 500 kb window and $r^2$ threshold of 0.2. ADMIXTURE v1.3.0 (Alexander et al. 2009) was run across K = 1-10 with 20 random-seed runs and 10-fold cross-validation for each K. The resulting ancestry estimates were used to identify individuals with strongly mixed assignments. After removing one individual collected from FLU (CRF_Acer-099) and another from PAN (SRR24007608) due to admixture, the final haplotype panel retained 44 individuals (Panama, n=9; Belize, Dry Tortugas, Jamaica: n=5 each; Dominican Republic, Curaçao, Aruba: n=4 each; Lower Florida Keys, Upper Florida Keys: n=3 each; Mexico, n=2).

To identify loci for the final reference panel, we identified and removed SNPs deviating from Hardy-Weinberg equilibrium while accounting for population structure using RUTH (v3.29.4, Kwong et al. 2021) and PCA eigenvectors generated from the LD-pruned n=44 dataset (Figure S1.1). First, a RUTH input vcf was generated from the merged WhatsHap-phased dataset by retaining biallelic SNPs with no missing data, excluding repeat-masked regions, recalculating allele-count tags, and removing monomorphic sites. RUTH was then run in site-only mode using PL genotype likelihoods. Positions passing the RUTH-based HWE filter (−5 < HWE_SLP_I < 5) and minor allele count threshold (MAC > 3) were extracted and used on the WhatsHap-phased VCF to produce a filtered n=44 VCF ready for population-backed phasing.

For the second phasing step, the filtered n=44 VCF was split by scaffold across the 15 retained scaffolds, then phased with Beagle 5.5 (beagle.27Feb25.75f.jar; Browning & Browning

2007) using scaffold-specific genetic maps, gp=true, and ne=10000. Beagle-phased scaffolds were concatenated together to produce the final phased *A. cervicornis* haplotype reference panel.

*Haplotype Reference Panel Validation*

To evaluate imputation accuracy, we used five *A. cervicornis* samples that had been independently sequenced at both high and low coverage (hereafter abbreviated “HiCo” and “LoCo”, respectively). These validation samples were all from the Florida region, including three FLL (Lower Florida Keys) samples, one FLU (Upper Florida Keys) sample, and one DRT (Dry Tortugas) sample. HiCo genotypes were treated as the truth set, while the corresponding LoCo datasets were processed through the imputation pipeline and compared back to the HiCo calls. To better test the reference panel’s ability to impute genotypes in new *A. cervicornis* samples, each validation sample was imputed using a leave-one-out reference panel specific to that individual.

Genotypes were called for LoCo samples with bcftools mpileup and bcftools call using the combined *A. cervicornis/S. fitti* reference genome (GCA_964034985.1_Acer_S.fitti.fa). Input VCFs were split by scaffold, normalized to the reference genome, restricted to biallelic SNPs, assigned standardized SNP IDs, and updated with allele-count and allele-frequency tags. Imputation was performed in two rounds. First, LoCo genotype likelihoods were imputed with Beagle 4.1 (beagle.27Jan18.7e1.jar; Browning & Browning 2007) using the leave-one-out haplotype reference panel in bref format and the *A. cervicornis* genetic map. Genotype probabilities were retained from this first round, and low-confidence calls were removed by setting genotypes with GP<0.99 to missing (./). This GP-filtered dataset represented the recalibrated, or adjusted, LoCo genotype set. A second imputation round was then run with Beagle 5.5 (beagle.27Feb25.75f.jar; Browning et al. 2021) using filtered genotypes and the leave-one-out reference panel in bref3 format. Final imputed genotypes were filtered using Beagle’s dosage $R^2$ metric “DR2,” retaining sites with DR2≥0.99.

Imputation accuracy was determined by comparing three LoCo-derived, or “test” genotype sets to the matching HiCo truth set: raw LoCo calls, adjusted calls (post-GP<0.99 filter), and newly imputed calls. For each comparison, we identified overlapping sites between the HiCo and LoCo-derived VCFs, extracted genotype calls, and calculated overall genotype concordance as the proportion of matching genotypes. Concordance was also summarized separately for 0/0, 0/1, and 1/1 genotypes to evaluate whether accuracy differed among genotype classes. Finally, genotype dosage accuracy was evaluated using global Pearson $R^2$ between HiCo “truth” and LoCo “test” genotype dosages. Reported values were averaged across the five validation samples.

### Appendix S7. Population structure results.

**Appendix S7.1. K-selection criteria across population structure analyses.**

We evaluated clustering solutions across four complementary inference approaches: DAPC (Figure S7.4), ADMIXTURE (Figure S7.6, Figure S7.7), sNMF (Figure S7.10, Figure S7.11), and PopCluster (Figure S7.14). Clustering solutions were evaluated using method-specific K-selection criteria, including CV error (Figure S7.8), cross-entropy (Figure S7.13), BIC (Figure S7.5), Evanno ΔK (Figure S7.9, Figure S7.12, Appendix S7.4), $F_{ST}/F_{IS}$ (Figure S7.15), and DLK2 (Figure S7.16), and summarized in Figure 1d. Although all analyses were run across values of K up to 10, representing up to 10 genetic clusters or "K" subpopulations, only three of the eight K-selection approaches meaningfully evaluated K=1: ADMIXTURE based on lowest CV error (Figure S7.8), sNMF based on lowest cross-entropy (Figure S7.13), and PopCluster based on highest $F_{ST}/F_{IS}$ (Figure S7.15). Each of these identified K=1 as the best-supported solution (Figure 1d), consistent with detectable gene flow and shared ancestry across the species' sampled range.

The remaining five approaches were restricted to evaluating K≥2, either because the inference method itself requires multiple groups to discriminate among clusters, as in DAPC, or because the downstream K-selection criterion excludes K=1, as in Evanno ΔK and DLK2 (Figure 1d). Among these K≥2 approaches, four of five identified K=5 as the most likely number of populations: DAPC based on lowest BIC score (Figure S7.4, Figure S7.5) and ADMIXTURE, sNMF, and PopCluster based on Evanno ΔK (respectively: Figure 1d; Figure S7.6, Figure S7.7, Figure S7.9; Figure S7.10, Figure S7.11, Figure S7.12; Figure S7.14, Appendix S7.4). The final approach favored a different solution: DLK2 selected K=6 for the PopCluster results, though it only marginally outperformed K=5 (Figure 1d; Figure S7.14, Figure S7.16).

Together, these results indicate that connectivity represents a dominant population structure signal in the neutral SNP panel, alongside moderate regional structure that becomes apparent when analyses are restricted to K≥2. We therefore interpret K=1 as the dominant ancestry signal but use K=5 to test whether regional structure reflects longer-term evolutionary divergence using downstream genomic approaches.

**Appendix S7.2. Jamaica and Dominican Republic ancestry coefficients at K=5.**

Although Jamaica and Dominican Republic individuals clustered together in the K=5 sNMF solution, their ancestry-coefficient profiles differed in both the magnitude and composition of secondary ancestry. Jamaica showed the strongest evidence of mixed ancestry among sampled locations, with 25.25% ± 5.14% of ancestry assigned outside its dominant JAM-DOM cluster on average (Table S7.1). Secondary ancestry in Jamaica was distributed across PAN (11.22% ± 1.39%), BEL-MEX (6.83% ± 4.57%), and DRT-FLL-FLU (6.53% ± 5.55%) dominant clusters, with only minimal assignment to ARU-CUR (0.67% ± 0.81%) (Figure 1a; Table S7.1).

In contrast, Dominican Republic individuals showed stronger assignment to the JAM-DOM cluster and lower secondary ancestry overall. Two Dominican Republic individuals had >99.9% assignment to JAM-DOM, and one had 91.65% assignment, with no secondary cluster exceeding 4% in that individual. The final Dominican Republic individual had 84.73% assignment to JAM-DOM and showed a larger secondary assignment to the geographically closer ARU-CUR cluster (11.35%), while assignment to the remaining, more geographically distant clusters remained low (DRT-FLL-FLU: 3.38%; PAN: 0.53%; BEL-MEX: 0.01%) (Figure 1a; Table S7.1).

Together, these ancestry coefficients indicate that the K=5 JAM-DOM cluster captures shared ancestry between Jamaica and Dominican Republic but also masks differences between the two locations. Jamaica showed broader mixed ancestry across multiple regional clusters, whereas Dominican Republic individuals were generally more strongly assigned to the shared JAM-DOM cluster, with secondary ancestry either low or primarily directed toward ARU-CUR in one individual.

**Appendix S7.3. Alternative K=6 ancestry-clustering solution.**

At the finer K=6 resolution selected by DLK2 for PopCluster (Figure 1d), the distinct PAN, ARU/CUR, and BEL/MEX clusters remained intact, including the same FLU individual assigned to BEL/MEX (Figure S7.14, Figure S7.16). This result suggests that the K=5 solution captures the main regional structure without obscuring the broader K=1 signal of substantial connectivity.

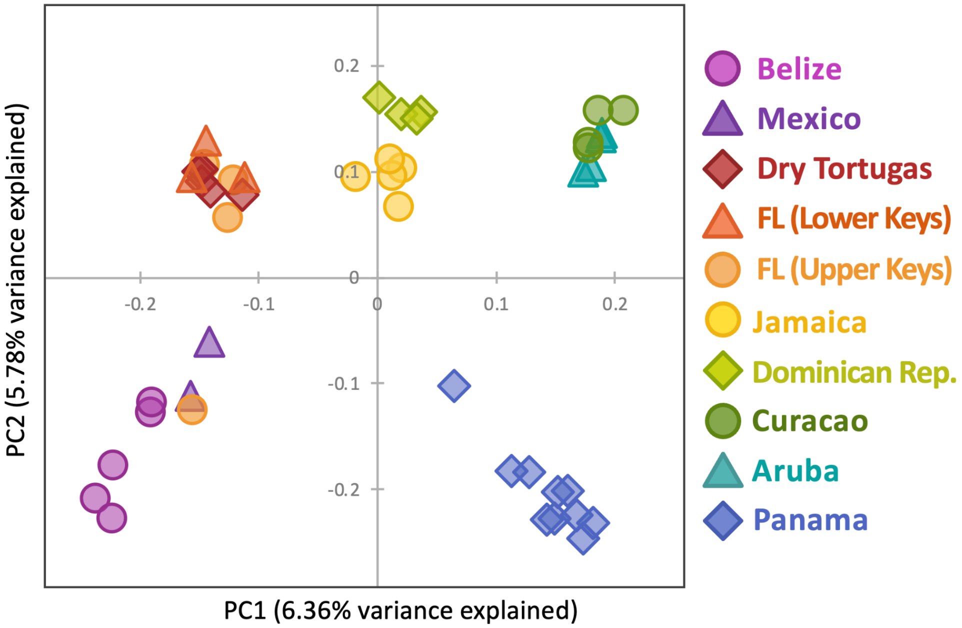

**Figure S7.1. PCA results for full SNP panel** (n= 2,527,725 SNPs).

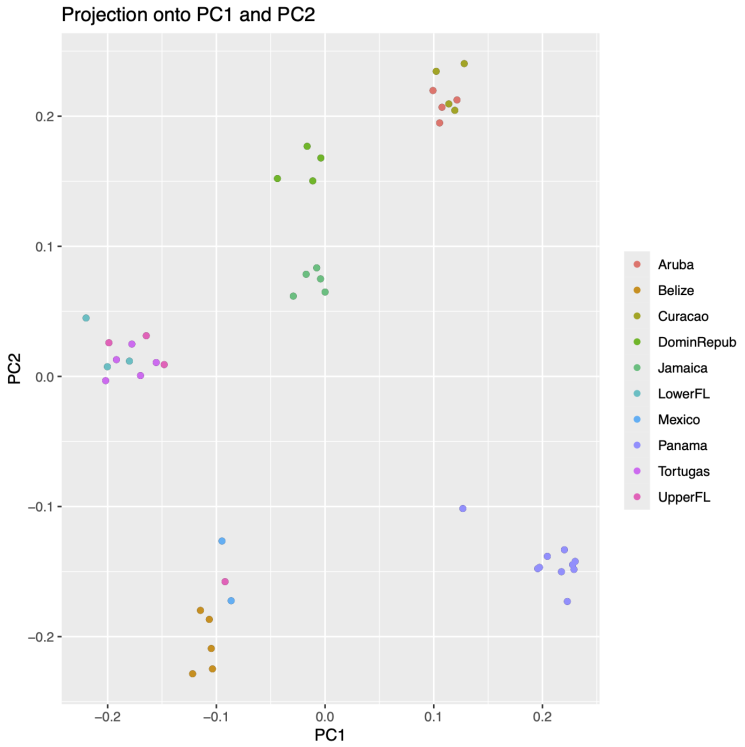

**Figure S7.2. Standard raw PCA plot output on the neutral SNP panel** (n= 177,516 SNPs).

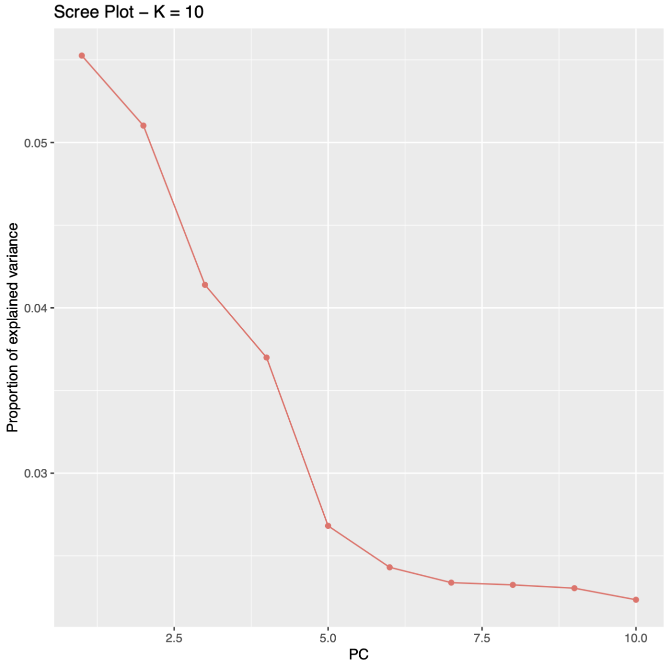

**Figure S7.3. Scree plot associated with PCA on the neutral SNP panel** (n= 177,516 SNPs).

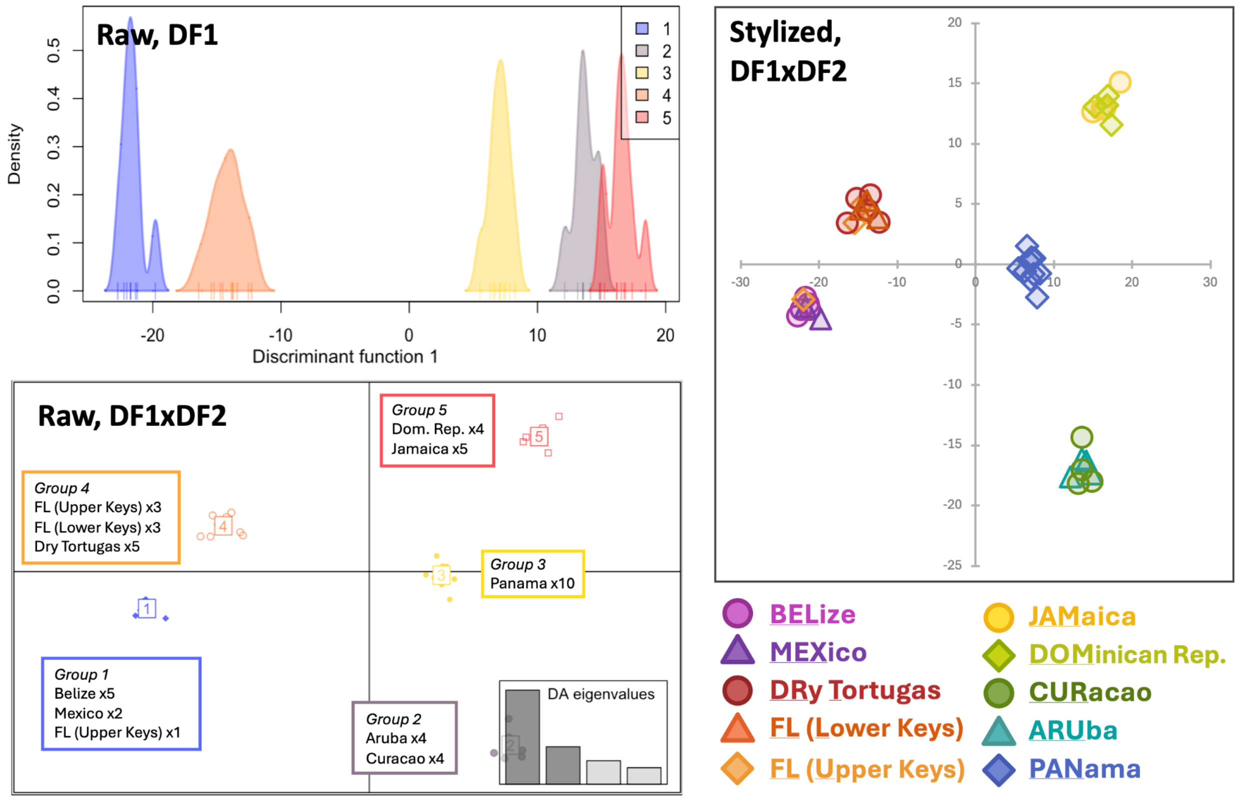

**Figure S7.4. DAPC plot results** on the neutral SNP panel (n= 177,516 SNPs).

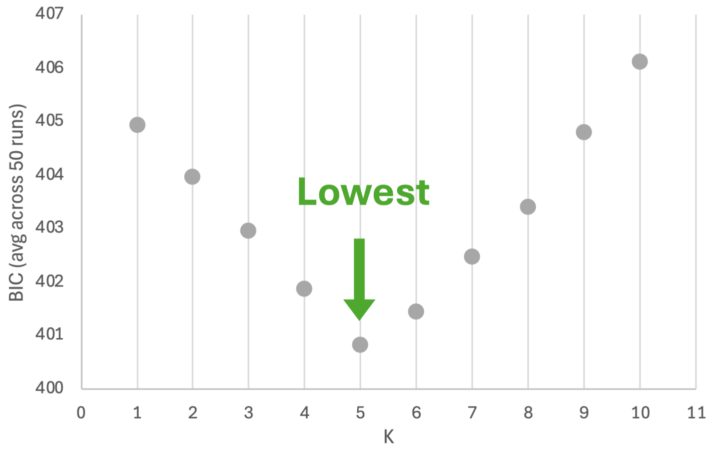

**Figure S7.5. Aggregated BIC scores across runs of DAPC** (n=50 for each of K=1 through K=10) on the neutral SNP panel (n= 177,516 SNPs).

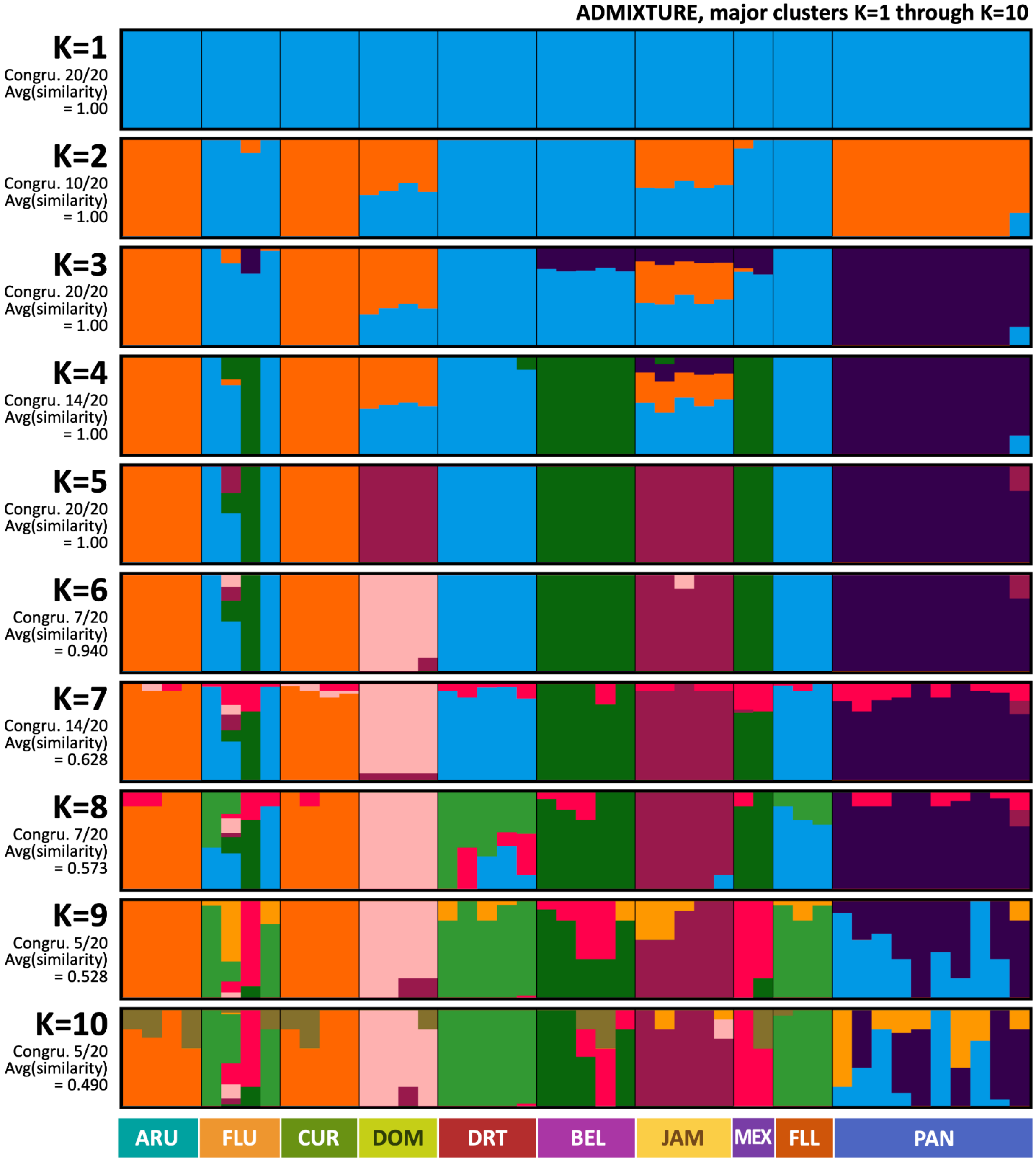

**Figure S7.6. ADMIXTURE results, displaying major modes** identified through CLUMPAK (https://clumpak.tau.ac.il/) summation of 20 random seed runs for each value of K (K=1 through K=10) on the neutral SNP panel (n= 177,516 SNPs).

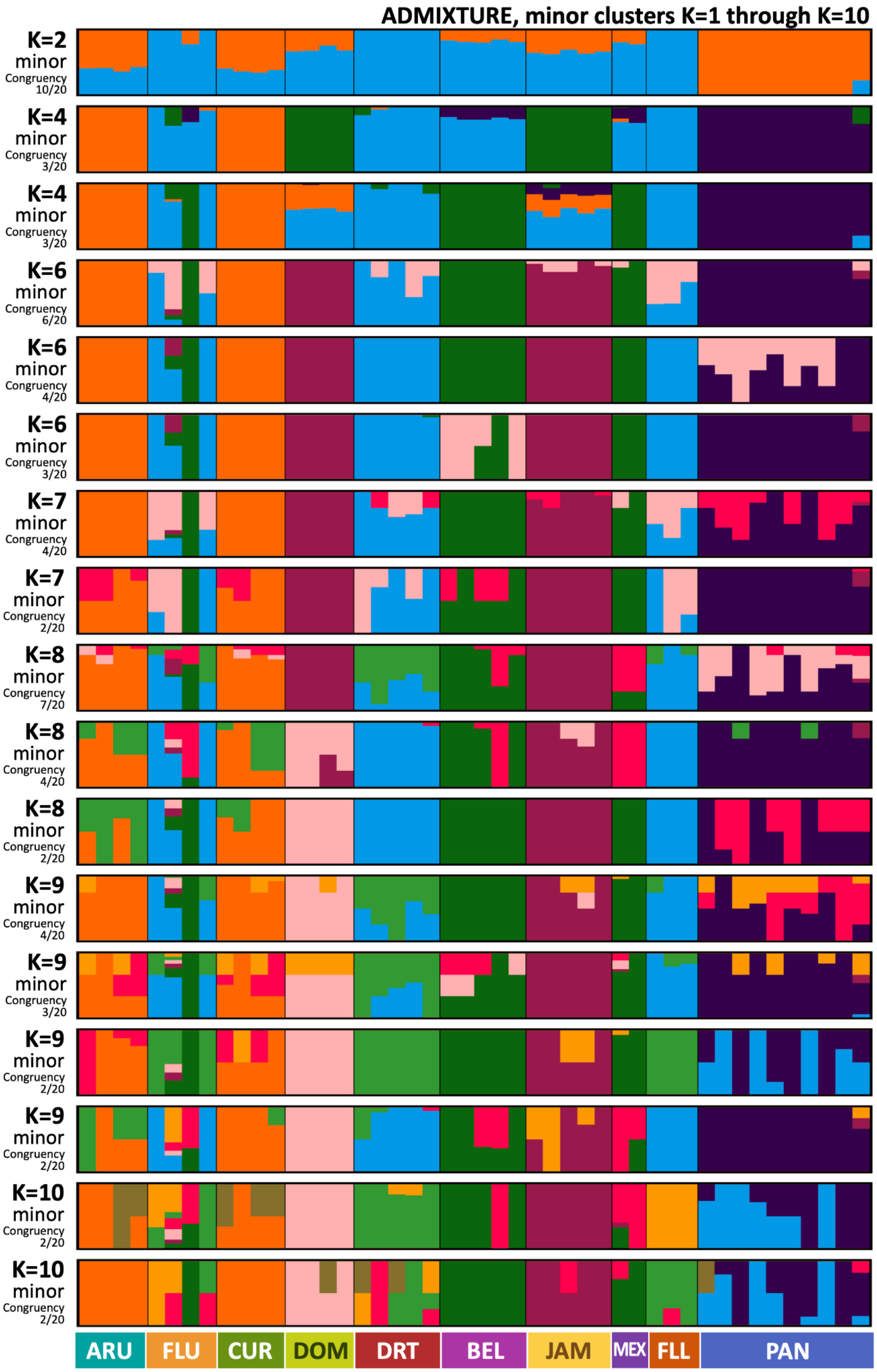

**Figure S7.7. ADMIXTURE results, displaying minor modes** identified through CLUMPAK (https://clumpak.tau.ac.il/) summation of 20 random seed runs for each value of K (K=1 through K=10) on the neutral SNP panel (n= 177,516 SNPs).

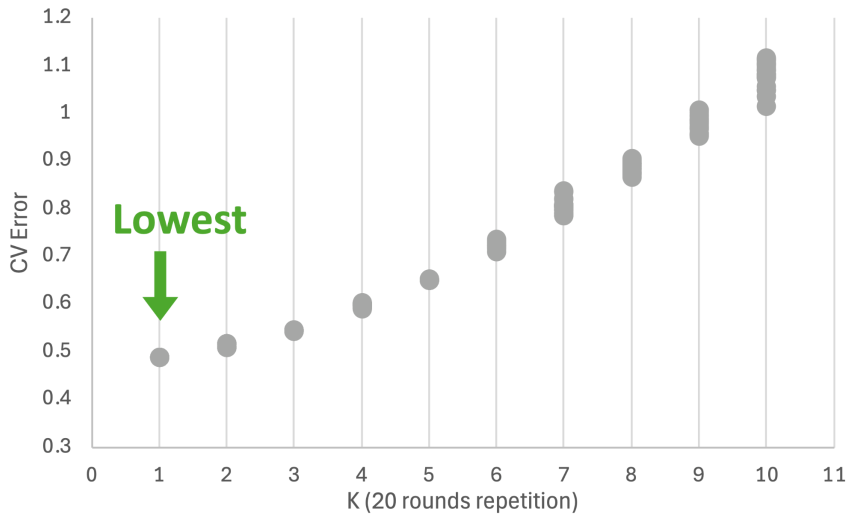

**Figure S7.8. ADMIXTURE cross-validation (CV) error statistic output** on the neutral SNP panel (n= 177,516 SNPs).

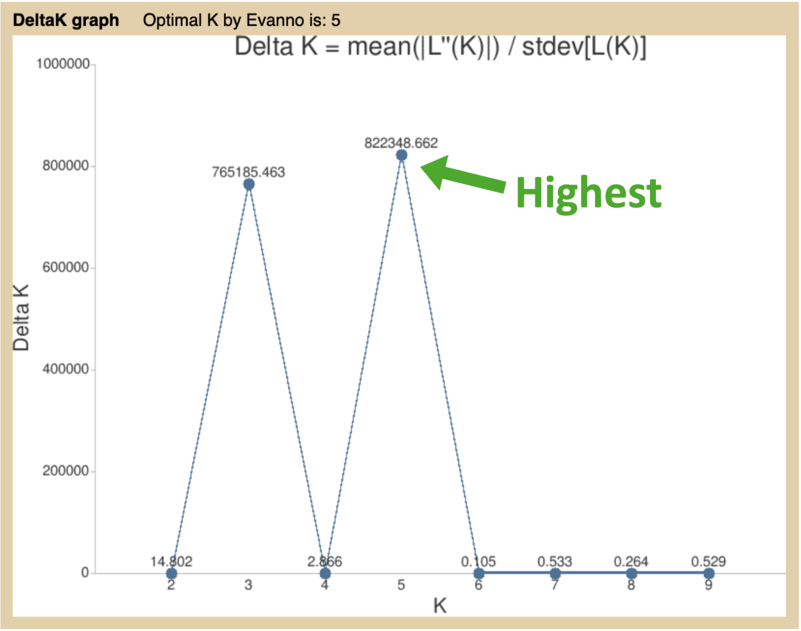

**Figure S7.9. CLUMPAK BestK output using the ΔK** (Evanno et al. 2005) **method, run on ADMIXTURE** loglikelihood statistics generated from the neutral SNP panel (n= 177,516 SNPs).

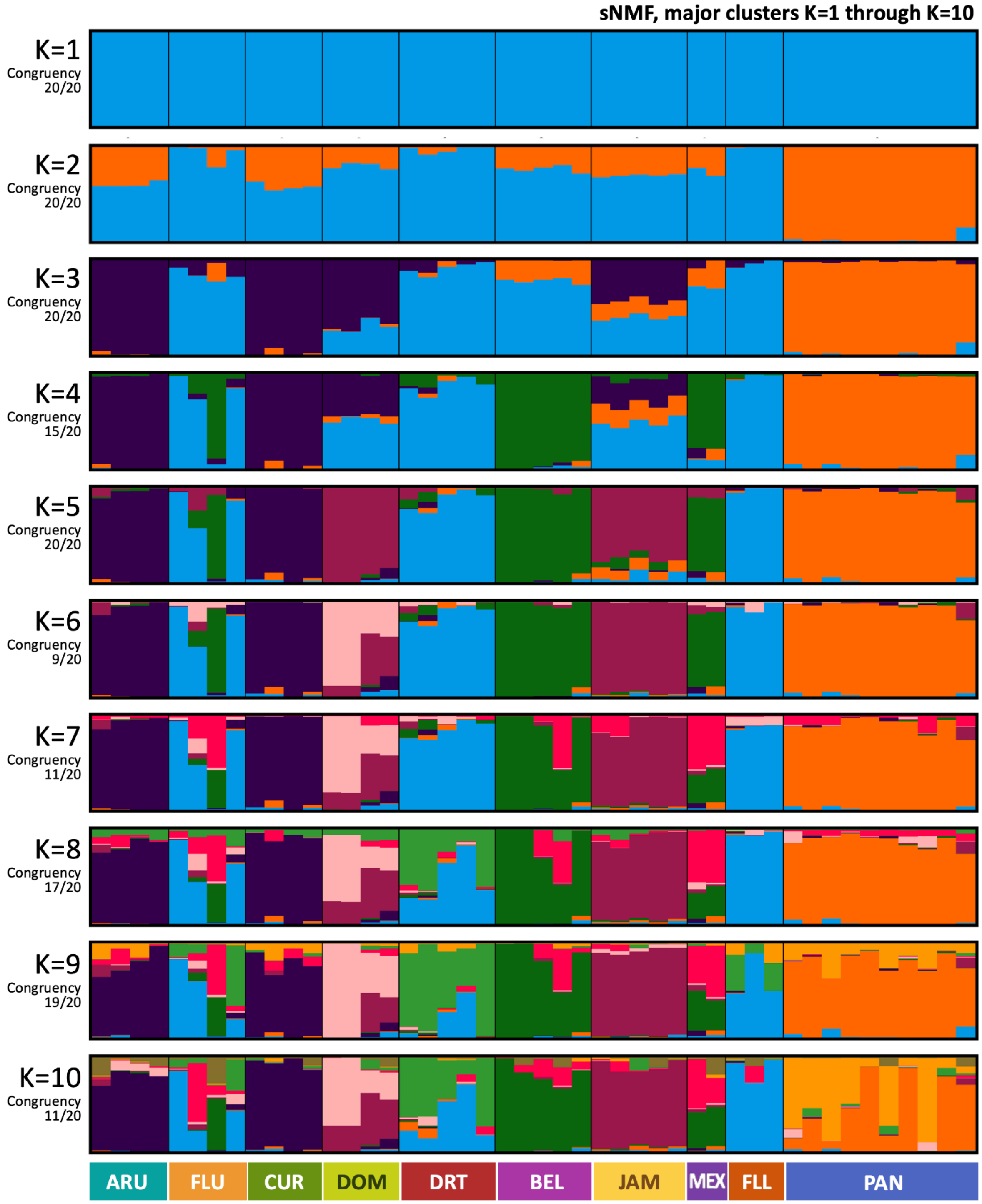

**Figure S7.10. sNMF results, displaying major modes** identified through CLUMPAK (https://clumpak.tau.ac.il/) summation of 20 random seed runs for each value of K (K=1 through K=10) on the neutral SNP panel (n= 177,516 SNPs).

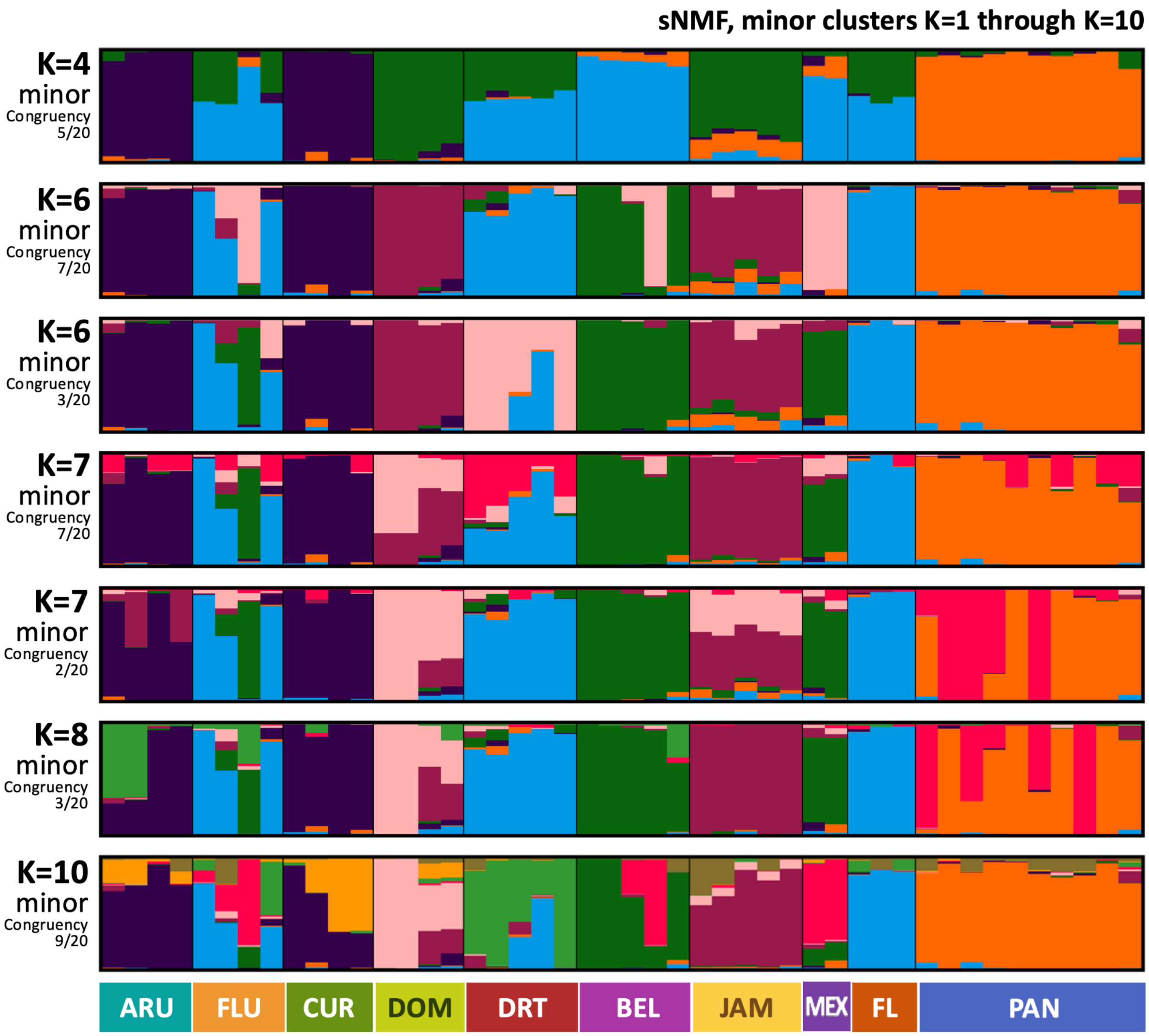

**Figure S7.11. sNMF results, displaying minor modes** identified through CLUMPAK (https://clumpak.tau.ac.il/) summation of 20 random seed runs for each value of K (K=1 through K=10) on the neutral SNP panel (n= 177,516 SNPs).

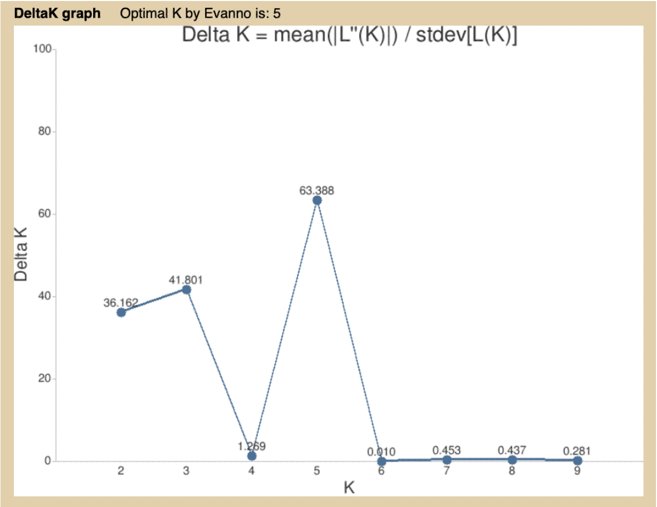

**Figure S7.12. CLUMPAK BestK output using the ΔK** (Evanno et al. 2005) **method**, **run on sNMF** loglikelihood statistics generated from the neutral SNP panel (n= 177,516 SNPs).

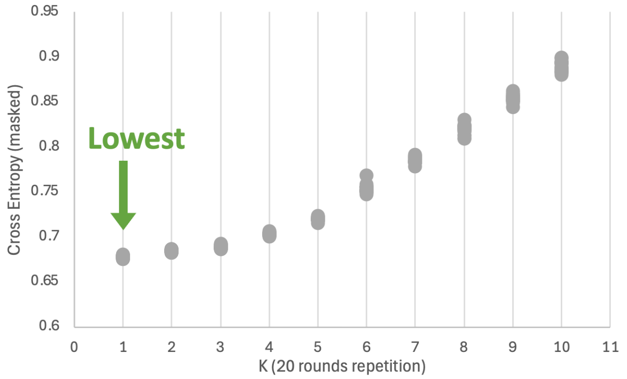

**Figure S7.13. sNMF cross-entropy criteria values** derived from predicted masked genotypes from the neutral SNP panel (n= 177,516 SNPs).

**Table S7.1. Average percent admixture** (1 - [major assignment prop.]) **from sNMF results**, summed using CLUMPAK (20 runs) for K=5 on the neutral SNP panel (n= 177,516 SNPs).

<table>
<tr><th rowspan="2"></th><th colspan="2">sampling location</th><th colspan="2">K=5 population</th></tr>
<tr><th>mean</th><th>SD</th><th>mean</th><th>SD</th></tr>
<tr><td>BEL</td><td>0.038</td><td>0.042</td><td rowspan="2">0.089</td><td rowspan="2">0.093</td></tr>
<tr><td>MEX</td><td>0.216</td><td>0.003</td></tr>
<tr><td>DRT</td><td>0.129</td><td>0.107</td><td rowspan="3">0.117</td><td rowspan="3">0.128</td></tr>
<tr><td>FLL</td><td>0.016</td><td>0.027</td></tr>
<tr><td>FLU</td><td>0.178</td><td>0.169</td></tr>
<tr><td>JAM</td><td>0.252</td><td>0.051</td><td rowspan="2">0.167</td><td rowspan="2">0.117</td></tr>
<tr><td>DOM</td><td>0.059</td><td>0.074</td></tr>
<tr><td>CUR</td><td>0.046</td><td>0.043</td><td rowspan="2">0.049</td><td rowspan="2">0.049</td></tr>
<tr><td>ARU</td><td>0.053</td><td>0.061</td></tr>
<tr><td>PAN</td><td>0.053</td><td>0.057</td><td>0.053</td><td>0.057</td></tr>
</table>

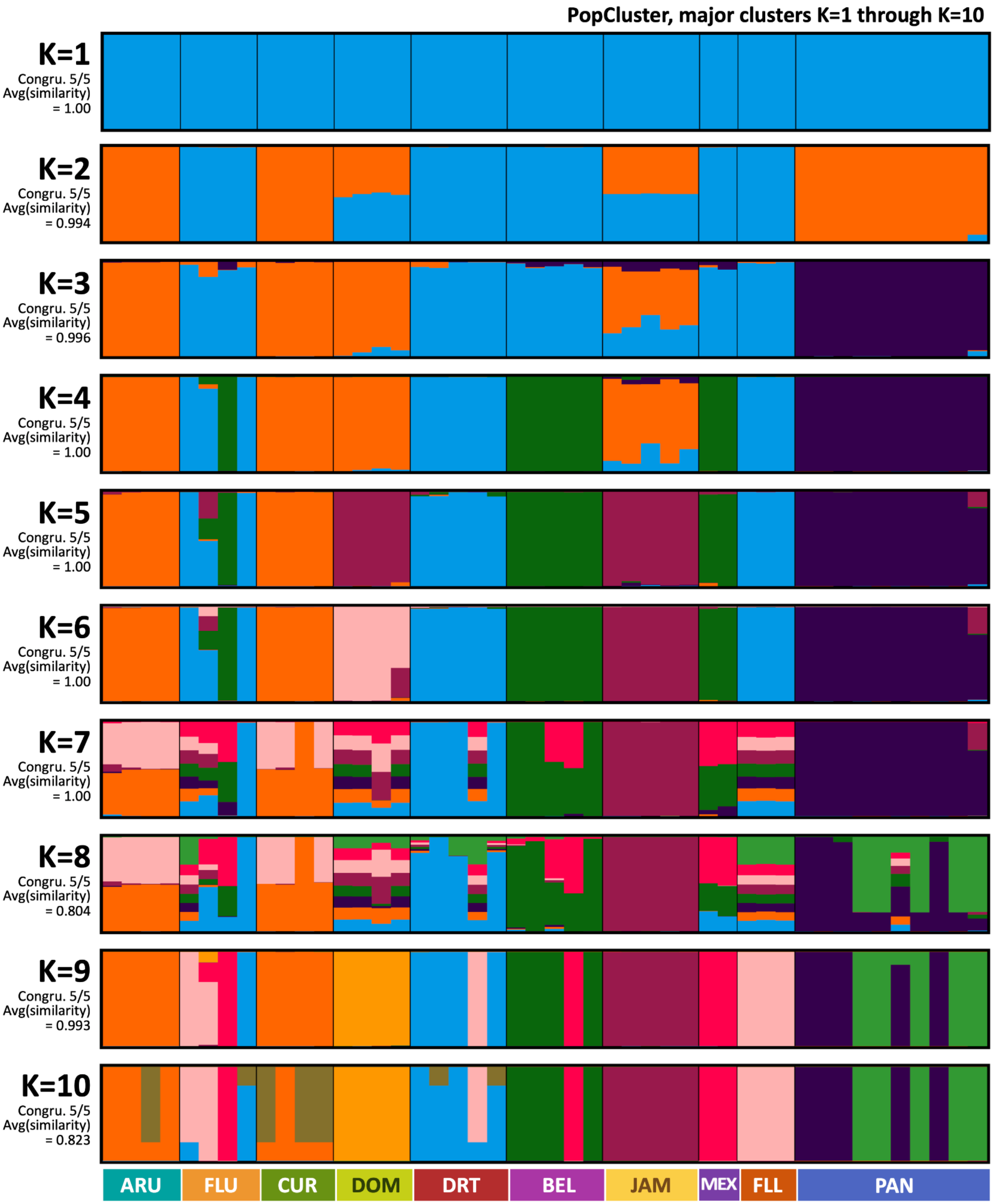

**Figure S7.14. PopCluster** (Wang 2025) **results, displaying major modes** identified through CLUMPAK (https://clumpak.tau.ac.il/) summation of 5 random seed runs for each value of K (K=1 through K=10) on the neutral SNP panel (n= 177,516 SNPs).

**Appendix S7.4. Log-likelihood adjustments for compatibility with Evanno et al. (2005) K-selection method**

**Issue:** Across 2 batches of n=5 runs of PopCluster (Wang 2025), multiple values of K produced log-likelihood (Ln) values with no deviation across runs. Evanno's 2005 K-selection method relies on standard deviation (SD) values >0 to calculate its best K metric (ΔK) with the formula:

$$\Delta K = |Ln''(K)| / SD[LnP(K)]$$

To facilitate interpretation of PopCluster results with the Evanno et al. (2005) method, the following data correction options were explored:

Data Correction 1: Omit problematic K value (K=9).

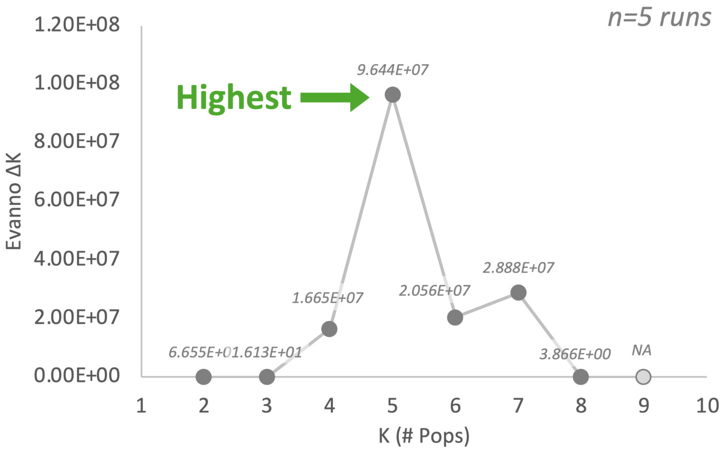

**Appendix S7.4.a (figure). CLUMPAK BestK output** using the ΔK (Evanno et al. 2005) method, run on PopCluster log-likelihood statistics (Wang 2025) generated from the neutral SNP panel (n= 177,516 SNPs) **using data correction method 1**, where the problematic K value with no SD(K=9) is omitted to facilitate the Evanno method-based calculation.

Data Correction 2: Add 1 to all standard deviation (SD) measurements before ΔK calculation.

*Rationale: A value of 1 in the denominator minimizes over- or under- inflation of the numerator as a byproduct of the correction step. While this is an imperfect correction under some hypothetical scenarios, biases that might impact Correction 2 would behave differently under Correction 1, supporting the conclusion that the main congruent trend in either outcome accurately reflects the true best K as K=5 according to the logic of the Evanno et al. (2005) K-selection method.*

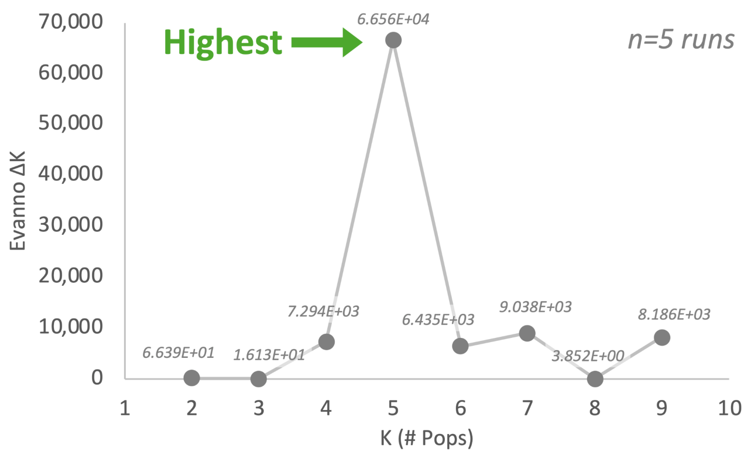

**Appendix S7.4.b (figure). CLUMPAK BestK output** using the ΔK (Evanno et al. 2005) method, run on PopCluster log-likelihood statistics (Wang 2025) generated from the neutral SNP panel (n= 177,516 SNPs) **using data correction method 2**, where log-likelihood standard deviation (SD) values equal to zero were corrected for by adding one (+1) to all SD estimates that the Evanno method tests for (K=2 through K=9).

More classic qualitative method of looking for the “elbow” in the plot of log-likelihood output corroborates K=5 as the best fit:

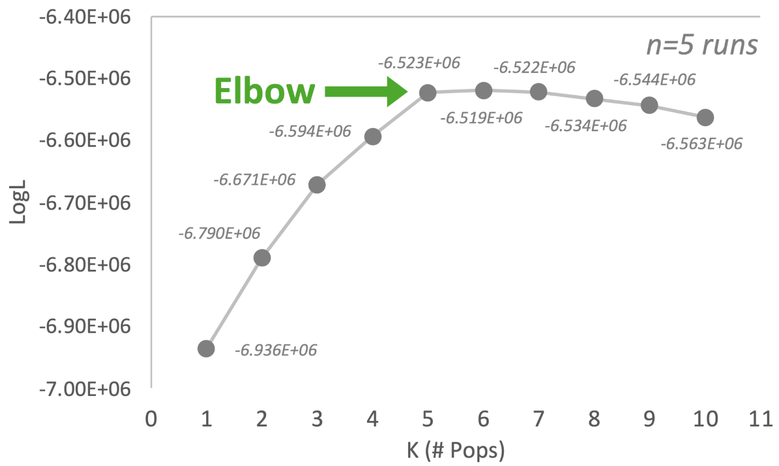

**Appendix S7.4.c (figure). Optimal K value according to the PopCluster log-likelihood statistic** (Wang 2025), calculated using the neutral SNP panel (n= 177,516 SNPs).

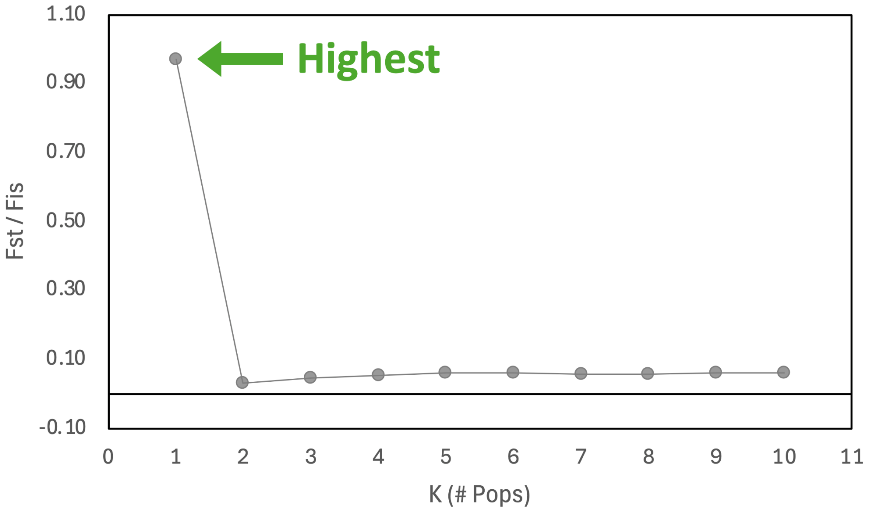

**Figure S7.15. Optimal K value according to their $F_{ST}/F_{IS}$ ratio**, an intrinsic statistic calculated **in PopCluster** (Wang 2025), calculated using the neutral SNP panel (n= 177,516 SNPs).

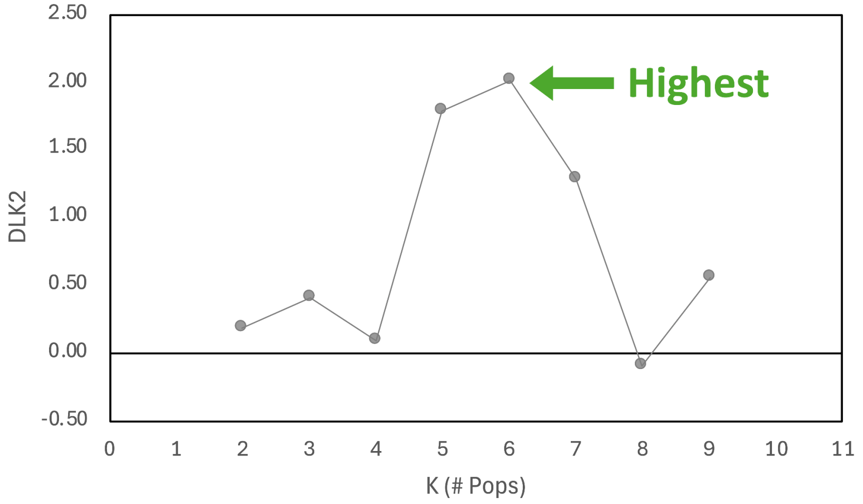

**Figure S7.16. Optimal K value according to the second-order rate of change (DLK2) value**, an intrinsic statistic calculated **in PopCluster** (Wang, 2024), calculated using the neutral SNP panel (n= 177,516 SNPs). Note: DLK2 does not evaluate K=1, the visual reflects a null value of 0.00 for this solution.

#### Appendix S8. $F_{ST}$, IBD, and migration rate results.

**Appendix S8.1. Results of pairwise $F_{ST}$ comparisons using full and neutral SNP panels among sampling locations and K=5 subpopulations.**

While $F_{ST}$ estimates suggested low differentiation across each pair of sampling locations, variation among pairwise values revealed informative structure in the relative strength of connectivity among regions. Four of the five lowest values reflected the same regional groupings seen in PCA and clustering analyses at K=5: ARU/CUR ($F_{ST}$=0.0023), DRT/FLU ($F_{ST}$=0.0040), FLL/FLU ($F_{ST}$=0.0071), and DRT/FLL ($F_{ST}$=0.0110) (Figure S8.1). BEL/MEX displayed only slightly more differentiation at $F_{ST}$=0.0161, but the final K=5 post hoc pair, DOM/JAM, showed relatively higher differentiation ($F_{ST}$=0.0331), consistent with their divergent secondary ancestry profiles in the K=5 sNMF results (Figure 1a) and their separation in the DLK2-supported PopCluster K=6 solution (Figure 1c; Figure S7.16).

One comparison outside the K=5 groupings showed lower differentiation than BEL/MEX and DOM/JAM: FLU/MEX ($F_{ST}$=0.0159) (Figure S8.1). This likely reflects the unusual placement of one FLU individual with BEL/MEX in the clustering analyses (e.g., Figure 1a,c), which may have increased within-location variation in FLU and reduced differentiation between FLU and western Caribbean sites. More broadly, all FLU pairings fell within the lower half of the sampling-location $F_{ST}$ distribution (Figure S8.2), consistent with the mixed clustering behavior observed among FLU samples. At the upper end of the distribution, the highest pairwise $F_{ST}$ values were generally observed between geographically distant sites (Figure S8.2), a trend explored further below through an isolation-by-distance (IBD) analysis. Finally, Panama showed moderate differentiation from most locations, with values ranging from $F_{ST}$=0.0571 with FLU to $F_{ST}$=0.0858 with FLL (Figure S8.1), consistent with its repeated, early (low K) emergence as a distinct cluster across ancestry-based models (Figure S7.6, Figure S7.10, Figure S7.14).

Pairwise $F_{ST}$ values among the post hoc K=5 population groupings were similarly low to moderate (Figure S8.1), and clarified trends suggested by the sampling location-based comparisons. The lowest differentiation occurred between DRT-FLL-FLU and JAM-DOM ($F_{ST}$=0.0430), followed by DRT-FLL-FLU and BEL-MEX ($F_{ST}$=0.0519), and ARU-CUR and JAM-DOM ($F_{ST}$=0.0563) (Figure S[FSTTABLES]), consistent with substantial shared ancestry among regional groups. Higher differentiation involved ARU-CUR, PAN, and BEL-MEX, including ARU-CUR vs. BEL-MEX ($F_{ST}$=0.0845), ARU-CUR vs. PAN ($F_{ST}$=0.0763), DRT-FLL-FLU vs PAN ($F_{ST}$=0.0746), and ARU-CUR vs DRT-FLL-FLU ($F_{ST}$=0.0705) (Figure S8.1). Together, these pairwise comparisons indicate detectable regional differentiation, with the strongest contrasts occurring among more geographically distinct K=5 groupings rather than as deep subdivision across the Caribbean.

**Appendix S8.2. Isolation-by-distance and effective migration rate analysis results.**

To further evaluate whether spatial separation contributed to the moderate regional structure observed across population clustering and pairwise $F_{ST}$ analyses, we tested for isolation by distance (IBD) and compared patterns of effective migration across the Caribbean. Pairwise genetic distance increased with geographic distance between sampling locations, supporting a significant but incomplete pattern of IBD (Mantel statistic=0.395, simulated p=0.022; Figure 1e). This pattern was consistent with the broader pattern observed in both the population structure analyses and pairwise $F_{ST}$ comparisons, in which genetic differentiation was generally lowest among geographically proximate or repeatedly clustered locations and higher among more distant regional comparisons (Figure 1e).

The lowest pairwise genetic distances occurred among nearby locations that also grouped together in population structure analyses, including ARU/CUR, comparisons among DRT, FLL, and FLU, and BEL/MEX (Figure 1e). The DOM/JAM comparison, which formed the remaining K=5 cluster, showed somewhat higher genetic distance but still fell within the lower-to-intermediate range of pairwise values. In contrast, many of the highest genetic distance values occurred in long-distance comparisons spanning western, southern, and eastern Caribbean locations, including comparisons involving Belize or Mexico with Curaçao, Aruba, Dominican Republic, or Panama (Figure 1e). These contrasts contributed to the overall positive IBD relationship and are consistent with geographic distance shaping part of the moderate regional structure observed across PCA, clustering, and $F_{ST}$ analyses.

However, the IBD relationship was not strictly linear, indicating that geographic distance alone did not fully account for the observed structure. Several geographically distant comparisons showed only moderate genetic distance, while some intermediate-distance comparisons showed relatively elevated differentiation. FLU comparisons consistently fell below the fitted trendline (Figure 1e), indicating that upper Florida Keys samples were, on average, more genetically similar to other locations than expected based on geographic distance alone. This pattern is consistent with the broader population structure results, including the assignment of one FLU individual to the BEL-MEX cluster across inference approaches. More generally, pairwise comparisons at the shortest geographic distances consistently fell below the fitted trendline, while comparisons exceeding the trendline occurred only at distances greater than 100 km (Figure 1e).

Effective migration-rate analyses further supported this interpretation by identifying regional differences in connectivity that were not captured by geographic distance alone. Effective migration was higher than expected under an IBD model between neighboring coastal sites (DRT-FLL-FLU, BEL-MEX, DOM-JAM, ARU-CUR), but lower than expected between sites across the Caribbean Sea (i.e. DOM-JAM vs ARU-CUR, PAN, BEL-MEX) (Figure 1f). This pattern was consistent across all deme sizes tested (Figure S8.3). However, it is important to note that migration rates only varied by a maximum of 10-fold faster or slower than the average. Nevertheless, relatively high migration between neighboring coastal sites are in support of sNMF subpopulation structure (K=5) and relatively low genetic differentiation ($F_{ST} < 0.034$) between these clustering pairs of sampling locations. However, more migrants than expected under the isolation by distance model suggests that these sampling locations are more connected than would be expected based on distance alone. For example, JAM and DOM are less differentiated than JAM compared to BEL or ARU, despite being similar geographic distance to one another, suggesting migration across the Caribbean Sea is lower than within Northern, Eastern, Western, and Southern regions (Figure 1). In addition to reduced migration across the Caribbean Sea, the Panama subpopulation (PAN) also has lower than average connectivity to neighboring coastal sites (BEL, ARU). No barriers to gene flow were detected between BEL-MEX and DRT-FLL-FLU subpopulations, due to a northern corridor in the Gulf of Mexico (Figure 1f).

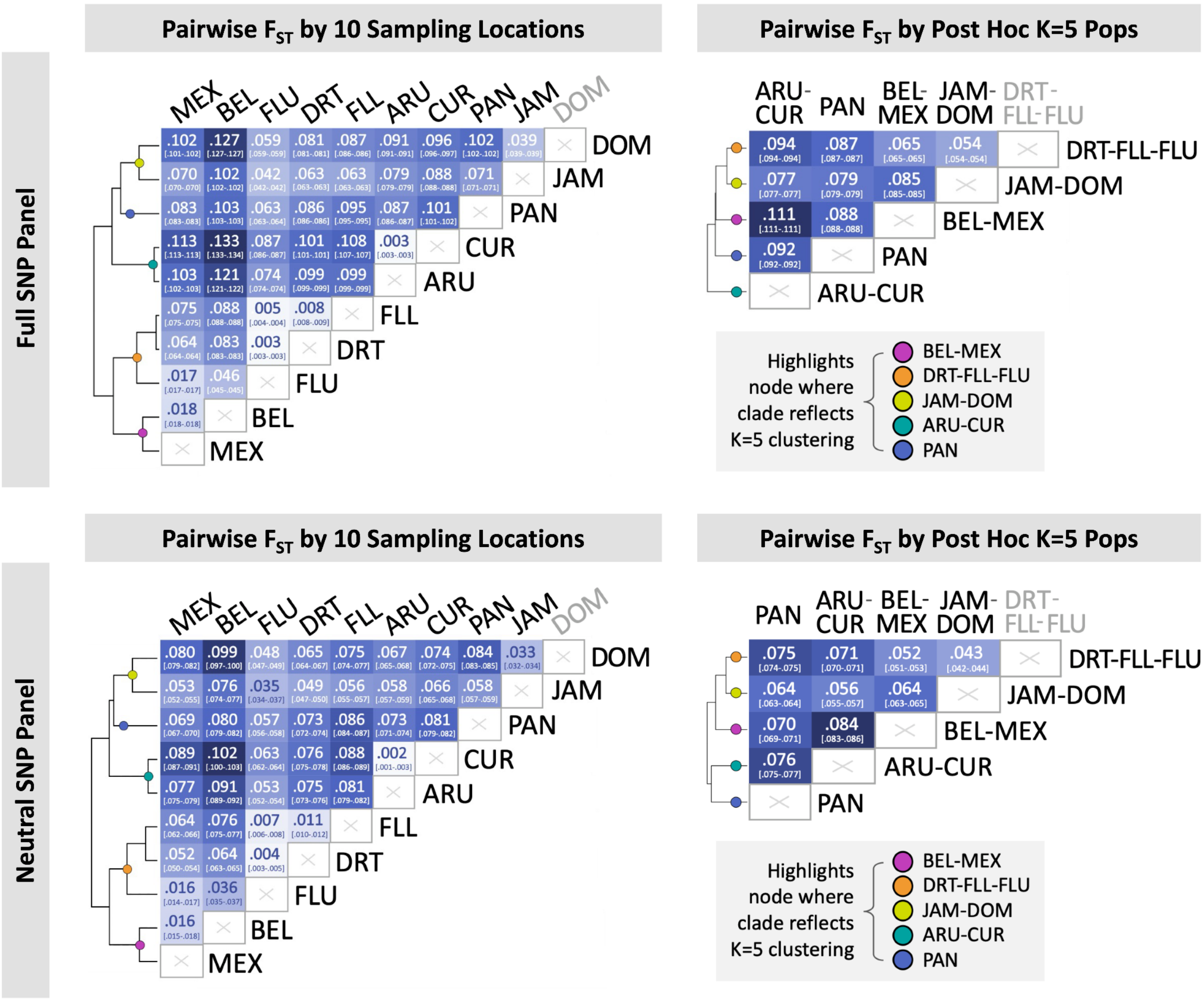

**Figure S8.1. Pairwise $F_{ST}$ calculations using full and neutral SNP panels among sampling locations and K=5 subpopulations**, organized by (1) columns: sampling location (left; n=37 subsampling to correct for unbalanced sample size, see Methods, Appendix S2.1) and by K=5 post hoc subpopulations (right; full n=46) and by (2) rows: full SNP panel (top; n= 2,527,725 SNPs) and neutral SNP panel (bottom; 177,516 SNPs). Colored circles highlight tree nodes of clades that correspond to K=5 subpopulations, illustrating the clustering of said groups even when sampling locations are treated independently and adjusted to balance sample size.

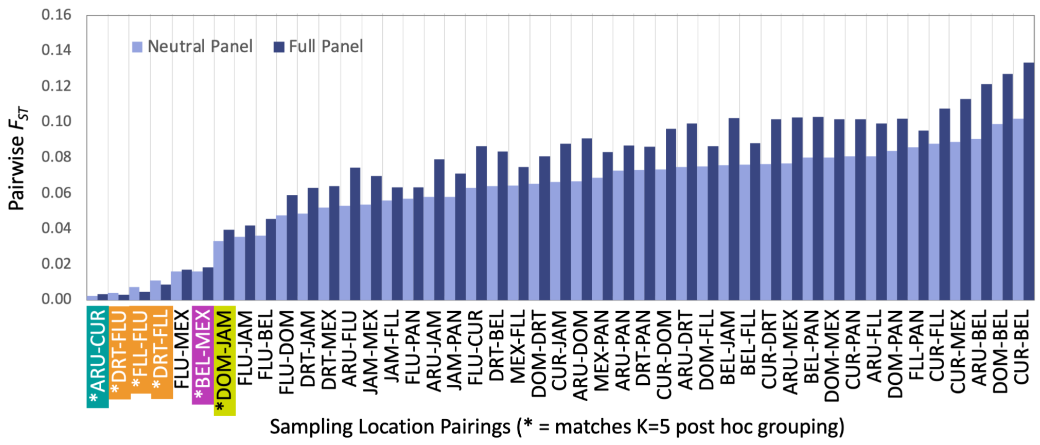

**Figure S8.2. Alternative visualization of pairwise $F_{ST}$ calculations** by sampling location across full (2,527,725 SNPs) and neutral SNP (177,516 SNPs) panels, with n=37 subsampling to correct for unbalanced sample size (details in Methods, Appendix S2.1).

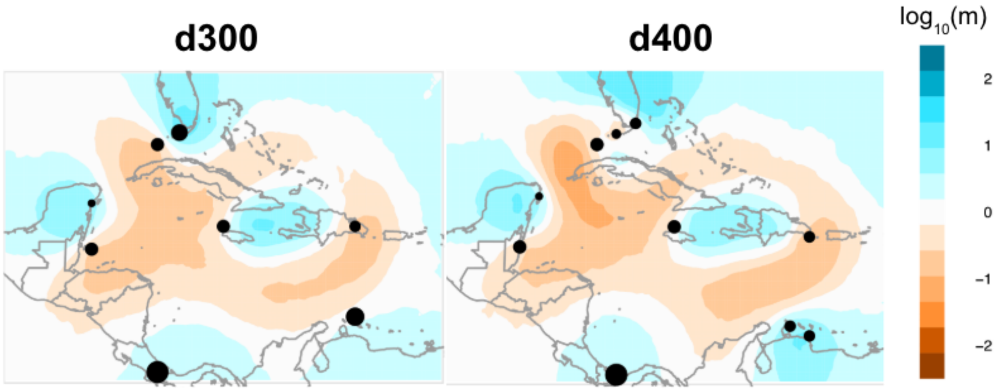

**Figure S8.3. Estimated Effective Migration Surfaces (EEMS) modeled across different deme sizes** (left - 300, right - 400). Points represent sampling locations. Colors represent the log10 of migration rates relative to average expectations under isolation by distance.

**Table S8.1. Lat long coordinates used for Haversine distance calculation in isolation by distance (IBD) analysis.**

| Longitude | Latitude | Sampling Location ID |
|---|---|---|
| -80.666967 | 24.813617 | UpperFLKeys |
| -81.413317 | 24.559483 | LowerFLKeys |
| -68.899321 | 12.085065 | Curacao |
| -68.82641 | 18.33915 | DominicanRepublic |
| -87.026667 | 20.368056 | Mexico |
| -77.764444 | 18.522778 | Jamaica |
| -69.968338 | 12.52111 | Aruba |
| -82.873187 | 24.628477 | Dry Tortugas |
| -87.75973 | 16.88806 | Belize |
| -82.140334 | 9.24254175 | Panama |

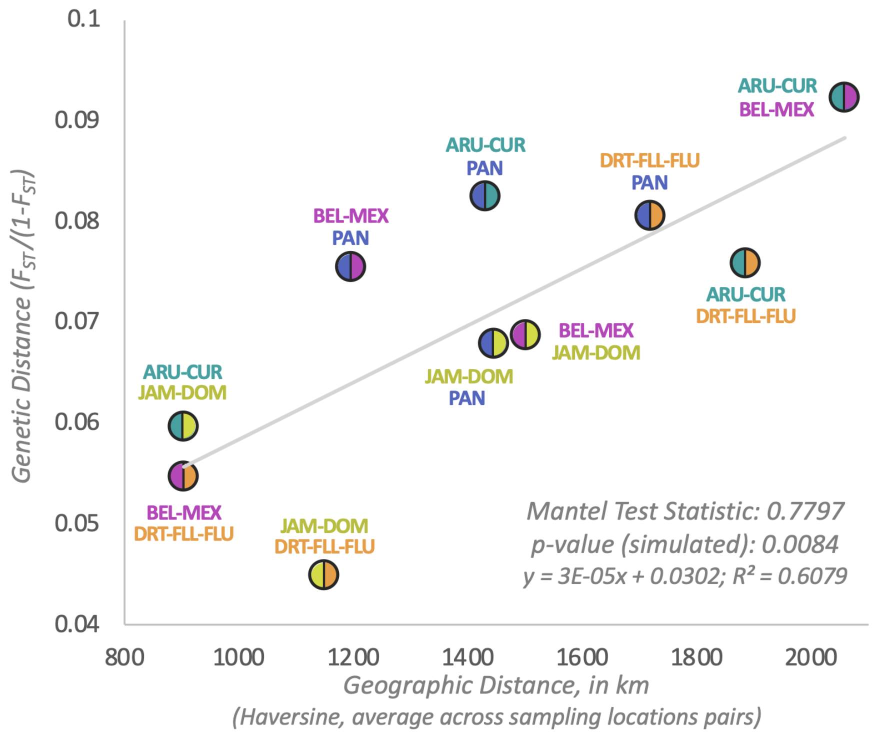

**Figure S8.3. IBD analysis exploring relationship between pairwise geographic distance** (km) **and genetic differentiation** (weighted $F_{ST}$ on neutral panel, 177,516 SNPs) **at the K=5 subpopulation scale**. Samples grouped according to K=5 subpopulation assignments. Geographic distances represent averages between each pair of samples included within broader subpopulation level comparison, calculated as described in sampling-location level IBD analysis.

### Appendix S9. Symbiont composition results.

**Table S9.1. Probe-based qPCR results for screening symbiont genera within each sample.** Values indicate the mean cycle number across technical replicates where fluorescence rose above background levels (Cq).

| Sample | Symbiodinium | Cladocopium | Durusdinium |
|---|---|---|---|
| CRF_Acer-102 | 22.63 | No Cq | No Cq |
| CRF_Acer-099 | 22.98 | No Cq | No Cq |
| CRF_Acer-059 | 23.245 | No Cq | No Cq |
| CRF_Acer-120 | 21.65 | No Cq | No Cq |
| Mote_AC80 | 22.305 | No Cq | No Cq |
| Mote_AC75 | 22.415 | No Cq | No Cq |
| Mote_AC76 | 22.64 | No Cq | No Cq |
| CU21E_1049 | 24.955 | No Cq | No Cq |
| CU21E_1050 | 22.655 | No Cq | No Cq |
| CU21E_1051 | 24.18 | No Cq | 39.9 |
| CU21E_1052 | 24.845 | No Cq | No Cq |
| CU21E_1053 | 23.345 | No Cq | No Cq |
| DR_C012 | 23.165 | No Cq | 39.545 |
| DR_C051 | 22.87 | No Cq | 39.645 |
| DR_C080 | 23.605 | No Cq | No Cq |
| DR_C171 | 25.95 | No Cq | No Cq |
| MEX_01 | 23.32 | No Cq | No Cq |
| MEX_03 | 23.225 | No Cq | No Cq |
| JAM_01 | 23.875 | No Cq | No Cq |
| JAM_02 | 25.615 | No Cq | No Cq |
| JAM_05 | 23.705 | No Cq | No Cq |
| JAM_07 | 25.475 | No Cq | No Cq |
| JAM_11 | 24.87 | No Cq | No Cq |
| IBBZ_118 | 22.91 | No Cq | No Cq |
| IBBZ_123 | 24.085 | No Cq | No Cq |
| IBBZ_13837 | 25.86 | No Cq | No Cq |
| IBBZ_13797 | 23.48 | No Cq | No Cq |
| IBBZ_13829 | 22.53 | No Cq | No Cq |
| AR_113 | 23.59 | 38.585 | No Cq |
| AR_115 | 24.355 | No Cq | No Cq |
| AR_117 | 24.605 | No Cq | No Cq |
| AR_119 | 24.32 | No Cq | No Cq |
| DRTO_151 | 22.81 | No Cq | No Cq |
| DRTO_178 | 22.76 | No Cq | No Cq |
| DRTO_143 | 23.57 | No Cq | No Cq |
| DRTO_114 | 22.815 | No Cq | No Cq |
| DRTO_52 | 23.595 | No Cq | No Cq |

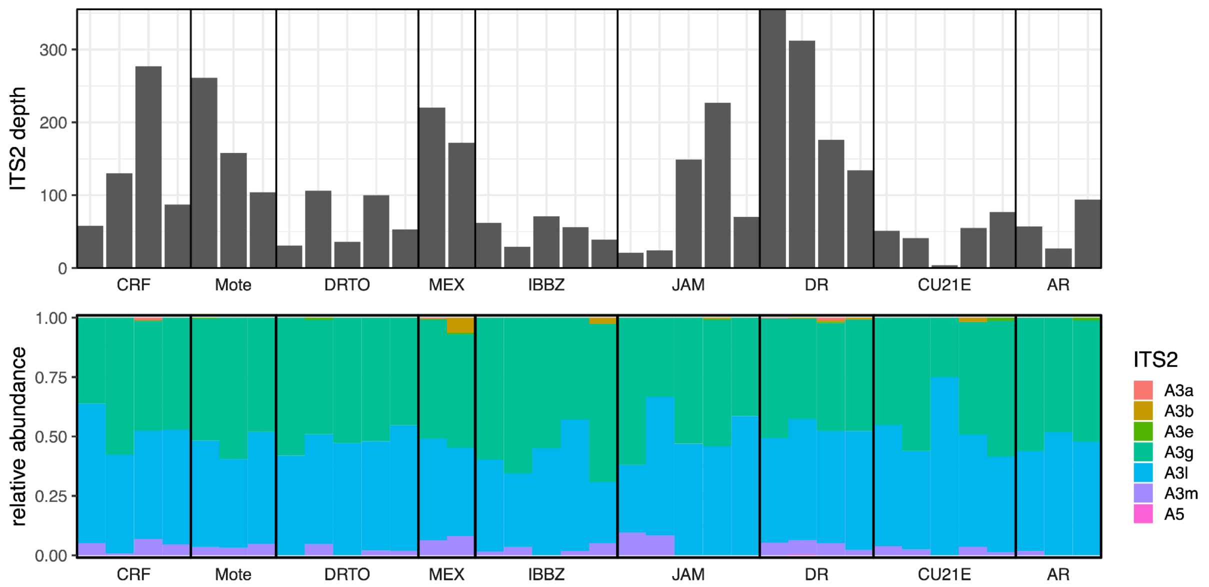

**Figure S9.1. Symbiont ITS2 read depth** (top) **and composition** (bottom). The top plot shows the read depth of the ITS2 region recovered from whole genome sequencing of the holobiont. The bottom plot shows the relative abundance of ITS2 variants. Each bar represents a single individual, grouped by sampling location. The bars in the bottom figure are colored by ITS2 variant.

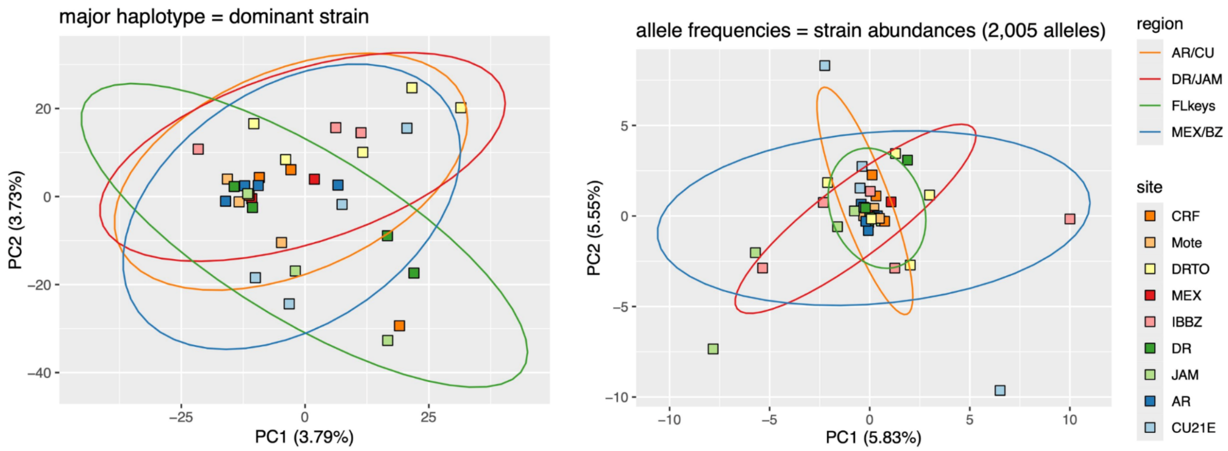

**Figure S9.2. Principal component analysis of dominant symbiont haplotypes** (left) **and overall allele frequencies** (right). Individual samples (points) are colored by location and grouped by region within ellipses. Dominant haplotypes (left) putatively represent dominant symbiont strains whereas allele frequencies (right) consider the abundance of multiple variants (symbiont population) which could be present within an individual coral colony.

### Appendix S10. Genomic diversity and demographic history results.

**Appendix S10.1. "*Patterns of genomic diversity & demographic history*" detailed results.**

JAM-DOM had the lowest global Tajima's D (-0.339), and DRT-FLL-FLU the highest, though still negative, value (-0.147); other subpopulations were intermediate (BEL-MEX: -0.256, PAN: -0.243, ARU-CUR: -0.218). Nucleotide diversity was highest in BEL-MEX (0.00240) and lowest in ARU-CUR (0.00219), but otherwise similar across subpopulations (DRT-FLL-FLU: 0.00224, JAM-DOM: 0.00224, PAN: 0.00226).

The mean fraction of the genome that is covered by ROH (FROH) across subpopulations was similar across subpopulations for runs > 100kb (ARU-CUR 0.0879; BEL-MEX 0.0722; FL 0.0639; JAM-DOM 0.0699, PAN 0.0921). When including all ROH > 100kb, subpopulation mean IDrisk scores ranged from 0.124 (FL) to 0.173 (PAN), with all subpopulations falling within a 'Moderate Risk' as assessed by Kyriazis et al. 2025 (ARU-CUR 0.164; BEL-MEX 0.141; JAM-DOM 0.130).

**Appendix S10.2. Results for genetic diversity versus genetic uniqueness analysis.**

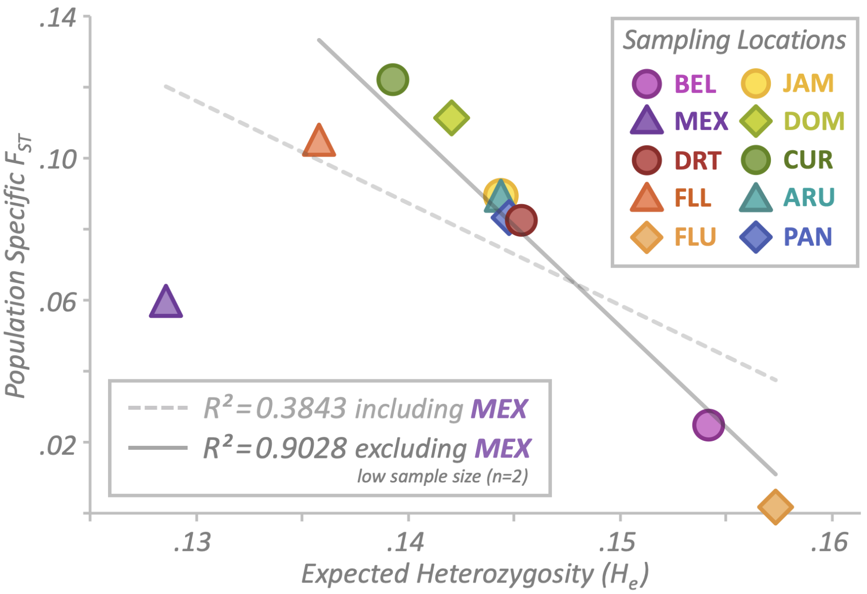

**Figure S10.1. Negative relationship between genetic "uniqueness"** (mean population-specific $F_{ST}$, full SNP panel) **and genetic diversity** (mean expected heterozygosity, $H_e$) **across the 10 sampling locations of this study.** $R^2$ results of linear regression shown both with and without MEX due to low sample size (n=2). Additional regression stats for the dataset including and excluding MEX, respectively: insignificant negative relationship (r=−0.620; p-value=0.056); highly significant negative relationship (r=−0.951; p-value=$8.2\times10^{-5}$).

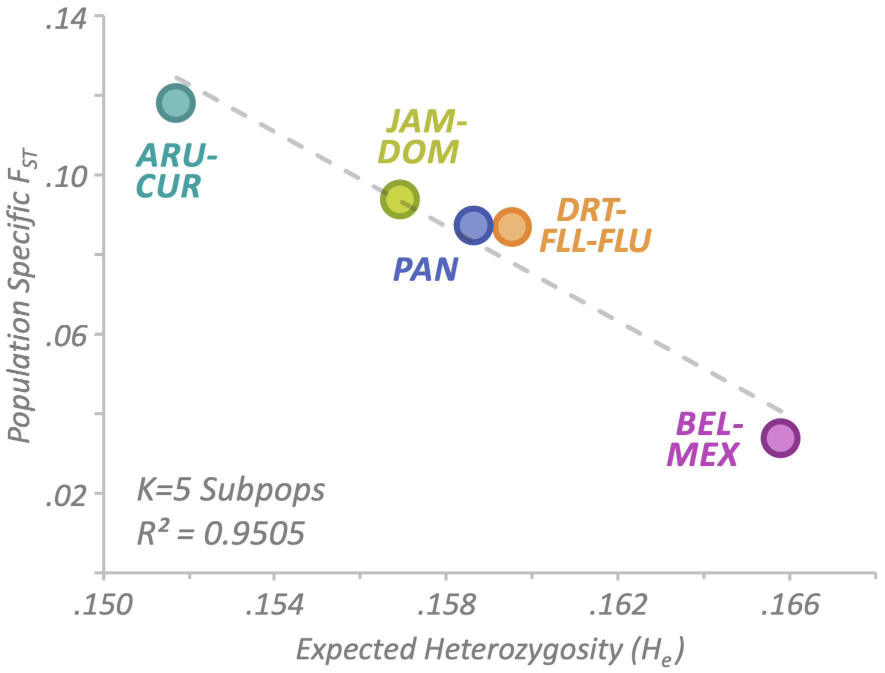

**Figure S10.2. Negative relationship between genetic "uniqueness"** (mean population-specific $F_{ST}$, full SNP panel) **and genetic diversity** (mean expected heterozygosity, $H_e$) **across K=5 subpopulations**. Additional regression stats: highly significant negative relationship (r=−0.975; p-value=0.0047).

**Table S10.1. Raw values calculated for mean population-specific $F_{ST}$** (full SNP panel) **and mean $H_e$ across the 10 sampling locations of this study.**

| Sampling Location | Expected Heterozygosity ($H_e$) | Population-Specific $F_{ST}$ |
|---|---|---|
| Belize | 0.1542 | 0.0246 |
| Mexico | 0.1285 | 0.0599 |
| Dry Tortugas | 0.1454 | 0.0826 |
| Lower FL Keys | 0.1358 | 0.1052 |
| Upper FL Keys | 0.1573 | 0.0016 |
| Jamaica | 0.1444 | 0.0895 |
| Dominican Republic | 0.1421 | 0.1114 |
| Curacao | 0.1393 | 0.1222 |
| Aruba | 0.1444 | 0.0897 |
| Panama | 0.1448 | 0.0833 |

**Table S10.2. Raw values calculated for mean population-specific $F_{ST}$** (full SNP panel) **and mean $H_e$ across K=5 subpopulations.**

| Subpopulation | Expected Heterozygosity ($H_e$) | Population-Specific $F_{ST}$ |
|---|---|---|
| BEL-MEX | 0.1658 | 0.0334 |
| DRT-FLL-FLU | 0.1595 | 0.0867 |
| JAM-DOM | 0.1569 | 0.0938 |
| ARU-CUR | 0.1517 | 0.1179 |
| PAN | 0.1587 | 0.0871 |

**Table S10.3. Results from analysis of Runs of Homozygosity (ROH) across all samples.** Both results when retaining only ROH > 100kb and > 200 kb are included. FROH = ROH fraction of genomic coverage; $ID_{risk}$ score = calculated using fraction of the genome contained in functionally long ROH and heterozygosity outside ROH regions, as in (Kyriazis et al. 2025).

| Sample ID | Heterozygosity/kb (in non-ROH regions) | Min. ROH Length = 100 kb | | Min. ROH Length = 200 kb | |
|---|---|---|---|---|---|
| | | FROH | $ID_{risk}$ Score | FROH | $ID_{risk}$ Score |
| AR_113 | 1.862 | 0.08618 | 0.1605 | 0.03249 | 0.06050 |
| AR_115 | 2.067 | 0.06312 | 0.1305 | 0.01416 | 0.02926 |
| AR_117 | 1.804 | 0.09428 | 0.1701 | 0.03992 | 0.07202 |
| AR_119 | 1.884 | 0.08855 | 0.1668 | 0.02123 | 0.04000 |
| CRF_Acer-059 | 1.958 | 0.06416 | 0.1256 | 0.01783 | 0.03491 |
| CRF_Acer-099 | 2.032 | 0.04963 | 0.1008 | 0.01167 | 0.02372 |
| CRF_Acer-102 | 2.157 | 0.05263 | 0.1135 | 0.02023 | 0.04365 |
| CRF_Acer-120 | 1.855 | 0.06761 | 0.1254 | 0.02410 | 0.04471 |
| CU21E_1050 | 1.856 | 0.09462 | 0.1757 | 0.03468 | 0.06438 |
| CU21E_1051 | 1.795 | 0.09489 | 0.1703 | 0.03470 | 0.06228 |
| CU21E_1052 | 1.823 | 0.09793 | 0.1786 | 0.03599 | 0.06562 |
| CU21E_1053 | 1.890 | 0.08371 | 0.1582 | 0.02551 | 0.04822 |
| DR_C012 | 1.752 | 0.08843 | 0.1549 | 0.02743 | 0.04805 |
| DR_C051 | 2.474 | 0.01756 | 0.0434 | 0.00307 | 0.00759 |
| DR_C080 | 1.736 | 0.09859 | 0.1712 | 0.03430 | 0.05955 |
| DR_C171 | 1.886 | 0.08207 | 0.1548 | 0.02516 | 0.04745 |
| DRTO_114 | 1.909 | 0.08074 | 0.1541 | 0.02449 | 0.04675 |
| DRTO_143 | 1.899 | 0.07350 | 0.1396 | 0.02354 | 0.04470 |
| DRTO_151 | 2.016 | 0.06756 | 0.1362 | 0.02509 | 0.05060 |
| DRTO_178 | 1.925 | 0.07133 | 0.1373 | 0.02107 | 0.04055 |
| DRTO_52 | 1.929 | 0.05840 | 0.1126 | 0.01428 | 0.02755 |
| IBBZ_118 | 2.002 | 0.08089 | 0.1619 | 0.02252 | 0.04509 |
| IBBZ_123 | 2.016 | 0.07321 | 0.1476 | 0.02716 | 0.05475 |
| IBBZ_13797 | 1.969 | 0.06676 | 0.1314 | 0.02153 | 0.04238 |
| IBBZ_13829 | 1.994 | 0.06253 | 0.1247 | 0.01726 | 0.03441 |
| IBBZ_13837 | 1.848 | 0.08561 | 0.1582 | 0.02570 | 0.04748 |
| JAM_01 | 1.833 | 0.06868 | 0.1259 | 0.01891 | 0.03465 |
| JAM_02 | 1.958 | 0.06165 | 0.1207 | 0.01412 | 0.02764 |
| JAM_05 | 1.833 | 0.07851 | 0.1439 | 0.02824 | 0.05177 |
| JAM_07 | 1.880 | 0.06256 | 0.1176 | 0.01722 | 0.03238 |
| JAM_11 | 1.936 | 0.07115 | 0.1377 | 0.02053 | 0.03975 |
| MEX_01 | 1.943 | 0.06178 | 0.1200 | 0.01708 | 0.03319 |

| | | | | | | | |
|---|---|---|---|---|---|---|---|
| MEX_03 | 1.931 | | 0.07477 | 0.1444 | | 0.02695 | 0.05205 |
| Mote_AC75 | 1.827 | | 0.06206 | 0.1134 | | 0.02058 | 0.03760 |
| Mote_AC76 | 1.972 | | 0.05480 | 0.1081 | | 0.01782 | 0.03513 |
| Mote_AC80 | 1.937 | | 0.06482 | 0.1256 | | 0.02038 | 0.03949 |
| SRR24007593 | 1.905 | | 0.09326 | 0.1777 | | 0.02678 | 0.05102 |
| SRR24007597 | 1.872 | | 0.09032 | 0.1690 | | 0.03676 | 0.06879 |
| SRR24007601 | 2.007 | | 0.08603 | 0.1727 | | 0.02644 | 0.05308 |
| SRR24007602 | 1.844 | | 0.09462 | 0.1744 | | 0.03543 | 0.06531 |
| SRR24007603 | 1.860 | | 0.10590 | 0.1970 | | 0.04128 | 0.07679 |
| SRR24007608 | 2.033 | | 0.05748 | 0.1169 | | 0.01340 | 0.02724 |
| SRR24007609 | 1.922 | | 0.08764 | 0.1684 | | 0.03012 | 0.05788 |
| SRR24007652 | 1.895 | | 0.09243 | 0.1751 | | 0.04873 | 0.09233 |
| SRR24007659 | 1.862 | | 0.09043 | 0.1684 | | 0.02883 | 0.05369 |
| SRR24007661 | 1.722 | | 0.12332 | 0.2123 | | 0.04582 | 0.07888 |

### Appendix S11. Results of adaptive outlier scans.

**Appendix S11.1. Detailed results associated with adaptive variation scans at the metapopulation (K=1) scale.**

Six 10kb regions were identified to be putatively under selection in the metapopulation (Figure S11.1). Three outlier regions had low nucleotide diversity (<6e-5) and Tajima's D (<-2), indicative of purifying selection. Five genes were identified within these three regions, two of which had functional annotations within the same window (OZ035975.1: 320487-330486), including WW domain-binding protein 11 (*WBP11*) and ATP-binding cassette subfamily A member 3 (*ABCA3*). The other three outlier regions exhibited high nucleotide diversity (>0.04) and Tajima's D (>2) indicating balancing selection. Two genes out of 10 total were annotated across these three windows: 2'-5'-oligoadenylate synthetase 3 (*OAS3*) and tetratricopeptide repeat domain 37 (TTC37). *OAS3* and *TTC37* were located within the same outlier window (OZ035966.1: 27680399-27690398) and represented two out of three possible coding sequences. Both *OAS3* and *TTC37* are involved in RNA degradatory pathways (Sadler & Williams 2008; Zhang et al. 2019). However, *OAS3* is an interferon-stimulated gene which is activated by viral dsRNA, playing a critical role in the innate immune response (Sadler & Williams 2008). Due to its possible role in coral disease, we further investigated whether the *OAS3* gene was under divergent selection across populations. Sample-specific genotypes spanning the *OAS3* gene did not cluster based on sampling region or population (Figure S11.2), suggesting that balancing selection of this gene is not being driven by local adaptation.

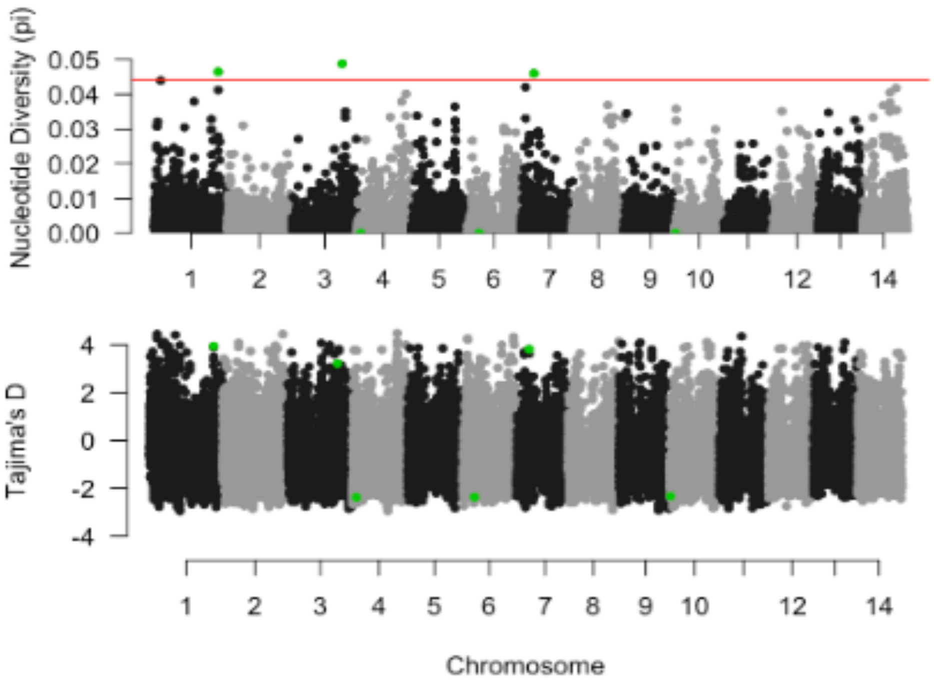

**Figure S11.1. Caribbean-wide nucleotide diversity** (top) **and Tajima's D** (bottom) **across the genome using 10kb windows.** Candidate outlier windows colored in green were identified as those within the top or bottom 0.01% of the genome-wide pi distribution (red line) that also exhibited signatures of balancing or purifying selection (Tajima's D >2 or <-2 respectively).

**Table S11.1. Annotations for putative regions under balancing/purifying selection at the metapopulation level.**

| chrom | start | end | selection | annot_chrom | annot_start | annot_end | annot |
|---|---|---|---|---|---|---|---|
| OZ035966.1 | 27680399 | 27690398 | balancing | OZ035966.1 | 27682169 | 27682478 | ID=FUN_003958; |
| OZ035966.1 | 27680399 | 27690398 | balancing | OZ035966.1 | 27683191 | 27691138 | ID=FUN_003959;Name=OAS3; |
| OZ035966.1 | 27680399 | 27690398 | balancing | OZ035966.1 | 27646304 | 27680786 | ID=FUN_003957;Name=TTC37; |
| OZ035968.1 | 21450004 | 21460003 | balancing | OZ035968.1 | 21403951 | 21485114 | ID=FUN_009759;Name=DCHS1_1; |
| OZ035972.1 | 5485645 | 5495644 | balancing | OZ035972.1 | 5485989 | 5487390 | ID=FUN_019118; |
| OZ035972.1 | 5485645 | 5495644 | balancing | OZ035972.1 | 5489820 | 5490376 | ID=FUN_019120; |
| OZ035972.1 | 5485645 | 5495644 | balancing | OZ035972.1 | 5490874 | 5491803 | ID=FUN_019121; |
| OZ035972.1 | 5485645 | 5495644 | balancing | OZ035972.1 | 5492798 | 5493168 | ID=FUN_019122; |
| OZ035972.1 | 5485645 | 5495644 | balancing | OZ035972.1 | 5487943 | 5488780 | ID=FUN_019119; |
| OZ035975.1 | 320487 | 330486 | purifying | OZ035975.1 | 316526 | 326745 | ID=FUN_027100;Name=WBP11; |
| OZ035975.1 | 320487 | 330486 | purifying | OZ035975.1 | 326016 | 327186 | ID=FUN_027101; |
| OZ035975.1 | 320487 | 330486 | purifying | OZ035975.1 | 328498 | 349259 | ID=FUN_027102;Name=ABCA3; |
| OZ035969.1 | 1733076 | 1743075 | purifying | NA | NA | NA | NA |
| OZ035971.1 | 5190089 | 5200088 | purifying | OZ035971.1 | 5181257 | 5272144 | ID=FUN_016522; |

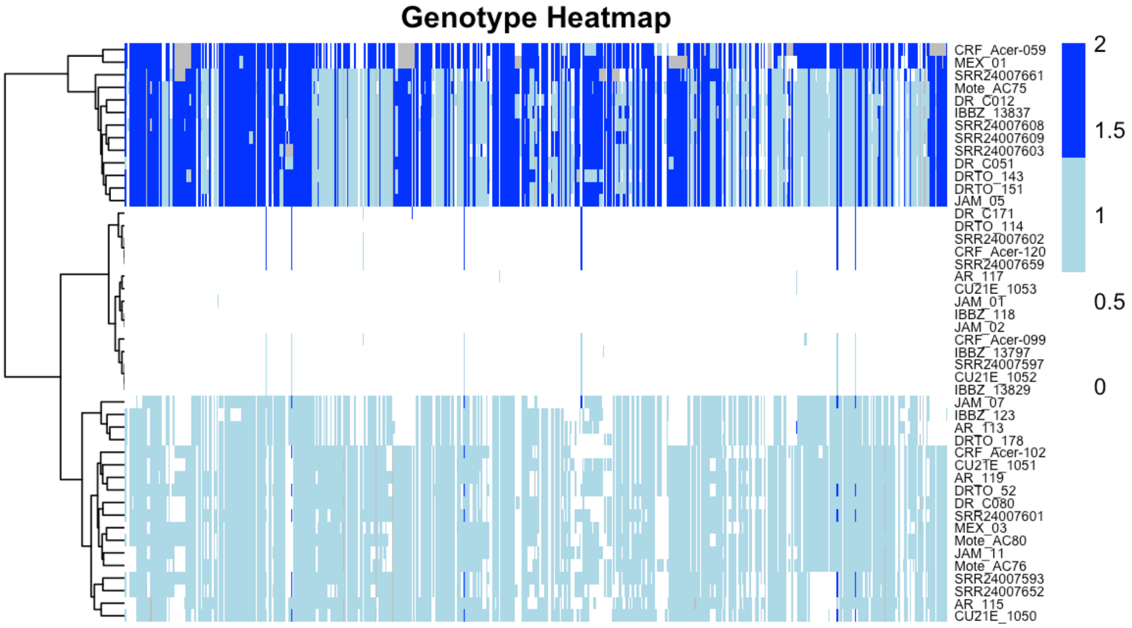

**Figure S11.2. Clustering analysis of sample-specific genotypes across the OAS3 gene.** Each column represents a position within the OAS3 gene. Genotypes are colored based on the number of alternative alleles as follows 0: white – homozygous reference, 1: light blue – heterozygous, 2: dark blue – homozygous alternate, with gray boxes denoting missing genotypes. Each row represents a single sample and samples are clustered by genotype.

**Appendix S11.2. Detailed meadow plot results associated with adaptive variation scans at the subpopulation (K=5) scale.**

Given the limited evidence from locus-based scans, we used meadow plots (Duffin et al. in press) as a complementary window-based approach to evaluate whether the K=5 structure signal was associated with broader, localized sweep-like patterns. Across the 10 pairwise comparisons across regions (BEL-MEX, DRT-FLL-FLU, JAM-DOM, ARU-CUR and PAN), candidate outlier windows with elevated $F_{ST}$ and extreme shifts in nucleotide diversity were detected but were generally sparse, rather than forming broad, repeated peaks shared across population pairs (Figure S11.3). The total number of 10 kb outlier windows varied substantially among comparisons, ranging from seven in JAM-DOM vs. PAN to 29 in DRT-FLL-FLU vs. JAM-DOM (Figure S11.4). However, these counts did not recapitulate trends in overall genetic differentiation or geographic distance. For example, ARU-CUR vs. BEL-MEX had the highest pairwise $F_{ST}$ among K=5 groupings but only 19 outlier windows, whereas DRT-FLL-FLU vs. JAM-DOM had the lowest pairwise $F_{ST}$ but the highest number of outlier windows (Figure S11.4, Figure S11.5). Moreover, the contrasts with the fewest and among the most outlier windows, JAM-DOM vs. PAN and ARU-CUR vs. PAN had nearly identical average geographic distances (~1447 and ~1431 km, respectively; range across all comparisons: ~904 to ~2059 km; Figure S11.6). This suggests that the number of regions showing signatures of local adaptation were not simply a reflection of the same patterns observed under genome-wide neutral differentiation, nor physical separation among groups.

Most candidate regions were short, although a small number of comparisons contained extended runs of consecutive outlier windows (Figure S11.4). These four widest peaks, each around 100 kb, all occurred in different meadow plot comparisons involving PAN or DRT-FLL-FLU/BEL-MEX (Table S11.2). The extended runs in the DRT-FLL-FLU vs. BEL-MEX and PAN vs. BEL-MEX comparisons mapped to chromosomes 2 and 5, respectively (scaffolds OZ035968.1 and OZ035971.1). Interestingly, the remaining two (ARU-CUR vs. PAN and DRT-FLL-FLU vs. PAN) co-localized to the same region of chromosome 9 (OZ035974.1:18780001-18990000) (Table S11.2). Further, all outlier windows across both comparisons were driven by markedly reduced nucleotide diversity in Panama, as indicated by the direction of the $\log_2$ pi-ratio values (Table S11.2). The shared chromosome 9 candidate region overlapped 21 predicted gene features, most of which were annotated only by FUN gene-model identifiers (Table S11.3). The only overlapping feature with an informative gene symbol was *FUN_026732/LYS1_7* (OZ035974.1:18766278-18788722) (Table S11.3). Because *LYS1* is associated with lysine biosynthesis and metabolism (Garrad and Bhattacharjee, 1992), this annotation may point to amino acid metabolism as a potentially relevant target for *A. cervicornis* (Fitzgerald and Szmant, 1997) within the shared chromosome 9 sweep-like region. However, given that most overlapping features lacked informative functional names and that the *LYS1_7* annotation is putative, this candidate should be interpreted cautiously pending additional validation.

Overall, meadow plots did not reveal dominant and repeated signatures of divergent selection among the sampling location-based groups (Figure S11.3) but instead identified a limited set of comparison-specific candidate regions (Table S11.3). This pattern is consistent with the broader population structure results, in which regional differentiation was detectable but weak relative to the prevailing signal of Caribbean-wide connectivity.

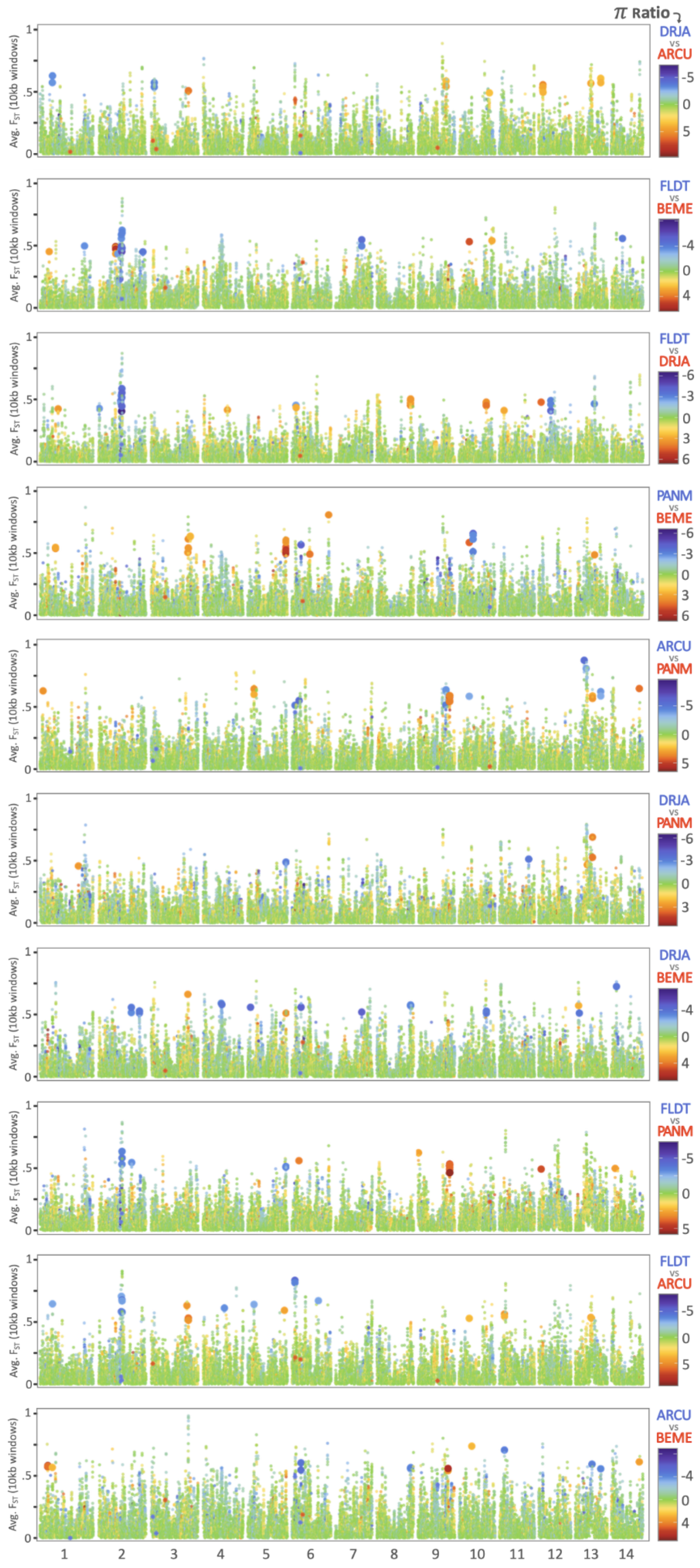
π Ratio
DRJA vs ARCU
FLDT vs BEME
FLDT vs DRJA
PANM vs BEME
ARCU vs PANM
DRJA vs PANM
DRJA vs BEME
FLDT vs PANM
FLDT vs ARCU
ARCU vs BEME
Avg. FST (10kb windows)
1 2 3 4 5 6 7 8 9 10 11 12 13 14

**Figure S11.3.** (previous page) **Meadow plots comparing pairwise K=5 subpopulations to visualize putative selective sweeps** as genomic regions of outlier $F_{ST}$ and $\log_2 \pi$ ratios across 10 kb windows. See Duffin et al. (in press) for conceptual description and Appendix S4.1 for methods details. Briefly, windows falling within both the top 1% of $F_{ST}$ and the top or bottom 1% of $\log_2 \pi$ ratios are considered outliers, with datapoints representing outlier windows scaled up in size for emphasis. Cooler and warmer colors distinguish reduced diversity in either subpopulation under comparison. Asterisk-shaped data points represent comparisons where one of the two populations under comparison had a π value of zero, resulting in a mathematical error calculation (+/- inf) for the $\log_2 \pi$ ratio. K=5 subpopulation abbreviations: BEME = Belize-Mexico, FLDT = Dry Tortugas-Florida Keys (Lower)-Florida Keys (Upper), DRJA = Dominican Republic-Jamaica, ARCU = Aruba-Curaçao, PANM = Panama.

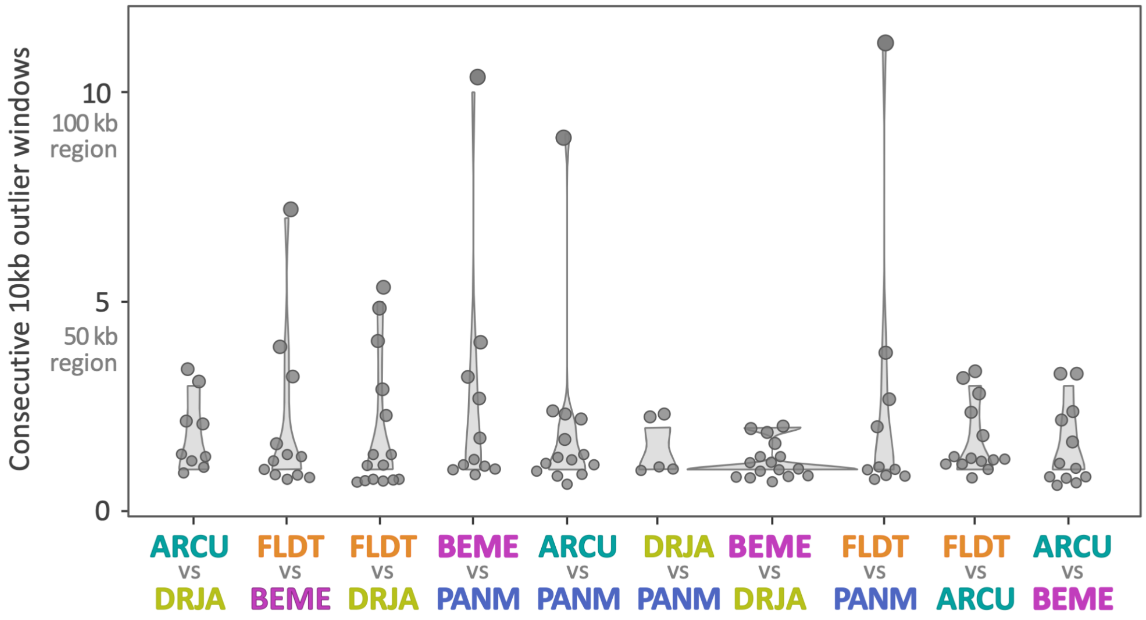

| | ARCU vs DRJA | FLDT vs BEME | FLDT vs DRJA | BEME vs PANM | ARCU vs PANM | DRJA vs PANM | BEME vs DRJA | FLDT vs PANM | FLDT vs ARCU | ARCU vs BEME |
|---|---|---|---|---|---|---|---|---|---|---|
| Geo. dist. (km) | 904 | 905 | 1151 | 1196 | 1431 | 1447 | 1502 | 1721 | 1886 | 2059 |
| $F_{ST}$ All | 0.077 | 0.065 | 0.054 | 0.088 | 0.092 | 0.079 | 0.085 | 0.087 | 0.094 | 0.111 |
| $F_{ST}$ Neutral | 0.056 | 0.052 | 0.043 | 0.070 | 0.076 | 0.064 | 0.064 | 0.075 | 0.071 | 0.084 |
| No. outlier windows | 19 | 24 | 29 | 28 | 26 | 7 | 20 | 22 | 22 | 19 |
| (Pop 1 / Pop 2) | 4 / 15 | 14 / 10 | 15 / 14 | 5 / 23 | 10 / 16 | 3 / 4 | 17 / 3 | 7 / 15 | 12 / 10 | 8 / 11 |

Pop 1, e.g. ARCU (left) & Pop 2, e.g. DRJA (right)

**Figure S11.4. Consecutive outlier windows among pairwise** subpopulation **m**eadow plot **comparisons.** Top: Violin plots summarizing "peak length distributions," or the spread of consecutive outlier windows among pairwise K=5 subpopulation-based comparisons. See main text methods and Table S11.2 for outlier window span scoring details. Bottom: Table displaying statistics of interest for each pairwise sampling-location comparison, including: geographic distances (calculated as described in Figure S8.3), genetic differentiation (pairwise $F_{ST}$) using full and neutral SNP panels (see Appendix S8.1; Figure S8.1); and total number of outlier windows summed for each meadow plot, collectively (top cell) and parsed by driving subpopulation (bottom cells). K=5 subpopulation abbreviations: BEME = Belize-Mexico, FLDT = Dry Tortugas-Florida Keys (Lower)-Florida Keys (Upper), DRJA = Dominican Republic-Jamaica, ARCU = Aruba-Curaçao, PANM = Panama.

**Table S11.2. Characterization of the four longest consecutive outlier windows from meadow plot comparisons.** See detailed text below table for explanation of window span calculations [briefly: total span = size (+ non- outlier winds.); span (adj) = size "score" (adjusted)]. K=5 subpopulation abbreviations: BEME = Belize-Mexico, FLDT = Dry Tortugas-Florida Keys (Lower)-Florida Keys (Upper), DRJA = Dominican Republic-Jamaica, ARCU = Aruba-Curaçao, PANM = Panama.

| comparison | chr | positions | outlier window pattern<br>*" ", with omitted windows ((#))* | total span | span (adj) | driven by low pi in... |
|---|---|---|---|---|---|---|
| FLDT vs. BEME | 2 | 9,900,001-9,980,000 | **2-[1]-5** = 7 "consecutive"<br>*(same as above; no omitted windows)* | 80 kb | 70 kb | BEME |
| PANM vs. BEME | 5 | 22,740,001-22,880,000 | **2-[2]-5-[1]-1-[1]-2** = 10 "consecutive"<br>*(same as above; no omitted windows)* | 140 kb | 100 kb | BEME |
| ARCU vs. PANM | 9 | 18,770,001-18,990,000 | **8-[1]-1** = 9 "consecutive"<br>*2-((4))-3-((1))-3-[2]-1-[1]-((1))-[3]-1* | 220 kb | 90 kb | PANM |
| FLDT vs. PANM | 9 | 18,770,001-18,990,000 | **8-[2]-1-[3]-2** = 11 "consecutive"<br>*2-((4))-3-((1))-3-[2]-1-[1]-((1))-[2]-2* | 220 kb | 110 kb | PANM |

*Shorthand notation of outlier window pattern example:* **8-[2]-1-[3]-2**

*Translates to: eight consecutive 10kb outlier windows, then two 10kb windows not meeting outlier threshold, followed by another one window meeting threshold, three not meeting threshold, then another two 10kb outlier windows. While the collective outlier peak window technically spans 220kb, non-outlier windows are excluded and the reported size "score" ["span (adj)"] is (8+1+2)=11 10kb windows, or 110kb*

*That said, (8+[2]+1+[3]+2)=16 10kb windows, or 160kb, which does not equal the 220kb total reported span ("total span" column) because several windows in the range of 18770kb to 18980kb were removed upstream due to <5kb SNPs present. See diagram below for additional clarification. The full notation including outlier windows, non-outlier windows "[#]", and excluded windows "((#))" is given beneath the bolded main notation in grey, italicized font in the column "outlier window pattern."*

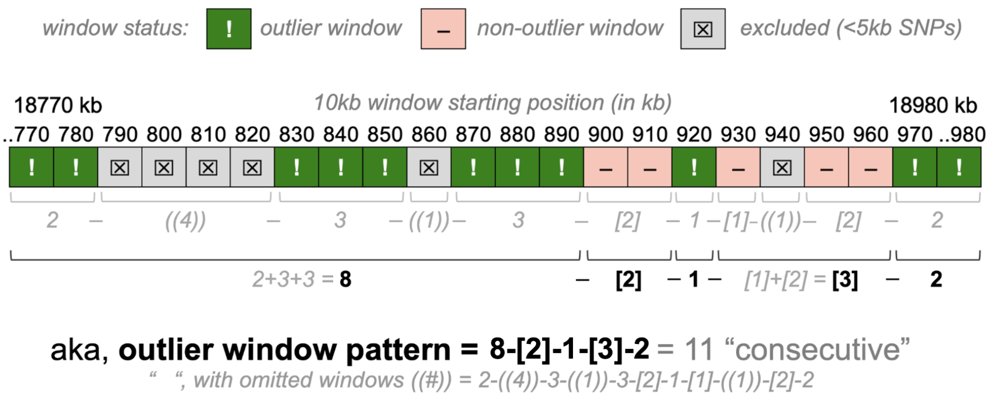

**Table S11.3. Predicted gene features overlapping the shared candidate region on chromosome 9 (OZ035974.1)**, jointly identified in ARU-CUR vs. PAN and DRT-FLL-FLU vs. PAN meadow plot analyses. Gene features were identified by intersecting the co-localized outlier interval (OZ035974.1:18,780,001-18,990,000) with the *A. cervicornis* functional annotation (Locatelli 2024). Gene model IDs are shown for all overlapping features; gene symbols are reported where available.

| scaffold | feature_type | start_pos | end_pos | gene_model_ID | gene_symbol |
|---|---|---|---|---|---|
| OZ035974.1 | gene | 18766278 | 18788722 | FUN_026732 | *LYS1_7* |
| OZ035974.1 | gene | 18794476 | 18794790 | FUN_026733 | -- |
| OZ035974.1 | gene | 18794918 | 18795505 | FUN_026734 | -- |
| OZ035974.1 | gene | 18797013 | 18797717 | FUN_026735 | -- |
| OZ035974.1 | gene | 18800083 | 18810514 | FUN_026736 | -- |
| OZ035974.1 | gene | 18817354 | 18844659 | FUN_026739 | -- |
| OZ035974.1 | gene | 18823003 | 18823353 | FUN_026737 | -- |
| OZ035974.1 | gene | 18826311 | 18832935 | FUN_026738 | -- |
| OZ035974.1 | gene | 18858797 | 18859360 | FUN_026740 | -- |
| OZ035974.1 | gene | 18863174 | 18865739 | FUN_026741 | -- |
| OZ035974.1 | gene | 18872529 | 18898905 | FUN_026742 | -- |
| OZ035974.1 | gene | 18885819 | 18886255 | FUN_026743 | -- |
| OZ035974.1 | gene | 18901627 | 18901932 | FUN_026745 | -- |
| OZ035974.1 | gene | 18905268 | 18905567 | FUN_026746 | -- |
| OZ035974.1 | gene | 18906063 | 18908434 | FUN_026747 | -- |
| OZ035974.1 | gene | 18909712 | 18911143 | FUN_026748 | -- |
| OZ035974.1 | gene | 18912144 | 18925735 | FUN_026749 | -- |
| OZ035974.1 | gene | 18937223 | 18959043 | FUN_026750 | -- |
| OZ035974.1 | gene | 18962635 | 18966262 | FUN_026751 | -- |
| OZ035974.1 | gene | 18966514 | 18982410 | FUN_026752 | -- |
| OZ035974.1 | gene | 18988686 | 18989061 | FUN_026753 | -- |

**Table S11.4. Top candidate outlier loci as identified by PCAdapt.** 196 loci were identified to be potentially candidate outlier loci by PCAdapt. The top 1% of significant loci were extracted from that list. Table split into two (sets of) columns for aesthetic reasons only (reduce overall length); loci organized by descending p-values read left to right, top to bottom of list.

| chromo | position (bp) | p-value | | chromo | position (bp) | p-value |
|---|---|---|---|---|---|---|
| OZ035967.1 | 13856438 | 4.37E-115 | | OZ035967.1 | 13856772 | 4.37E-115 |
| OZ035967.1 | 13857787 | 4.37E-115 | | OZ035967.1 | 13858860 | 4.37E-115 |
| OZ035967.1 | 13861378 | 4.37E-115 | | OZ035967.1 | 13861962 | 4.37E-115 |
| OZ035967.1 | 13862717 | 4.37E-115 | | OZ035967.1 | 13862974 | 4.37E-115 |
| OZ035967.1 | 13863931 | 4.37E-115 | | OZ035967.1 | 13864547 | 4.37E-115 |
| OZ035967.1 | 13865346 | 4.37E-115 | | OZ035967.1 | 13865545 | 4.37E-115 |
| OZ035967.1 | 13868175 | 4.37E-115 | | OZ035967.1 | 13874745 | 4.37E-115 |
| OZ035967.1 | 13875848 | 4.37E-115 | | OZ035967.1 | 13879943 | 4.37E-115 |
| OZ035967.1 | 13880097 | 4.37E-115 | | OZ035967.1 | 13886651 | 4.37E-115 |
| OZ035967.1 | 13886893 | 4.37E-115 | | OZ035967.1 | 13887136 | 4.37E-115 |
| OZ035967.1 | 13888415 | 4.37E-115 | | OZ035967.1 | 13889688 | 4.37E-115 |
| OZ035967.1 | 13891033 | 4.37E-115 | | OZ035967.1 | 13896776 | 4.37E-115 |
| OZ035967.1 | 13898164 | 4.37E-115 | | OZ035967.1 | 13899414 | 4.37E-115 |
| OZ035967.1 | 13900099 | 4.37E-115 | | OZ035967.1 | 13907458 | 4.37E-115 |
| OZ035967.1 | 12824578 | 9.42E-81 | | OZ035967.1 | 15790917 | 1.21E-75 |
| OZ035968.1 | 23020605 | 1.21E-75 | | OZ035968.1 | 23021760 | 1.21E-75 |
| OZ035969.1 | 1591780 | 1.21E-75 | | OZ035969.1 | 1591900 | 1.21E-75 |
| OZ035972.1 | 9772791 | 1.21E-75 | | OZ035972.1 | 18757152 | 1.21E-75 |
| OZ035974.1 | 10372882 | 1.21E-75 | | OZ035975.1 | 12061783 | 1.21E-75 |
| OZ035978.1 | 5754502 | 1.21E-75 | | OZ035978.1 | 5971462 | 1.21E-75 |
| OZ035978.1 | 6377259 | 1.21E-75 | | OZ035978.1 | 6542288 | 1.21E-75 |
| OZ035978.1 | 6544010 | 1.21E-75 | | OZ035978.1 | 6559528 | 1.21E-75 |
| OZ035978.1 | 6601121 | 1.21E-75 | | OZ035978.1 | 6626887 | 1.21E-75 |
| OZ035978.1 | 6627357 | 1.21E-75 | | OZ035978.1 | 6632521 | 1.21E-75 |
| OZ035978.1 | 6636042 | 1.21E-75 | | OZ035978.1 | 6639074 | 1.21E-75 |
| OZ035978.1 | 6655887 | 1.21E-75 | | OZ035978.1 | 6961086 | 1.21E-75 |
| OZ035978.1 | 7034132 | 1.21E-75 | | OZ035978.1 | 7035962 | 1.21E-75 |
| OZ035978.1 | 7036846 | 1.21E-75 | | OZ035978.1 | 7037084 | 1.21E-75 |
| OZ035978.1 | 7037498 | 1.21E-75 | | OZ035978.1 | 7037800 | 1.21E-75 |
| OZ035978.1 | 7037934 | 1.21E-75 | | OZ035978.1 | 7037958 | 1.21E-75 |
| OZ035978.1 | 7038392 | 1.21E-75 | | OZ035978.1 | 7038861 | 1.21E-75 |
| OZ035978.1 | 7038922 | 1.21E-75 | | OZ035978.1 | 7039955 | 1.21E-75 |
| OZ035978.1 | 7040246 | 1.21E-75 | | OZ035978.1 | 7040823 | 1.21E-75 |
| OZ035978.1 | 7041911 | 1.21E-75 | | OZ035978.1 | 7043905 | 1.21E-75 |
| OZ035978.1 | 7312030 | 1.21E-75 | | OZ035978.1 | 7322182 | 1.21E-75 |
| OZ035978.1 | 7356285 | 1.21E-75 | | OZ035978.1 | 7410158 | 1.21E-75 |
| OZ035978.1 | 7550190 | 1.21E-75 | | OZ035978.1 | 8323877 | 1.21E-75 |
| OZ035979.1 | 8667681 | 1.21E-75 | | OZ035972.1 | 54395 | 4.16E-75 |
| OZ035967.1 | 13690425 | 2.22E-74 | | OZ035978.1 | 7038449 | 5.44E-74 |

| | | | | | | |
|---|---|---|---|---|---|---|
| OZ035967.1 | 13404949 | 7.62E-74 | | OZ035967.1 | 13658308 | 1.86E-72 |
| OZ035967.1 | 13595992 | 1.01E-67 | | OZ035969.1 | 10626344 | 4.25E-65 |
| OZ035967.1 | 13828250 | 4.73E-65 | | OZ035967.1 | 13785951 | 4.83E-62 |
| OZ035978.1 | 11023486 | 1.94E-61 | | OZ035978.1 | 7073083 | 3.57E-60 |
| OZ035967.1 | 13822146 | 5.20E-60 | | OZ035967.1 | 13156949 | 6.28E-59 |
| OZ035969.1 | 10254860 | 6.82E-58 | | OZ035969.1 | 10528812 | 6.82E-58 |

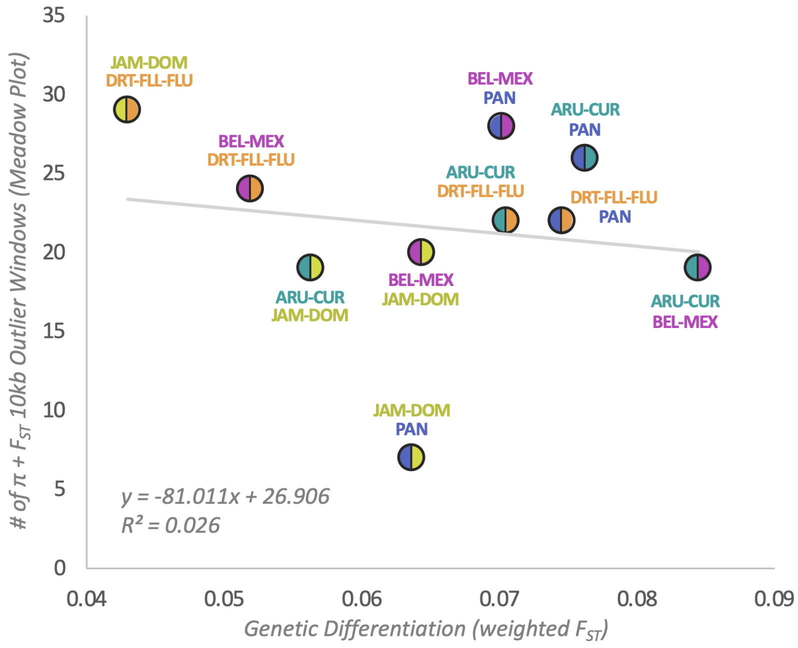

**Figure S11.5. Correlation exploring relationship between pairwise genetic differentiation** (weighted $F_{ST}$ on neutral panel, 177,516 SNPs) **and meadow plot outlier window counts** (see Figure S11.4, Table S11.2 for details) **among K=5 subpopulations**.

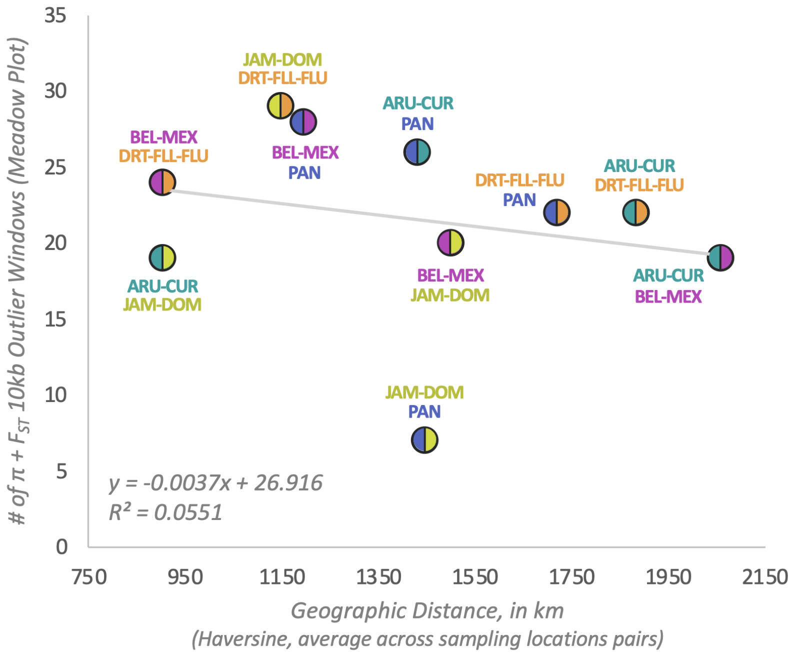

**Figure S11.6. Correlation exploring relationship between geographic distance** (km) **and meadow plot outlier window counts** (see Figure S11.4, Table S11.2 for details) **among K=5 subpopulations**.

### Appendix S12. Haplotype reference panel results.

**Table S12.1. Depth summary stats across BAMs used in haplotype reference panel construction**, calculated across retained chromosomes and across the 1,677,307 SNPs retained in the final filtered reference panel, with and without n=9 Panama ["PAN"] samples from Vollmer et al. (2023).

| | across retained chrs | | | | across ref pan SNPs | | | |
|---|---|---|---|---|---|---|---|---|
| for n=44 | average | std dev | max | min | average | std dev | max | min |
| all samples | 39.7182 | 8.63733 | 71.139 | 23.216 | 39.7182 | 8.63733 | 71.139 | 23.216 |
| exclud PAN | 37.4731 | 2.40737 | 41.9859 | 32.783 | 37.4731 | 2.40737 | 41.9859 | 32.783 |

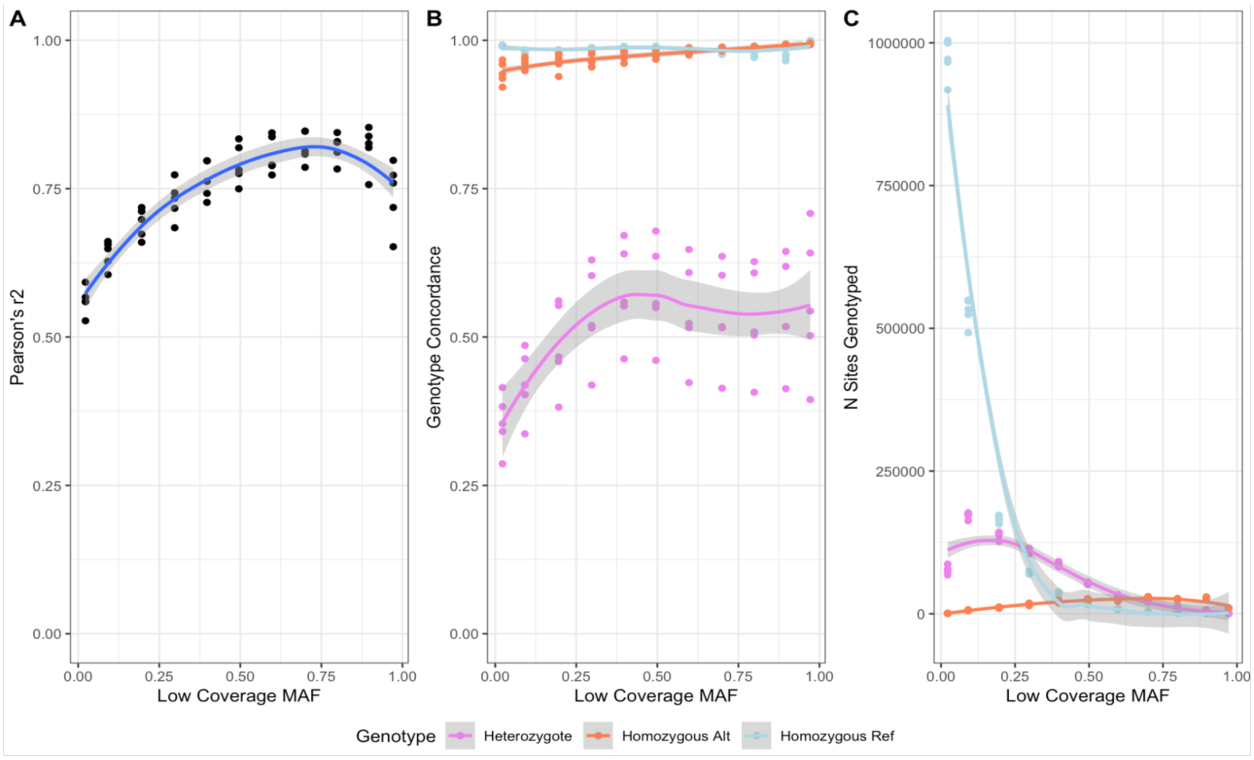

**Figure S12.1. Concordance of raw low coverage calls corresponding with "true" high coverage calls by genotypic class.**

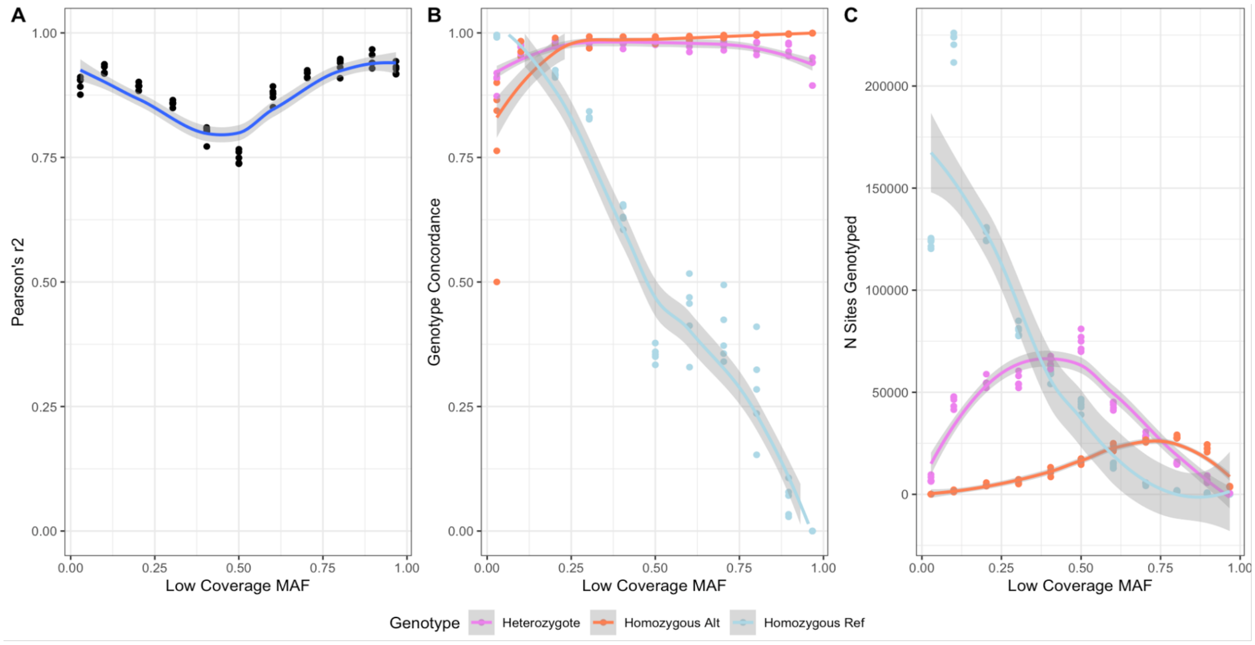

**Figure S12.2. Concordance of adjusted low coverage calls** corresponding with “true” high coverage calls by genotypic class, following Imputation Part I and removal of SNPs with low GP (retain GP > 0.99).

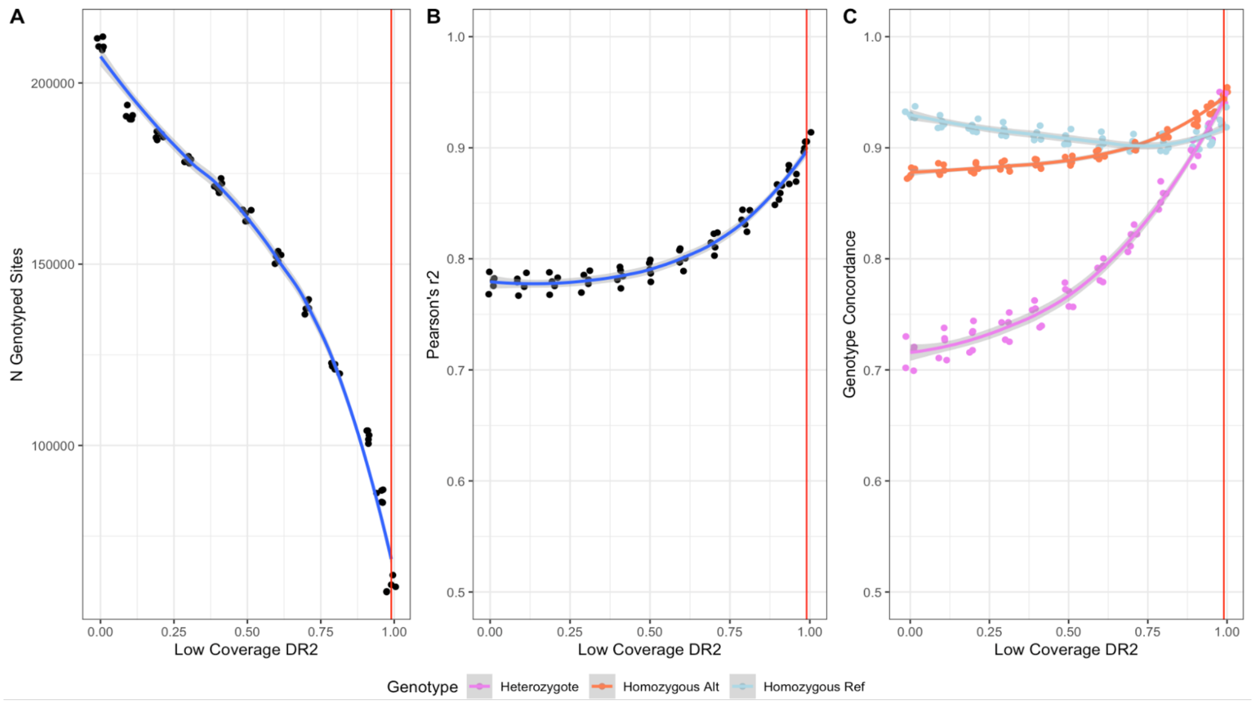

**Figure S12.3. Concordance of imputed sites** (“IMP” flag) against corresponding “true” high coverage calls by genotypic class, following Imputation Part II. Vertical red line indicates threshold chosen for DR2 filter (retain DR2 > 0.99).

### Appendix S13. Inbreeding and outbreeding depression risk evaluation.

**Appendix S13.1. Overview and rationale for the inbreeding and outbreeding depression risk evaluation.**

We used the decision-making framework proposed and implemented by (Liddell et al. 2021) to assess the risks of inbreeding and outbreeding depression for *A. cervicornis samples* represented in this study.

For inbreeding depression, we considered both observed genomic conditions and their expected near-term trajectory (marked with "Projected" notation in Figure S13.1). This distinction reflects the temporal lag between demographic decline and its genomic consequences: populations can retain relatively high diversity and low realized inbreeding for several generations after reductions in $N_e$ and connectivity, even as drift and mating among increasingly related individuals make future inbreeding more likely.

In contrast, we assessed outbreeding-depression risk (Figure S13.2) primarily from current and historical evidence because the signals informing that assessment are not expected to increase under the projected demographic trajectory. Although continued isolation may increase differentiation among subpopulations through genetic drift, drift-driven divergence does not itself indicate local adaptation or greater risk of outbreeding depression (Weeks et al. 2016). Moreover, local adaptation is generally strongest in large populations and, at least in plants, weak or absent in small populations, suggesting that declining and genetic diversity are more likely to constrain than promote the evolution of further adaptive divergence (Leimu & Fischer 2008; Weeks et al. 2016). Thus, near-term demographic decline is expected to increase inbreeding risk while providing little basis for projecting a corresponding increase in outbreeding-depression risk.

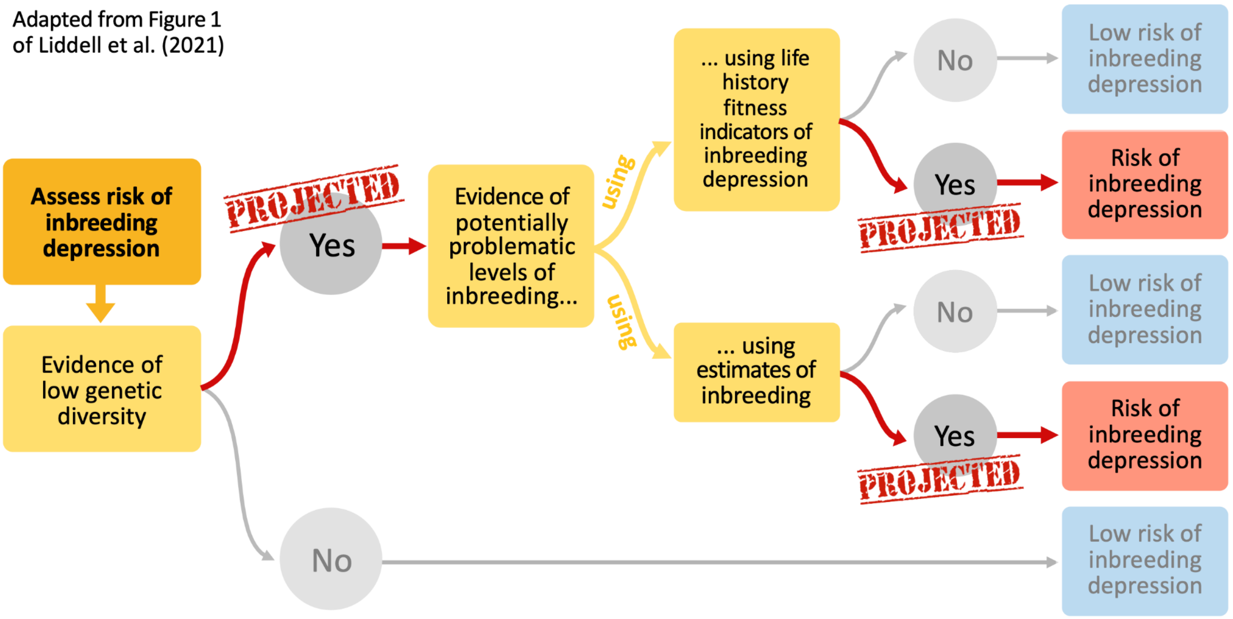

**Figure S13.1. Decision-making framework to assess the risks of inbreeding depression for *A. cervicornis* samples assessed in this study.** Adapted from Liddell et al. (2021) and assessed using both observed genomic conditions and their expected near-term trajectory.

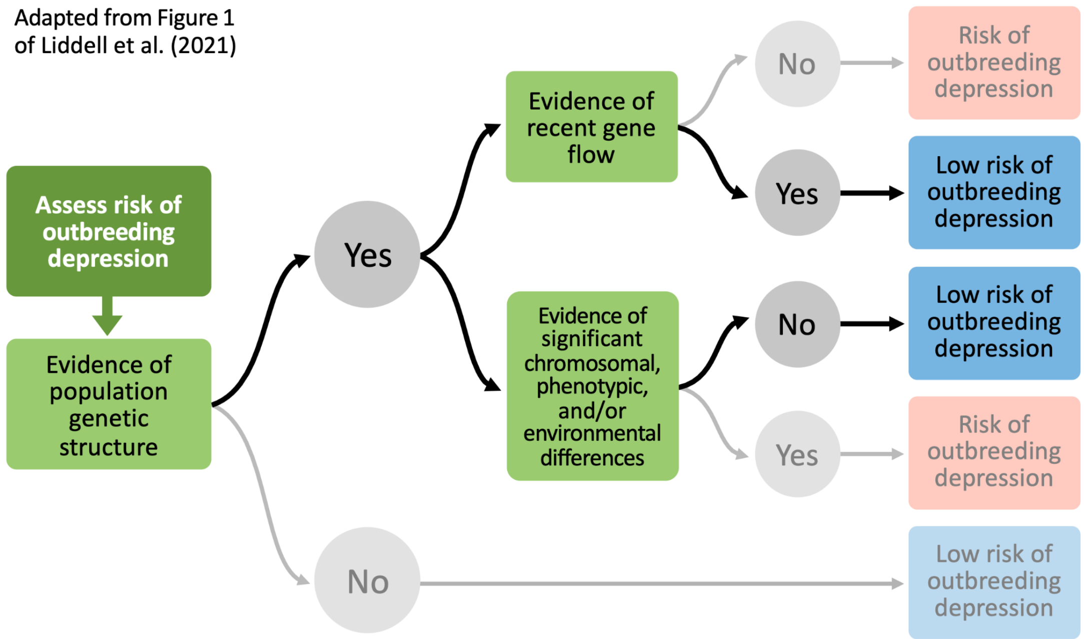

**Figure S13.2. Decision-making framework to assess the risks of outbreeding depression for *A. cervicornis* samples assessed in this study.** Adapted from Liddell et al. (2021).

#### **Appendix S14. Supplement references.**

Alexander DH, Novembre J, Lange K. 2009. Fast model-based estimation of ancestry in unrelated individuals. Genome Research **19**:1655–1664. Cold Spring Harbor Laboratory.

Blant A, Kwong M, Szpiech ZA, Pemberton TJ. 2017. Weighted likelihood inference of genomic autozygosity patterns in dense genotype data. BMC Genomics **18**:928. Springer Science and Business Media LLC.

Breusing C, Genetti M, Russell SL, Corbett-Detig RB, Beinart RA. 2022. Horizontal transmission enables flexible associations with locally adapted symbiont strains in deep-sea hydrothermal vent symbioses. Proceedings of the National Academy of Sciences of the United States of America **119**:e2115608119. Proceedings of the National Academy of Sciences.

Browning BL, Tian X, Zhou Y, Browning SR. 2021. Fast two-stage phasing of large-scale sequence data. American Journal of Human Genetics **108**:1880–1890. Elsevier BV.

Browning SR, Browning BL. 2007. Rapid and accurate haplotype phasing and missing-data inference for whole-genome association studies by use of localized haplotype clustering. American Journal of Human Genetics **81**:1084–1097. Elsevier BV.

Chang CC, Chow CC, Tellier LC, Vattikuti S, Purcell SM, Lee JJ. 2015. Second-generation PLINK: rising to the challenge of larger and richer datasets. GigaScience **4**:7. Oxford University Press (OUP).

Coleman RA, Weeks AR, Hoffmann AA. 2013. Balancing genetic uniqueness and genetic variation in determining conservation and translocation strategies: a comprehensive case study of threatened dwarf galaxias, Galaxiella pusilla (Mack) (Pisces: Galaxiidae). Molecular Ecology **22**:1820–1835. Wiley.

Cunning R, Baker AC. 2013. Excess algal symbionts increase the susceptibility of reef corals to bleaching. Nature Climate Change **3**:259–262. Springer Science and Business Media LLC.

Danecek P et al. 2021. Twelve years of SAMtools and BCFtools. GigaScience **10**:giab008. Oxford University Press (OUP).

Dixon P. 2003. VEGAN, a package of R functions for community ecology. Journal of Vegetation Science: Official Organ of the International Association for Vegetation Science **14**:927–930. Wiley.

Dray S, Dufour A-B. 2007. Theade4Package: Implementing the duality diagram for ecologists. Journal of Statistical Software **22**:1–20. Foundation for Open Access Statistic.

Duforet-Frebourg N, Bazin E, Blum MGB. 2014. Genome scans for detecting footprints of local adaptation using a Bayesian factor model. Molecular Biology and Evolution **31**:2483–2495. Oxford University Press (OUP).

Evanno G, Regnaut S, Goudet J. 2005. Detecting the number of clusters of individuals using the software STRUCTURE: a simulation study. Molecular Ecology **14**:2611–2620. Wiley.

Frichot E, François O. 2015. LEA: An R package for landscape and ecological association studies. Methods in Ecology and Evolution **6**:925–929. Wiley.

Frichot E, Mathieu F, Trouillon T, Bouchard G, François O. 2014. Fast and efficient estimation of individual ancestry coefficients. Genetics **196**:973–983. Oxford University Press (OUP).

Fuller ZL et al. 2020. Population genetics of the coral Acropora millepora: Toward genomic prediction of bleaching. Science (New York, N.Y.) **369**:eaba4674. American Association for the Advancement of Science (AAAS).

Garrison E, Marth G. 2012, July 17. Haplotype-based variant detection from short-read sequencing. Available from http://arxiv.org/abs/1207.3907.

Gaunt TR, Rodríguez S, Day IN. 2007. Cubic exact solutions for the estimation of pairwise haplotype frequencies: implications for linkage disequilibrium analyses and a web tool "CubeX." BMC Bioinformatics **8**:428. Springer Science and Business Media LLC.

Goudet J. 2005. hierfstat, a package for r to compute and test hierarchical *F*-statistics.

Molecular Ecology Notes **5**:184–186. Wiley.
Hijmans RJ, Williams E, Vennes C, Hijmans MRJ. 2017. Package ‘geosphere. Spherical trigonometry **1**:1–45.
Hume BCC, Smith EG, Ziegler M, Warrington HJM, Burt JA, LaJeunesse TC, Wiedenmann J, Voolstra CR. 2019. SymPortal: A novel analytical framework and platform for coral algal symbiont next-generation sequencing ITS2 profiling. Molecular Ecology Resources **19**:1063–1080. Wiley.
Jombart T, Devillard S, Balloux F. 2010. Discriminant analysis of principal components: a new method for the analysis of genetically structured populations. BMC Genetics **11**:94.
Kimura M, Ota T. 1969. The average number of generations until extinction of an individual mutant gene in a finite population. Genetics **63**:701–709. Oxford University Press (OUP).
Knaus BJ, Grünwald NJ. 2017. vcfr: a package to manipulate and visualize variant call format data in R. Molecular Ecology Resources **17**:44–53. Wiley.
Kolde R. 2025. Pretty heatmaps. cran.ms.unimelb.edu.au. Available from https://cran.ms.unimelb.edu.au/web/packages/pheatmap/pheatmap.pdf.
Kopelman NM, Mayzel J, Jakobsson M, Rosenberg NA, Mayrose I. 2015. Clumpak: a program for identifying clustering modes and packaging population structure inferences across K. Molecular Ecology Resources **15**:1179–1191. Wiley.
Korunes KL, Samuk K. 2021. pixy: Unbiased estimation of nucleotide diversity and divergence in the presence of missing data. Molecular Ecology Resources **21**:1359–1368. Wiley.
Krueger F. 2015. Trim Galore!: A wrapper around Cutadapt and FastQC to consistently apply adapter and quality trimming to FastQ files, with extra functionality for RRBS data. Babraham Institute. Available from https://cir.nii.ac.jp/crid/1370294643762929691 (accessed December 11, 2023).
Kwong AM et al. 2021. Robust, flexible, and scalable tests for Hardy-Weinberg equilibrium across diverse ancestries. Genetics **218**:iyab044. Oxford University Press (OUP).
Kyriazis CC, Robinson JA, Lohmueller KE. 2025. Long runs of homozygosity are reliable genomic markers of inbreeding depression. Trends in Ecology & Evolution **40**:874–884. Elsevier BV.
Leimu R, Fischer M. 2008. A meta-analysis of local adaptation in plants. PloS One **3**:e4010. Public Library of Science (PLoS).
Liddell E, Sunnucks P, Cook CN. 2021. To mix or not to mix gene pools for threatened species management? Few studies use genetic data to examine the risks of both actions, but failing to do so leads …. Biological Conservation. Elsevier. Available from https://www.sciencedirect.com/science/article/pii/S0006320721001245.
Li H. 2011. A statistical framework for SNP calling, mutation discovery, association mapping and population genetical parameter estimation from sequencing data. Bioinformatics (Oxford, England) **27**:2987–2993. Oxford University Press (OUP).
Li H. 2013, March 16. Aligning sequence reads, clone sequences and assembly contigs with BWA-MEM. Available from http://arxiv.org/abs/1303.3997.
Li H, Durbin R. 2011. Inference of human population history from individual whole-genome sequences. Nature **475**:493–496.
Locatelli N. 2024. Genomic insights into coral evolution and adaptation: A comparative study of Caribbean reef-builders. Pennsylvania State University.
Locatelli NS, Kitchen SA, Stankiewicz KH, Osborne CC, Dellaert Z, Elder H, Kamel B, Koch HR, Fogarty ND, Baums IB. 2024. Chromosome-level genome assemblies and genetic maps reveal heterochiasmy and macrosynteny in endangered Atlantic Acropora. BMC Genomics **25**:1119. Springer Science and Business Media LLC.
Luu K, Bazin E, Blum MGB. 2017. pcadapt: an R package to perform genome scans for selection based on principal component analysis. Molecular Ecology Resources **17**:67–77. Wiley.

Martin M, Patterson M, Garg S, O Fischer S, Pisanti N, Klau G, Schöenhuth A, Marschall T. 2016. WhatsHap: fast and accurate read-based phasing. bioRxiv. Available from http://dx.doi.org/10.1101/085050.
Matz MV, Treml EA, Aglyamova GV, Bay LK. 2018. Potential and limits for rapid genetic adaptation to warming in a Great Barrier Reef coral. PLoS Genetics **14**:e1007220.
Palacio-Castro AM, Dennison CE, Rosales SM, Baker AC. 2021. Variation in susceptibility among three Caribbean coral species and their algal symbionts indicates the threatened staghorn coral, Acropora cervicornis, is particularly susceptible to elevated nutrients and heat stress. Coral Reefs **40**:1601–1613. Springer Science and Business Media LLC.
Pemberton TJ, Absher D, Feldman MW, Myers RM, Rosenberg NA, Li JZ. 2012. Genomic patterns of homozygosity in worldwide human populations. American Journal of Human Genetics **91**:275–292. Elsevier BV.
Pembleton LW, Cogan NOI, Forster JW. 2013. StAMPP: an R package for calculation of genetic differentiation and structure of mixed-ploidy level populations. Molecular Ecology Resources **13**:946–952. Wiley.
Petkova D, Novembre J, Stephens M. 2016. Visualizing spatial population structure with estimated effective migration surfaces. Nature Genetics **48**:94–100. Springer Science and Business Media LLC.
Picard. (n.d.). Available from http://broadinstitute.github.io/picard (accessed June 9, 2026).
Privé F, Luu K, Vilhjálmsson BJ, Blum MGB. 2020. Performing highly efficient genome scans for local adaptation with R package pcadapt version 4. Molecular Biology and Evolution **37**:2153–2154. Oxford University Press (OUP).
Reich HG, Kitchen SA, Stankiewicz KH, Devlin-Durante M, Fogarty ND, Baums IB. 2021. Genomic variation of an endosymbiotic dinoflagellate (Symbiodinium “fitti”) among closely related coral hosts. Molecular Ecology **30**:3500–3514. Wiley.
Sadler AJ, Williams BRG. 2008. Interferon-inducible antiviral effectors. Nature Reviews. Immunology **8**:559–568. Springer Science and Business Media LLC.
Szpiech ZA, Blant A, Pemberton TJ. 2017. GARLIC: Genomic autozygosity regions likelihood-based inference and classification. Bioinformatics (Oxford, England) **33**:2059–2062. Oxford University Press (OUP).
Szpiech ZA, Xu J, Pemberton TJ, Peng W, Zöllner S, Rosenberg NA, Li JZ. 2013. Long runs of homozygosity are enriched for deleterious variation. American Journal of Human Genetics **93**:90–102. Elsevier BV.
Terhorst J, Kamm JA, Song YS. 2017. Robust and scalable inference of population history from hundreds of unphased whole genomes. Nature Genetics **49**:303–309. Springer Science and Business Media LLC.
Vollmer SV, Selwyn JD, Despard BA, Roesel CL. 2023. Genomic signatures of disease resistance in endangered staghorn corals. Science (New York, N.Y.) **381**:1451–1454.
Wang J. 2025. PopCluster: A population genetics model-based toolset for simulating, inferring and visualising individual admixture and population structure. Molecular Ecology Resources **25**:e14058. Wiley.
Weeks AR, Stoklosa J, Hoffmann AA. 2016. Conservation of genetic uniqueness of populations may increase extinction likelihood of endangered species: the case of Australian mammals. Frontiers in Zoology **13**:31. Springer Science and Business Media LLC.
Weir BS, Cockerham CC. 1984. Estimating F-statistics for the analysis of population structure. Evolution; International Journal of Organic Evolution **38**:1358. Oxford University Press (OUP).
Whitlock MC, Lotterhos KE. 2015. Reliable Detection of Loci Responsible for Local Adaptation: Inference of a Null Model through Trimming the Distribution of FST. The American Naturalist **186**:S24–S36.
Zhang E, Khanna V, Dacheux E, Namane A, Doyen A, Gomard M, Turcotte B, Jacquier A,

Fromont-Racine M. 2019. A specialised SKI complex assists the cytoplasmic RNA exosome in the absence of direct association with ribosomes. The EMBO Journal **38**:e100640. Springer Science and Business Media LLC.

**<u>In press:</u>**

Duffin PJ, Schiebelhut LM, Dawson MN, Wares JP. In press. Genomic separation of Salish Sea and Pacific outer coast populations of the keystone sea star *Pisaster ochraceus*. Evolution.
